# Castration-mediated IL-8 Promotes Myeloid Infiltration and Prostate Cancer Progression

**DOI:** 10.1101/651083

**Authors:** Zoila A. Lopez-Bujanda, Michael C. Haffner, Matthew G. Chaimowitz, Nivedita Chowdhury, Nicholas J. Venturini, Aleksandar Obradovic, Corey S. Hansen, Joanna Jacków, Karen S. Sfanos, Charles J. Bieberich, Paula J. Hurley, Mark J. Selby, Alan J. Korman, Angela M. Christiano, Angelo M. De Marzo, Charles G. Drake

**Author notes:** Correspondence and request for materials should be addressed to C.G.D.

## Abstract

Immunotherapy is a treatment for many types of cancer, primarily due to deep and durable clinical responses mediated by immune checkpoint blockade (ICB)^1, 2^. Prostate cancer is a notable exception in that it is generally unresponsive to ICB. The standard treatment for advanced prostate cancer is androgen-deprivation therapy (ADT), a form of castration (CTX). ADT is initially effective, but over time patients eventually develop castration-resistant prostate cancer (CRPC). Here, we focused on defining tumor-cell intrinsic factors that contribute to prostate cancer progression and resistance to immunotherapy. We analyzed cancer cells isolated from castration-sensitive and castration-resistant prostate tumors, and discovered that castration resulted in significant secretion of Interleukin-8 (IL-8) and it’s likely murine homolog Cxcl15. These chemokines drove subsequent intra-tumoral infiltration with polymorphonuclear myeloid-derived suppressor cells (PMN-MDSCs), promoting tumor progression. PMN-MDSC infiltration was abrogated when IL-8 was deleted from prostate cancer epithelial cells using CRISPR/Cas9, or when PMN-MDSC migration was blocked with antibodies against the IL-8 receptor CXCR2. Blocking PMN-MDSC infiltration in combination with anti-CTLA-4 delayed the onset of castration resistance and increased the density of polyfunctional CD8 T cells in tumors. Taken together, our findings establish castration-mediated IL-8 secretion and subsequent PMN-MDSC infiltration as a key suppressive mechanism in the progression of prostate cancer. Targeting of the IL-8/CXCR2 axis around the time of ADT, in combination with ICB, represents a novel therapeutic approach to delay prostate cancer progression to advanced disease.

## Main

After primary therapy with surgery or radiation, approximately 40% of prostate cancer patients develop progressive disease. The standard treatment for recurrent prostate cancer is androgen-deprivation therapy (ADT), but the majority of these patients eventually develop castration-resistance (CR). Although some patients with metastatic castration-resistant prostate cancer (mCRPC) benefit from the cancer vaccine sipuleucel-T^3^, neither CTLA-4 blockade^4, 5^ nor PD-1 blockade^6^ has reliably produced meaningful clinical responses. Potential reasons for this include a low total mutational burden (TMB) as well as poor infiltration by CD8 T cells^7^.

We and others have shown that ADT initially increases CD8 T cell infiltration into prostate tumors^8–10^, and this response is augmented pre-clinically with anti-CTLA-4^11^. Emerging data suggest that immune-resistance in prostate cancer involves dysfunctional myeloid cells known as myeloid-derived suppressor cells (MDSCs) in the tumor microenvironment (TME)^12, 13^. MDSCs secrete IL-23, which acts directly on prostate cancer epithelial cells to drive castration-resistance^14^. Importantly, the mechanism(s) by which suppressive MDSCs are recruited to the prostate TME are largely unknown.

To identify immune-related tumor-cell intrinsic factors involved in prostate cancer progression, we performed expression analyses on murine prostate cancer cells pre- and post-castration. We used the MCRedAL prostate cancer cell line; an RFP expressing version of the Myc-Cap cell line characterized by *MYC* overexpression^15^. Like human prostate cancer, MCRedAL tumors are initially castration-sensitive (CS), but castration-resistance (CR) develops approximately 30 days after castration (Extended Data Fig. 1a). Pre- and post-ADT tumor cells were sorted to > 96% purity (Extended Data Fig. 1b) and analyzed (Fig. 1a-b and Extended Data Fig. 1c). A number of cytokine and chemokine transcripts were significantly up-regulated post-ADT (Fig. 1b right), including *Cxcl15*, a CXC chemokine with a conserved ELR motif (Extended Data Table 1), which is the likely murine homolog of human *IL-8* (*CXCL8*)^16–19^. qRT-PCR and ELISA assays confirmed the upregulation of Cxcl15 post-ADT at the protein level (Extended Data Fig. 1d). In addition to the chemokines above, GSEA revealed the upregulation of several pro-inflammatory pathways post-ADT (Fig. 1c). *In vitro* experiments using the human androgen-responsive LNCaP cell line corroborated a role for these pro-inflammatory signals, showing that in the absence of androgen, TNFα upregulated IL-8 expression in a dose-dependent manner (Fig. 1d left); while AR signaling in the absence of inflammation did not affect IL-8 expression (Fig. 1d right). These data led to the hypothesis that AR signaling directly suppresses *IL-8* expression in prostate cancer cells. We performed *in silico* ChIP-Seq analyses using human LNCaP cells (GSE83860) and found AR binding at the *IL-8* promoter in the presence of the potent androgen dihydrotestosterone (DHT; Fig. 1e top). This androgen dependent binding was verified by ChIP-qRT-PCR (Fig. 1f).

**Figure 1.**
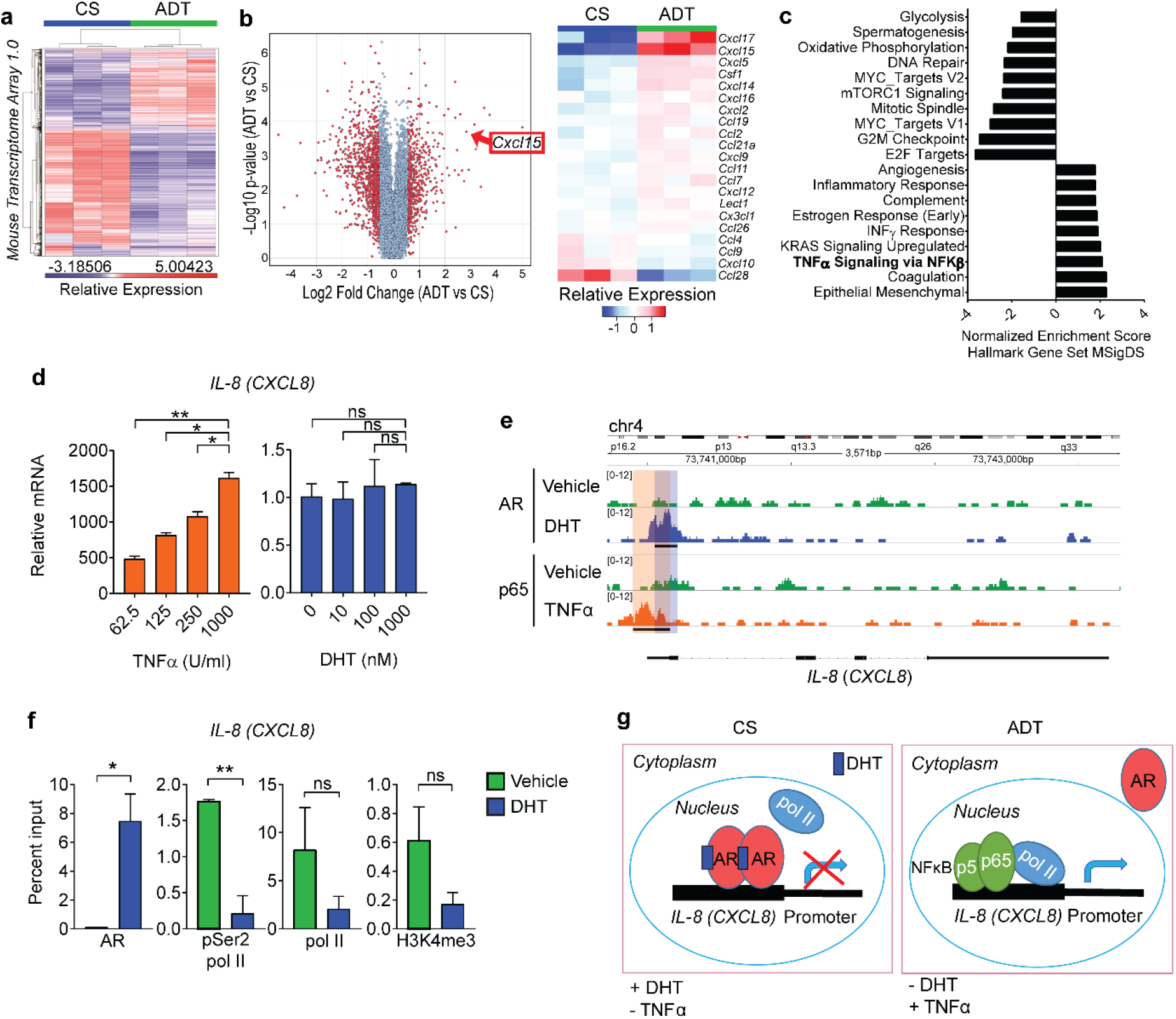
Androgen-Deprivation Therapy (ADT) Increases IL-8 Expression in Prostate Cancer Cells. **a,** Differential expression profile of tumor epithelial cells isolated from castration-sensitive (CS) and ADT-treated MCRedAL tumor bearing mice. Heatmap showing transcripts 3 standard deviations away from the mean (n = 3 per group). **b,** Differential chemokine expression of tumor epithelial cells isolated from CS and ADT tumor bearing mice (replicate numbers as in **a**). Left, volcano plot showing gene expression among all MTA 1.0 microarray transcripts. Right, heatmap of normalized chemokine transcripts. **c,** Hallmarks gene sets pathway analysis post-ADT shows NF-κB up-regulation post-ADT. **d,** qRT-PCR quantification of *IL-8* in LNCaP cells cultured at indicated concentrations of TNFα and DHT, cells cultured in androgen-free media as described in materials and methods (n = 3 per condition, repeated × 2). Expression levels normalized to mean ΔCT level in samples cultured in androgen free media without TNFα or DHT. **e,** ChIP-Seq analysis of AR at the *IL-8* (*CXCL8*) promoter in LNCaP cells cultured in the presence of either vehicle (DMSO), DHT (100 nM), or TNFα (1000 U/ml) (n = 2 per group; GSE83860). **f,** ChIP quantitative RT-PCR (qRT-PCR) analysis of AR, pSer2 Pol II, pol II, and H3K4me3 at the *IL-8* (*CXCL8*) promoter (n = 3 per group). Transfected LNCaP cells treated for 24 hours with or without DHT (100 nM). **g,** Schematic model of the interplay between AR and NFkB in the regulation of IL-8 transcription. For **e**, loci with significant differential binding (black bar) were identified as described in materials and methods. Error bars represent standard error. Unpaired t-tests were performed, *p*-values ≤ 0.05 (*), 0.01 (**), 0.001 (***) and 0.0001 (****); *p*-values ≥ 0.05 (ns).

To further explore the role of AR in IL-8 regulation, we interrogated RNA polymerase binding and transcription marks found at sites of active promoters^20^. In the presence of DHT, binding of RNA polymerase II (pol II), phosphorylated serine 2 RNA polymerase II (pSer2 pol II) and histone H3 tri-methyl Lys4 (H3K4me3) to the IL-8 locus were substantially reduced, consistent with reduced transcriptional activity (Fig. 1f). Conversely, pSer2 pol II binding to the promoter of the well-established AR-regulated gene *PSA* (*KLK3*), was significantly increased in the presence of DHT as expected (Extended Data Fig. 1e). Consistent with a role for inflammation, TNFα significantly increased p65 binding at the *IL-8* (*CXCL8*) promoter in LNCaP cells (Fig. 1e bottom). No significant binding of AR was detected at the promoters of the chemokines *CXCL1*, *CXCL2*, *CXCL5* or *CXCL12* (Extended Data Fig. 1f). These data suggest that AR directly suppresses *IL-8* expression through repressive AR binding to the *IL-8* promoter. Taken together, we found that IL-8 transcription is up-regulated by pro-inflammatory signaling, and down-regulated by AR signaling (Fig. 1g).

We next investigated the effects of ADT on the expression of *Cxcl15 in vivo*, using RNA in situ hybridization (RISH) to study Myc-Cap tumors. We found that CR tumors expressed increased *Cxcl15* as compared to CS tumors, particularly in epithelial (PanCK^+^) tumor cells (Fig. 2a, Extended Data Fig. 2a). These findings were confirmed *in vitro*, both at the mRNA and protein level (Fig. 2b). To investigate these findings in the context of human prostate cancer, we used three paired cell lines in which isogenic CR lines were derived from CS progenitors. For each pair, the CR line expressed significantly increased IL-8 as compared to the CS counterpart, both at the mRNA and protein level (Fig. 2c-d). This observation held across a panel of AR expressing prostate cancer cell lines; with higher levels of *IL-8* expression in cell lines from castration-resistant disease (Extended Data Fig. 2b). To test whether AR modulates *Cxcl15* expression in <u>benign</u> prostate epithelium, we used RISH to study WT mice treated with ADT, and WT mice treated with ADT followed by testosterone (T) repletion (Extended Data Fig. 2c). These data (Fig. 2e-f) showed increased epithelial *Cxcl15* expression in ADT samples with expression significantly decreased by testosterone repletion (Fig. 2f). This observation was further corroborated by interrogating a dataset (GSE8466) profiling human prostate epithelial cells isolated by laser-capture microdissection (LCM) from men undergoing ADT and ADT with testosterone supplementation. Testosterone repletion significantly reduced *IL-8* mRNA expression (Fig. 2g), supporting the hypothesis that AR signaling down-regulates IL-8 expression. In agreement with these data from benign prostate tissues, we LCM-enriched tumor prostate epithelium from high-risk PCa patients treated with ADT on a neo-adjuvant trial (NCT01696877) and found increased *IL-8* expression as compared to tumors from age and stage-matched untreated controls (Fig. 2h). Taken together, analyses using human tissues strongly support the notion that castration increases *IL-8* expression in prostate epithelial cells.

**Figure 2.**
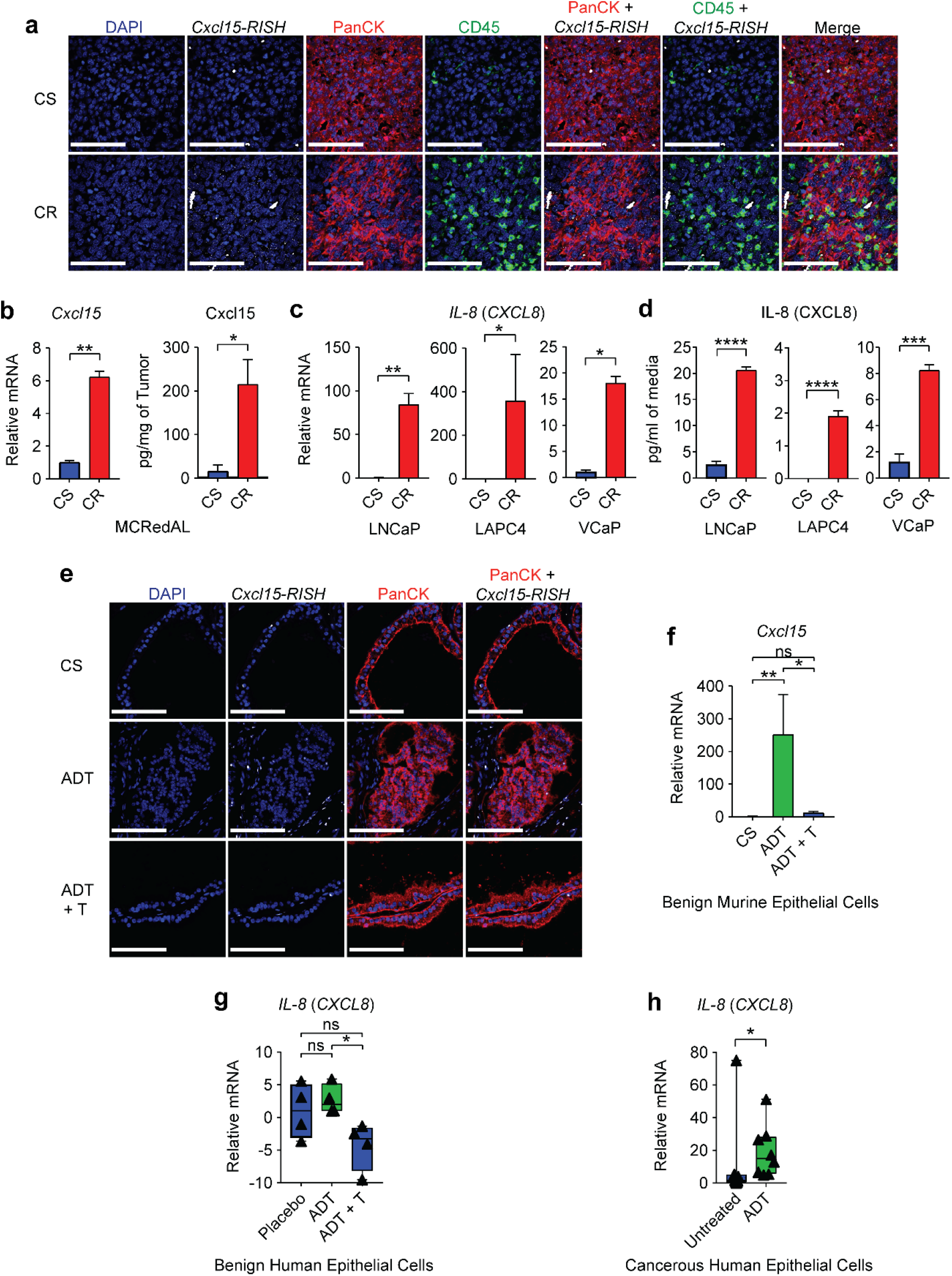
IL-8 is Up-Regulated in Post-Castration and Castration-Resistant Prostate Cancer Cells. **a,** Representative images of *Cxcl15* fluorescent detection (murine homologue of IL-8) in Myc-Cap tumors. Tumors were harvested when volumes reached ∼500mm^3^ (CS group), 7 days after androgen-deprivation (ADT), or at the time of castration-resistance (CR) and hybridized with CF568-labeled probe sets (white) to *Cxcl15*, CF640-labeled anti-PanCK antibody (red), and CF488-labeled anti-CD45 antibody (green). Nuclei counterstained with DAPI (blue). Repeated × 3. **b,** Gene and protein expression of Cxcl15 in MCRedAL cells of indicated tumor samples by qRT-PCR and ELISA, respectively (n = 3 per group, repeated × 2). **c,** qRT-PCR quantification of *IL-8* in human AR positive castration-sensitive cells (CS: LNCaP, LAPC4, and VCaP) and their castration-resistant counterparts (CR: LNCaP-abl, LAPC4-CR, and VCaP-CR), replicate numbers as in **b**. **d,** IL-8 protein expression in the isogenic cell pairs from **c** quantified by ELISA, replicate numbers as in **c**. **e,** Representative images of *Cxcl15* fluorescent detection in benign murine prostate tissue samples from castration-sensitive (CS), androgen-deprivation treated (ADT), and ADT-treated mice that received testosterone repletion (ADT + T). Tissue sections hybridized with CF568-labeled probe sets (white) to *Cxcl15*, and CF640-labeled anti-PanCK antibody (red). Nuclei were counterstained with DAPI (blue). Repeated × 3. **f,** qRT-PCR analysis of *Cxcl15* expression in prostate luminal epithelial cells from indicated treatment groups (n = 3 per group). Prostate luminal epithelial cells were isolated based on their GFP^+^CD49f^int^CD24^+^CD45^-^F4/80^-^CD11b^-^ expression by flow sorting into Trizol LS. **g,** Expression of *IL-8* in human prostate epithelial cells micro-dissected from patients in a clinical trial (NCT00161486) receiving placebo, androgen-deprivation treatment (ADT), or ADT plus testosterone repletion (ADT + T). Z-score values of microarray transcripts from benign prostate biopsies were normalized to placebo samples (n = 4 per group; GSE8466). **h,** Expression of *IL-8* in human prostate cancer epithelial cells micro-dissected from untreated or ADT-treated (NCT01696877; n = 8 per group) patients as determined by qRT-PCR. RISH images are at 60X magnification; scale bar = 100 μm. Gene expression levels were normalized to the mean ΔCT level in samples from CS, untreated or placebo groups. For **b-g**, unpaired t-tests were performed; for **h** a Mann-Whitney U test was used due to the non-normal data distribution observed. *p*-values ≤ 0.05 (*) and 0.01 (**); *p*-values ≥ 0.05 (ns) shown. The range in box and whiskers plots shows min and max values such that all data are included.

We next quantified castration-mediated immune infiltration in Myc-Cap allografts (Fig. 3a). Consistent with prior data^11^, ADT promoted a transient T cell influx, without significant changes in tumor associated macrophage (TAM) populations (Fig. 3b). By contrast, PMN-MDSC infiltration was significantly increased in CR tumors (Fig. 3b), as verified by IHC (Fig. 3c). We found similar results in human prostate cancer xenografts (Extended Data Fig. 3a-b). PMN-MDSC infiltration also increased in WT mice treated with ADT, but not in WT mice treated with ADT then repleted with testosterone (Extended Data Fig. 3c), supporting a causal relationship between ADT and PMN-MDSC infiltration. Molecular profiling of the infiltrating myeloid cells revealed a signature consistent with functional PMN-MDSCs, including up-regulation of *IL-1b*, *Arg2* and *IL-23a*^14^ (Fig. 3d; Extended Data Table 2). In particular, increased expression of *IL-23a* and *Cxcr2* was verified by qRT-PCR (Fig. 3e) and flow cytometry (Extended Data Fig. 3d). To test whether blocking the IL-8/CXCR2 axis was sufficient to attenuate post-ADT PMN-MDSC infiltration, we treated prostate-tumor bearing mice with anti-CXCR2 and found that blocking CXCR2 significantly diminished tumor infiltration with PMN-MDSCs in both human (PC3) and murine (Myc-Cap) immunodeficient and immunocompetent models (Fig. 3f and Extended Data Fig. 3e-f). To confirm this observation at the genetic level, we used CRISPR/Cas9 to generate human (PC3) and mouse (Myc-Cap) lines that were knocked out for human IL-8 or the murine IL-8 homolog Cxcl15, respectively. We observed a clear decrease in PMN-MDSC infiltration in both settings (Fig. 3g and Extended Data Fig. 3e-f).

**Figure 3.**
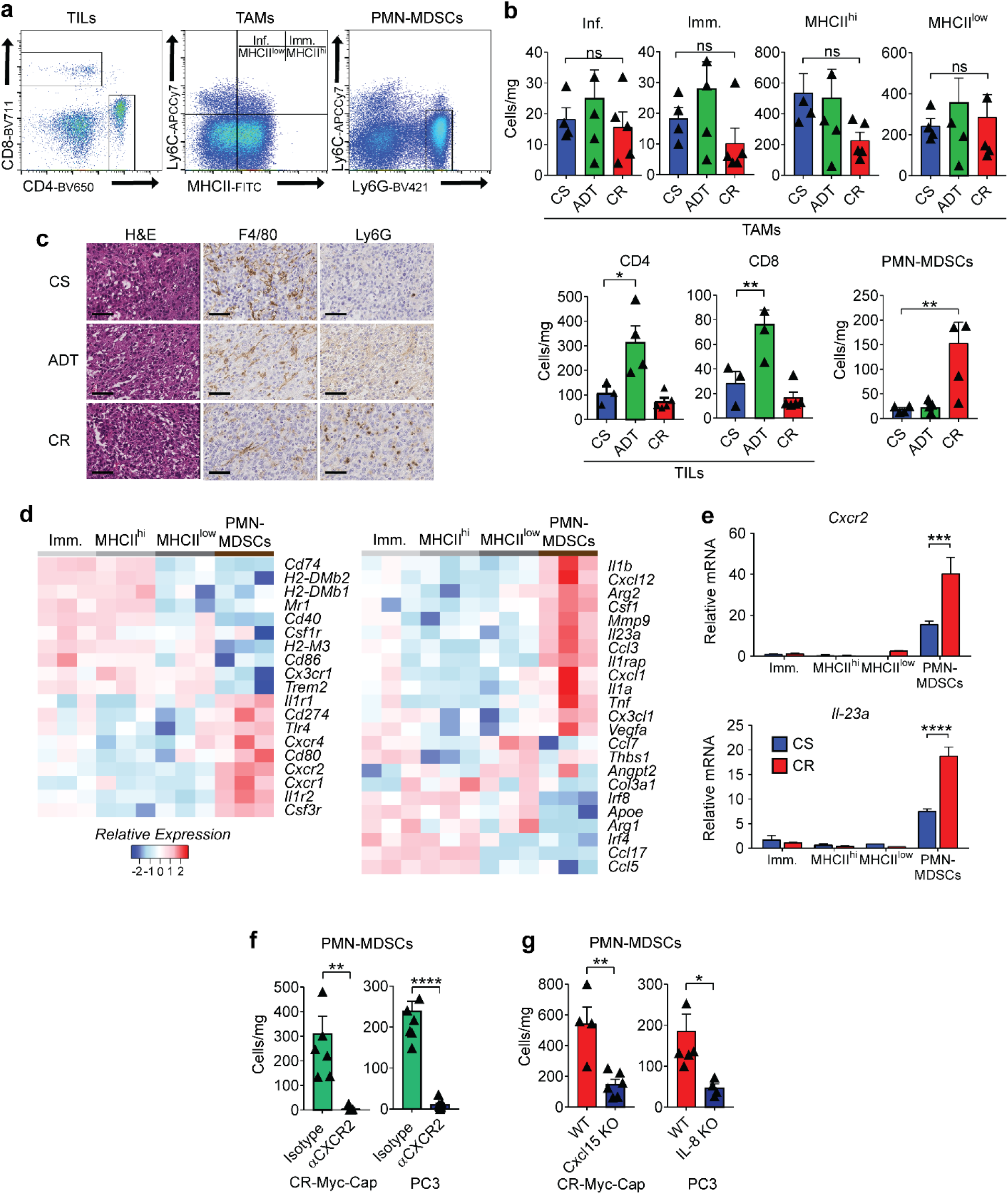
Castration-mediated IL-8 Up-Regulation Promotes PMN-MDSC Infiltration. **a,** Gating strategy used to profile the immune compartment of the TME by flow cytometry. Tumor associated macrophages (TAMs) gated based on CD45^+^Ly6G^-^ F4/80^+^CD11b^+^, Inflammatory (Inf.) TAMs as CD45^+^CD11b^+^F4/80^+^Ly6C^+^MHCII^-^, immature (Imm.) TAMs as CD45^+^CD11b^+^F4/80^+^Ly6C^+^MHCII^+^, MHCII^hi^ TAMs as CD45^+^CD11b^+^F4/80^+^Ly6C^-^MHCII^+^, MHCII^low^ TAMs as CD45^+^CD11b^+^F4/80^+^Ly6C^-^MHCII^-^, tumor Infiltrating Lymphocytes (TILs) CD45^+^CD4^+^ or CD45^+^CD8^+^, tumor infiltrating polymorphonuclear myeloid-derived suppressor cells (PMN-MDSCs) as CD45^+^CD11b^+^Ly6C^+^Ly6G^+^. **b,** TAM, TIL, and PMN-MDSC density normalized to mg of tumor weight (cells/mg; n ≥ 3 per group, repeated × 2). **c,** Representative H&E and immunohistochemistry (F4/80 and Ly6G) of indicated murine allografts (repeated × 3). **d,** Normalized expression of selected genes determined by NanoString nCounter gene analysis in sorted myeloid fractions defined as in **a** (n = 3 per group). **e,** qRT-PCR quantification of *Cxcr2* and *Il-23* in indicated populations of Myc-Cap tumors (n = 3 per group). **f** and **g,** Density of PMN-MDSCs normalized to mg of tumor weight (cells/mg) in Myc-Cap and PC3 tumors (n ≥ 4 per group, repeated × 2). Cells quantified by flow cytometry as in **a**, tumors implanted and harvested as in materials and methods. H&E and IHC images at 40X magnification; scale bar = 50 μm. Gene expression levels normalized to the mean ΔCT level in samples from the Immature TAMs (Imm.) group. Unpaired t-tests performed, *p*-values ≤ 0.05 (*), 0.01 (**), 0.001 (***) and 0.0001 (****); *p*-values ≥ 0.05 (ns).

We next asked whether the supernatants from castration-resistant MCRedAL (CR-MCRedAL) cells were sufficient to drive PMN-MDSC migration *in vitro*. In line with *in vivo* results (Fig. 3f-g and Extended Data Fig. 4a-c), we found that PMN-MDSC migrated towards the supernatant of CR tumors and migration was significantly attenuated by CXCR2 blockade (Extended Data Fig. 4d). Human prostate cancer (PC3) showed an identical pattern. To confirm a role for IL-8 in PMN-MDSC migration, we generated IL-8 KO CR-LNCaP (LNCaP-abl) using CRISPR/Cas9. Supernatants from IL-8 KO cells were significantly attenuated in their ability to promote PMN-MDSC migration (Extended Data Fig. 4e). These PMN-MDSCs were functional and suppressed CD8 T cell proliferation in a dose-dependent manner (Extended Data Fig. 4f-i). Although CXCR2 blockade decreased PMN-MDSC migration, it did not significantly alter their suppressor function (Extended Data Fig. 4j). Similarly, Cxcl15 loss did not diminish the suppressive function of PMN-MDSCs (Extended Data Fig. 4k). Taken together these findings reinforce a functional role for castration-mediated IL-8 secretion in PMN-MDSC migration.

Finally, we investigated the pre-clinical activity of blocking the IL-8/CXCR2 axis at the time of androgen-deprivation in the Myc-Cap model. Notably, in the absence of immunotherapy the combination of ADT and CXCR2 blockade was not effective (Extended Data Fig. 5a). In contrast, combining CXCR2 blockade with ICB (anti-CTLA-4; Fig. 4a) resulted in significantly increased survival (Fig. 4b). This triple combination (ADT + anti-CXCR2 + anti-CTLA-4) was effective even when tumors were relatively advanced (400 mm^3^) at the time of treatment (Extended Data Fig. 5b&d). Macrophage modulation with anti-CSF1R was not effective therapeutically in this setting (Extended Data Fig. 5c&e). Mechanistically, the increased anti-tumor effects mediated by the addition of anti-CXCR2 to ADT + anti-CTLA-4 did not appear to be due to increased T cell infiltration (Fig. 4c and Extended Data Fig. 5f-h), nor due to decreased Treg infiltration (Fig. 4d), but rather correlated with an increase in polyfunctional effector CD8 T cells in tumor-draining lymph nodes (TDLN) and spleens (Fig. 4e&f).

**Figure 4.**
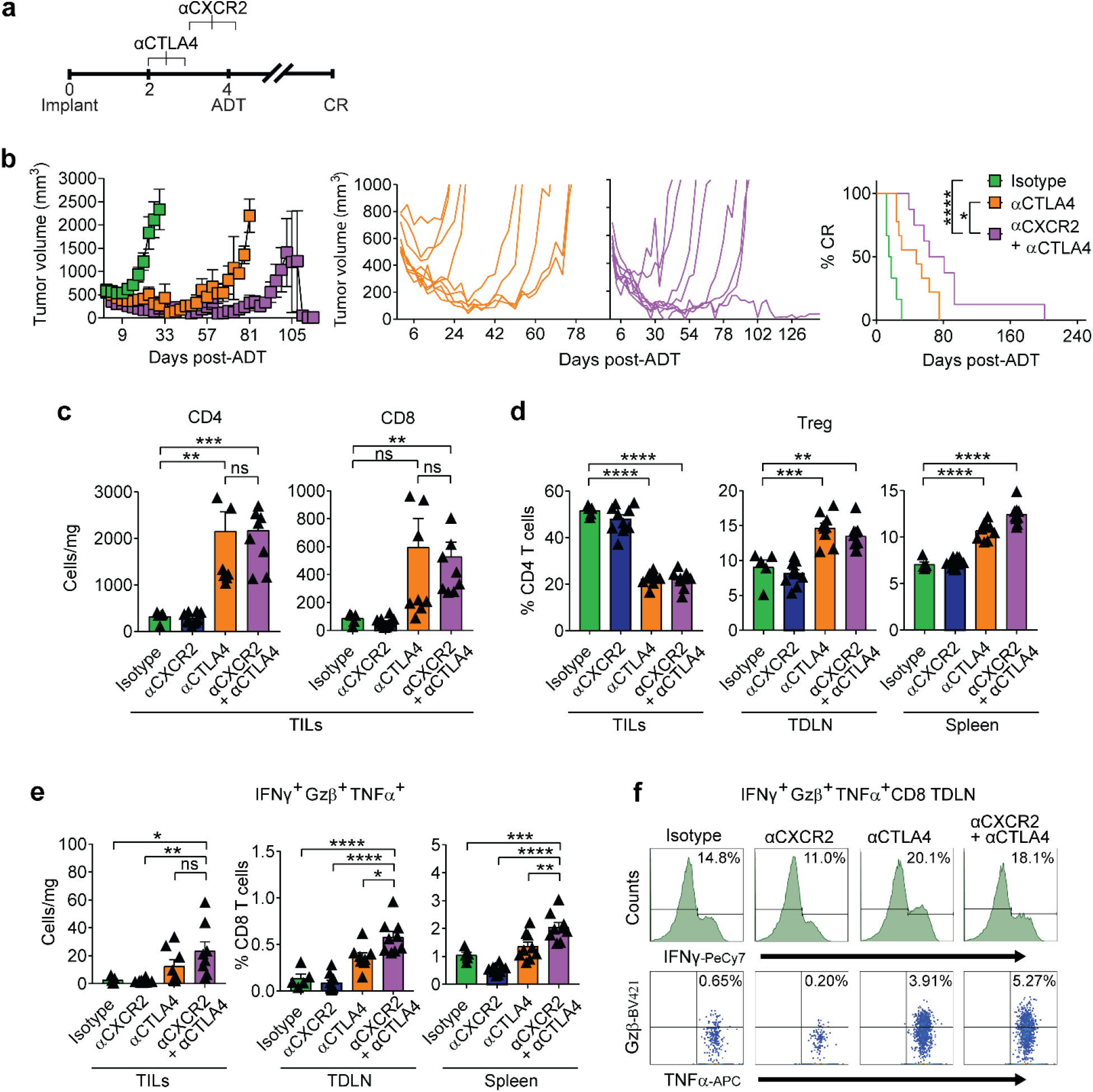
CXCR2 Blockade Improves Response to Immune Checkpoint Blockade Following Androgen-Deprivation Therapy. **a,** Treatment scheme, scale = weeks. Animals sacrificed for immune phenotyping 1 week post-ADT. **b,** Tumor growth and survival curves of mice from isotype vs. anti-CTLA-4 vs. anti-CTLA-4 + anti-CXCR2 groups treated as described in **a** (black line vs. orange line vs. purple line, respectively; n ≥ 8 per group, repeated × 2). **c,** Tumor infiltrating lymphocyte (TILs) density in indicated treatment groups (n ≥ 5 per group, repeated × 2). **d,** Treg percentages (as fraction of CD4) in indicated tissues (n ≥ 5 per group, repeated × 2). **e,** Polyfunctional CD8 T cells, left panel = density, center/right panels = percentage of total CD8, animals numbers as in **d**. **f,** Representative histograms and dot plots of polyfunctional CD8^+^ IFNγ^+^Gzβ^+^TNFα^+^ from tumor draining lymph nodes (TDLN). Repeated × 2. For **a-f**, treatment was initiated when tumor volumes reached 200mm^3^. Average tumor volume (±s.e.m.) for each experimental group. Wilcoxon test used for survival analysis. Flow cytometry as in materials and methods. Unpaired t-tests performed, *p*-values ≤ 0.05 (*), 0.01 (**), 0.001 (***) and 0.0001 (****); *p*-values ≥ 0.05 (ns).

In summary, these studies showed that castration mediates increased IL-8 secretion by prostate cancer epithelial cells by releasing AR-mediated transcriptional repression. IL-8 (and Cxcl15) up-regulation then drives prostate tumor infiltration with PMN-MDSCs. We found that blocking CXCR2 at the time of androgen-deprivation therapy attenuates PMN-MDSC infiltration, rendering prostate tumors more responsive to ICB. It is noteworthy that in other murine models the recruitment of PMN-MDSC and neutrophils may be driven by other chemokines, including Cxcl1^21^ and Cxcl12^22^. Our findings are corroborated by clinical data showing that PMN-MDSCs accumulate in the blood of patients with advanced prostate cancer^23–25^, and that an intratumoral PMN signature is associated with poor outcome^26^. Our data are also supported by pre-clinical studies showing that blocking MDSC function increases the efficacy of ICB in animal models of CRPC^12^. Consistent with recent data, we found that the PMN-MDSCs infiltrating prostate tumors express IL-23^14^. We further showed that inhibiting the recruitment of these cells peri-castration augmented the CD8 T cell effector function initiated by ICB. Based on these findings, we have initiated a phase 1b/2 trial (NCT03689699) to test whether adding ICB and anti-IL-8 to a short course of ADT can prevent PMN-MDSC infiltration and delay progression in men with castration-sensitive prostate cancer. In summary, targeting the IL-8/CXCR2 pathway following ADT in combination with immune checkpoint blockade may represent a novel treatment paradigm to improve responses to immunotherapy and delay the onset of castration-resistance.

## Acknowledgments

We thank members of the Drake Lab for discussion and insightful comments; F. Veglia for advice with *in vitro* suppression assays; K.C. Smith, A. Floratos, and the Center for Computational Biology and Bioinformatics at Columbia University for ChIP-Seq analysis; S. Coley, T. Swayne, E. Munteanu, and the Confocal and Specialized Microscopy Shared Resource at Columbia University for help with microscopy; L. Dasko-Vincent from the Sidney Kimmel Comprehensive Cancer Center Imaging Facility at Johns Hopkins for support with LCM, J. Pevsner for assistance on protein homology analyses, and B. Johnson for help with statistical analyses. This study was supported by U.S. Department of Defense (W81XWH-13-1-0369), U.S. National Institutes of Health National Cancer Institute (R01: CA127153), the Patrick C. Walsh Fund, the OneInSix Foundation, and the Prostate Cancer Foundation. Research reported in this publication was performed in the CCTI Flow Cytometry Core, supported in part by the Office of the Director, National Institutes of Health under awards S10OD020056. H&E/IHC staining and image collection for this work was performed in the Molecular Pathology Shared Resource and the Confocal and Specialized Microscopy Shared Resource of the Herbert Irving Comprehensive Cancer Center at Columbia University, supported by NIH grant #P30 CA013696 (National Cancer Institute). The content is solely the responsibility of the authors and does not necessarily represent the official views of the National Institutes of Health.

## Author contributions

Z.A.L.B., M.C.H., M.G.C., N.C., N.J.V., and A.O. performed experiments; C.H., J.J., C.J.B., P.J.H., M.J.S., and A.J.K. contributed essential reagents; Z.A.L.B., M.C.H., A.M.C. and C.G.D. designed and supervised experiments; M.C.H., K.S.S., and A.D.M. coordinated the study on human samples; C.G.D. supervised the study. Z.A.L.B. and C.G.D. wrote the manuscript, which was edited by all authors.

## Author information

C.G.D. has stock or ownership interests in Compugen, Harpoon, Kleo, Potenza, and Tizona Therapeutics, and has served as a consultant for Agenus, Dendreon, Janssen Oncology, Eli Lilly, Merck, AstraZeneca, MedImmune, Pierre Fabre, Genentech, and Genocea Biosciences. A.M.C. is a shareholder of Aclaris Therapeutics, Inc, and a consultant for Dermira, Inc. and Aclaris Therapeutics, Inc. Columbia University has filed a US patent claiming the benefit of U.S. Provisional Patent Application No. 62/809,060 (inventors C.G.D. and Z.A.L.B.) on the use of IL-8/CXCR2 blockade of PMN-MDSC recruitment to the TME for the treatment of prostate cancer. The remaining authors declare no competing financial interest.

## Data Availability

The data that support the findings of this study are available from the corresponding author upon reasonable request.

## Biological Materials

Biological materials used in this study may be requested from the corresponding author, with the exception of anti-CTLA-4 and anti-CXCR2 antibodies which were obtained through an MTA with A.K and M.S.

## Materials and Methods

### Patient Samples

Formalin fixed, paraffin embedded (FFPE) human prostate cancer samples were obtained from consented patients treated with ADT (degarelix; 240 mg SQ) in a neo-adjuvant trial (NCT01696877)^1^ and matched control radical prostatectomies were obtained from patients treated at the Johns Hopkins Sidney Kimmel Comprehensive Cancer Center (Baltimore, MD) under IRB-approved clinical protocol J1265. All patients provided written, informed consent.

### Cell Lines

Myc-Cap, derived from spontaneous prostate cancer in c-Myc transgenic mice ^2, 3^, was a generous gift from Dr. C. Sawyers. To generate MCRedAL, Myc-Cap cells were transfected with pRetroQ-mCherry-C1 (Clontech) using lipofectamine 2000 (Invitrogen) and isolated by FACS sorting based on mCherry expression (Extended Data Fig. 1a). Myc-Cap and MCRedAL cells were cultured in DMEM as previously described^2^. LNCaP, VCaP, E006AA, CWR22Rv1, DU145, and PC3 cell lines were obtained and cultured as recommended by the ATCC. LAPC4 (a gift from Dr. S. Yegnasubramanian) were maintained in RPMI-1640 (Corning) supplemented with 10% fetal bovine serum (FBS; Gemini Bio-Products). Androgen independent LNCaP-abl cells were a gift from Dr. Z. Culig and cultured as descrived previously^4^. LAPC4-CR and VCaP-CR (a gift from S. Yegnasubramanian) were derived by passaging LAPC4 and VCaP cells through castrated animals and further subculturing in RPMI-1640 supplemented with 10% Neuronal Supplement (Gibco). For experiments when cells were grown in androgen-free conditions, 10% FBS was substituted for 10% CSS in complete media. For migration/chemotaxis assays, prostate cancer cell lines were cultured in complete media containing either 0.5% or 2.5% FBS for human and murine cells, respectively. All cell lines were cultured in 1% penicillin/streptomycin media at 37°C, 5% CO_2_.

### Mouse Strains

Seven-week-old FVB/NJ, J:NU, C57BL/6-Tg(TcraTcrb)1100Mjb/J (OT-I), and B6.SJL-PtprcaPepcb/BoyJ (CD45.1) male mice were purchased from The Jackson Laboratory. A breeding pair of *Hoxb13-rtTA|TetO-H2BGFP* (HOXB13-GFP) mice^5^ was received from UMBC and experimental animals were bred in-house. Animals were kept in a specific pathogen-free facility at either Johns Hopkins University School of Medicine or Columbia University Medical Center. All animal experiments were performed in accordance with protocols approved by the Institutional Animal Care and Use Committee (IACUC) at the respective institutions.

### Tumor Allografts and Xenografts

Eight-week-old male FVB/NJ and J:NU mice were subcutaneously inoculated with either Myc-Cap or MCRedAL (1×10^6^ cells/mouse), and LNCaP or PC3 (3×10^6^ cells/mouse) in the right flank, respectively. Tumor diameters were measured with electronic calipers every 3 days as indicated and the tumor volume was calculated using the formula: [longest diameter × (shortest diameter)^2^]/2. Myc-Cap tumor bearing mice received androgen-deprivation therapy (ADT) 4 weeks after tumor implantation when tumor volume reached ∼500mm^3^, as indicated in figure legends. ADT was administered via subcutaneous (sc) injection of degarelix acetate (a GnRH receptor antagonist; Ferring Pharmaceuticals Inc.) at a dosage of 0.625 mg/100 μl H_2_O/25 g body weight every 30 days, unless otherwise indicated. Onset of castration-resistance was defined as the time to tumor size increased by 30% (∼650 mm^3^) after ADT. Chemical castration by ADT was compared to bilateral orchiectomy as described in Extended Data Fig. 1a.

### Luminal Epithelial Regression/Regeneration

Eight-week-old male HOXB13-GFP mice carrying the *Hoxb13-rtTA* transgene and a Tetracycline operator–Histone 2B-Green Fluorescent Protein (TetO-H2BGFP), which results on GFP expression being restricted to luminal epithelial Hoxb13^+^ cells (described previously^5^), were castrated via bilateral orchiectomy. A cycle of prostate regression/regeneration was induced as described previously^6^. Briefly, mice were allowed to regress for six weeks to reach the fully involuted state. Mice were randomized to ADT or ADT + testosterone (T) treatment groups. Testosterone was administered for four weeks for prostate regeneration by subcutaneous pellets; this regimen yields physiological levels of serum testosterone. All mice received 2mg/ml of Doxycycline (Sigma) in the drinking water to induce GFP expression^5^ under the control of the luminal epithelial promoter, HoxB13, one week prior euthanizing them for their analysis.

### Antibody Blockade

Anti-CXCR2 (murine IgG1-D265A, clone: 11C8; a non-FcγR-binding mutant with deficient FcγR-mediated depletion), anti-CSF1R (rat IgG2a, clone: AFS98; with competent FcγR-mediated depletion), and anti-CTLA-4 (murine IgG2a, clone: 12C11; with competent FcγR-mediated depletion)^7^ were used. Antibody treatment was administered via intraperitoneal (ip) injection at a dose of 50 mg/kg body weight for 3 doses every 4 days for CXCR2, 50 mg/kg body weight every 3 days for the duration of the experiment for CSF1R, and/or10 mg/kg body weight for 3 doses every 3 days for CTLA-4. Mouse IgG1 (clone: 4F7), rat IgG2a (clone: 2A3), and mouse IgG2a (clone: 4C6) were used as isotype controls. Anti-CXCR2 and anti-CSF1R treatments started 7 days before ADT; while anti-CTLA-4 treatment was started either 3 or 12 days before ADT (400mm^3^ vs. 200mm^3^, respectively).

### Flow cytometry

Single-cell suspensions from prostate tumor and tissues were prepared using the mouse tumor dissociation kit according to the manufacturer’s recommendations (Miltenyi). Single-cell suspensions of tumor-draining lymph nodes (TDLNs) and spleens were homogenized mechanically with the back of a syringe. Cells were Fc-blocked with purified rat anti-mouse CD16/CD32 (Clone: 2.4 G2, Becton Dickinson BD) for 15 minutes at RT. Dead cells were discriminated using the LIVE/DEAD (L/D) fixable viability dye eFluor 506 or near-IR dead cell stain kit (Thermo Fisher) and samples were stained for extracellular and intracellular markers. The following antibodies were used: CD45 (30F-11), CD45.2 (104), CD24 (M1/69), CD49f (GOH3), Ly6C (HK1.4), Ly6G (1A8), Gr1 (RB6-8C5), CD11b (M1/70), F4/80 (BM8), MHCII (2G9), PD-L1 (10F.9G2), CD4 (RM4-5), CD8 (53-6.7), CD44 (IM7), CD62L (MEL-14), CD25 (PC61), Ki67 (16A8), IFN-γ (XMG1.2), TNF-α (MP6-XT22), IL-2 (JES6-5H4), GZβ (GB11), CXCR2 (242216), and IL-23 (FC23CPG). For intracellular staining, cells were fixed and permeabilized using BD Perm/Wash (BD Biosciences) at room temperature for 45 minutes. For intracellular cytokine staining, cells were stimulated with PMA (50 ng/ml) and ionomycin (500 ng/ml) for 4 hours in the presence of protein transport inhibitor cocktail (eBiosciences). Gates of cytokines were determined by fluorescence minus one (FMO) controls. Staining was visualized by fluorescence activated cell sorting (FACS) analysis using a BD FACSCelesta™ (BD Biosciences) and analyzed using FlowJo® (Flowjo LLC). Prostate luminal epithelial cells are defined as CD45^-^CD11b^-^F4/80^-^CD24^+^CD49f^int^GFP^+^, and prostate epithelial tumor cells are defined as CD45^-^CD11b^-^F4/80^-^mCherry^+^. Tumor associated macrophages (TAMs) are referred to as CD45^+^CD11b^+^F4/80^+^, inflammatory TAMs as CD45^+^CD11b^+^F4/80^+^Ly6C^+^MHCII^-^, immature TAMs as CD45^+^CD11b^+^F4/80^+^Ly6C^+^MHCII^+^, MHCII^hi^ TAMs as CD45^+^CD11b^+^F4/80^+^Ly6C^-^MHCII^+^, MHCII^low^ TAMs as CD45^+^CD11b^+^F4/80^+^Ly6C^-^MHCII^-^. PMN-MDSCs are defined as CD45^+^CD11b^+^Ly6C^+^Ly6G^+^. CD4 T cells as CD45^+^CD4^+^, regulatory T cells as CD45^+^CD4^+^CD25^+^, CD8 T cells as CD45^+^CD8^+^, polyfunctional CD8 T Cells as CD45^+^CD8^+^IFNγ^+^TNFα^+^Gzβ^+^, and memory CD8 T cells as CD45^+^ CD8^+^CD44^+^CD62L^-^. 123Count eBeads counting beads (Thermo Fisher) were used to normalize the numbers of PMN-MDSCs in migration/chemotaxis experiments.

### Protein Quantification

Tumors collected at different treatment time points were minced, lysed in CelLytic MT (Sigma) containing halt protease and phosphatase inhibitor (Thermo Fisher) in a 1:100 ratio, and incubated on ice for 30 minutes with intermittent vortexing. Tumor lysates were assayed for raw protein concentration with Coomassie assay (Bio-Rad). IL-8 and Cxcl15 were analyzed by ELISA kits following the manufacturer’s instructions (BD Bioscience and R&D Systems, respectively).

### Immunohistochemical staining (IHC)

Tumor and tissue samples were fixed with either 10% formalin (Fisher Scientific, Pittsburgh, PA) or zinc fixative (BD) for 24 hours before paraffin embedding and sectioning. Sections were stained with hematoxylin and eosin (H&E), and antibodies against mouse Ly6G (1A8; BD Pharmingen) and F4/80 (BM8; eBioscience). Staining was performed by the Molecular Pathology core of the Herbert Irving Comprehensive Cancer Center at Columbia University. All images were acquired on a Leica SCN 400 system with high throughput 384 slide autoloader (SL801) and a 40X objective; files were processed with Aperio ImageScope v12.3.1.6002.

### RNA In Situ Hybridization (RISH) and Immunohistochemistry

Manual fluorescent RISH was performed on formalin-fixed and zinc-fixed paraffin embedded sections using company protocols. Briefly, 5µm sections were cut, baked at 60 ℃ for 1 hour, dewaxed, and air-dried before pre-treatments. RISH *Cxcl15* probe, 3-plex positive control probes (*Polr2a*, *Ppib*, *Ubc*) and 3-plex negative control probes (*DapB* of Bacillus subtilis strain) from Advanced Cell Diagnostics (ACD) were used in this study. Detection of specific probe binding sites was performed with RISH Multiplex Fluorescent Reagent Kit v2 Reagent kit from ACD following the manufacturer’s instructions. Tyramide

### CF568 (Biotium) was used to visualize RISH signal

For a more precise identification of cells expressing *Cxcl15*, RISH was coupled to immunohistochemistry of PanCK (Poly; Dako) and CD45 (30-F11; BD Biosciences). Immediately after RISH detection, samples were permeabilized with 0.2% TBS-Tween 20 for 10 minutes at RT, and then blocked with 2.5% of normal goat serum (Vector) for 30 minutes at RT. Primary antibody for PanCK was diluted 1/400 in renaissance background reducing diluent (Biocare Medical) and incubated overnight at 4 °C. After washing off the primary antibody, the slides were incubated 15 minutes at RT horseradish peroxidase (HRP) secondary antibody (Vector). Tyramide CF640R (Biotium) was used to visualize PanCK staining. In some cases, CD45 staining was also performed. For this, HRP signal was abolished by a 30 minute incubation at RT with PeroxAbolish (Biocare Medical) and then blocked with 2.5% of normal goat serum (Vector) for 30 minutes at RT. Primary antibody for CD45 was diluted 1/50 in renaissance background reducing diluent (Biocare Medical) and incubated 90 minutes at RT. After washing off the primary antibody, the slides were incubated 15 minutes at RT HRP-secondary antibody (Vector). Tyramide CF488A (Biotium) was used to visualize CD45 staining. All images were acquired on a Nikon A1RMP confocal microscope using a 60X objective. Comparisons of ISH-IHC results were performed using ImageJ.

### Whole Genome Expression Profiling and Analysis

MCRedAL tumor were harvested when their tumor volume reached ∼500mm^3^ (CS group), and 7 days after chemical castration (ADT). MCRedAL cells were isolated based on their mCherry^+^ CD45^-^ F4/80^-^ CD11b^-^ expression by flow sorting on a DakoCytomation MoFlo. RNA was extracted using Trizol LS (Invitrogen) and treated with DNAse-I using RNA clean & Concentrator (Zymo Research). The analysis was performed using Affymetrix Mouse Clariom D (MTA 1.0) array according to the manufacturer’s instructions. Resulting CEL files were analyzed in Affymetrix Expression Console (v. 1.4) using the SST-RMA method, and all samples passed the quality control. Log2 probe intensities were extracted from CEL (signal intensity) files and normalized using RMA quantile normalization, then further analyzed using Partek Genomics Suite v6.6. Illustrations (volcano plots, heatmaps, and histograms) were generated using TIBCO Spotfire DecisionSite with Functional Genomics. Gene set enrichment analysis (GSEA) of differently expressed genes was performed using the hallmark gene sets Molecular Signature Database (MSigDB).

### Nanostring

RNA extraction was performed using the Trizol LS reagent (Thermo Fisher) as per manufacturer’s instructions. For NanoString analysis, the nCounter mouse PanCancer Immune Profiling panel was employed using the nCounter Analysis System (NanoString, Seattle, WA). Analysis was conducted using nSolver software (NanoString). Heatmap analyses were performed using The R Project for Statistical Computing (https://www.r-project.org/).

### Pairwise Alignment

The homology of the murine chemokines Cxcl1, Cxcl2, Cxcl5, Cxcl15, Cxcl12, and Cxcl17 to human IL-8 was evaluated using BLASTP 2.9.0+ (https://blast.ncbi.nlm.nih.gov/Blast.cgi?PAGE=Proteins)^8^. Proteins were consider homologous if they shared > 30% amino acid identity^9^. Expected values of <0.05 were consider statistically significant. The expected value includes an inherent Bonferroni correction.

### Chromatin immunoprecipitation assay (ChIP)-Seq

ChIP-Seq data was obtained from https://www.ncbi.nlm.nih.gov/geo/query/acc.cgi?acc=GSE83860 which contains ChIP-Seq data acquired with androgen receptor (AR) and nuclear factor NF-kappa-B p65 subunit (p65) specific antibodies on cell lysates from LNCaP cells cultured under the following treatments: DMSO, DHT, and TNFα. For each treatment the dataset contains two ChIP-Seq replicates pulled down using the AR and p65 antibodies^10^. ChIP-Seq data were aligned to the hg38 reference version using the subread package, and then the BAM files were sorted and indexed using SAMtools. Loci with significant differential binding (FDR = 0.05) of pulled-down proteins to DNA were identified using the csaw package for ChIP-Seq analysis, closely following Lun and Smyth’s script^7^. ChIP-Seq visualization was performed using the Integrative Genomics Viewer (IGV) from the Broad Institute (http://software.broadinstitute.org/software/igv/).

### ChIP-qRT-PCR

Chromatin immunoprecipitation was performed as described^11^. In brief, LNCaP cells were washed with serum-free media and then grown in media containing 10% charcoal stripped FBS for 48 hours. Cells were treated with 100nM DHT or vehicle for 8 hours. DNA was cross-linked with 1% formaldehyde in PBS for 10 minutes and crosslinking was quenched by addition of 0.125 M glycine. Fixed cells were then lysed in lysis buffer (1% SDS, 5mM EDTA, 50mM Tris HCl, pH8.1) and sonicated to a fragment size of 200-600 bp using a Covaris water bath sonicator (Woburn, MA). Sheared chromatin was then incubated with primary antibodies (AR [06-680, Millipore], H3K4me3 [ab8580, Abcam], phospho-Ser5 RNA polymerase 2 [ab5131, Abcam], RNA polymerase 2 [4H8, Cell Signaling Technologies] or control IgG [Cell Signaling Technologies]) overnight at 4°C. Complexes were immobilized on Dynabeads (Thermo Fisher) by incubating for 4 hours at 4°C. Beads were sequentially washed with TSEI (0.1% SDS, 1% Triton X-100, 2mM EDTA, 20mM Tris HCl, pH 8.1, 150mM NaCl), TSEII (0.1% SDS, 1% Triton X-100, 2mM EDTA, 20mM Tris HCl, pH 8.1, 500mM NaCl) and TSEIII (0.25 M LiCl, 1% NP-40, 1% deoxycholate, 1mM EDTA, 10mM Tris HCl, pH 8.1). DNA was eluted with IP Elution buffer (1% SDS, 0.1M NaHCO_3_, proteinase K) and incubated at 56°C for 15 minutes. Enriched DNA libraries were analyzed using primers specific to *IL-8* locus: Forward: 5’ AGCTGCAGAAATCAGGAAGG 3’ and Reverse: 5’ TATAAAAAGCCACCGGAGCA 3’ using quantitative (q) RT-PCR. Data is shown as relative enrichment normalized to input DNA.

### Quantitative (q) RT-PCR

Total RNA was extracted using Trizol (Ambion). cDNA was prepared from total RNA preps using the RNA to cDNA EcoDry Premix (Clontech). Real-time assays were conducted using TaqMan real-time probes (Applied Biosystems). ΔΔ CT method was used for relative gene expression. Expression of the target gene was normalized to the reference gene (18S) and the mean expression level of the control group. LCM samples were normalized to 18S, TBP, and GAPDH reference genes.

### Laser Capture Microscopy (LCM)

Formalin fixed-paraffin embedded radical prostatectomy specimens, from patients enrolled in a neoadjuvant clinical trial (NCT01696877)^1^ who received 240 mg (SQ) of degarelix and matched control cases (patients that did not receive any hormone therapy), were sectioned at a thickness of 8 μm and transferred onto PEN membrane glass slides (Leica). Sections were deparaffinized, hydrated and stained with hematoxylin prior to microdissection. Individual cancer cells and cancer cell clusters were microdissected by a trained pathologist using a LMD 7000 laser capture microscope (Leica). RNA was recovered from the microdisseceted material using the RNeasy FFPE kit (Qiagen). Quantitative RT-PCR was performed as described above. For the analysis, a Mann-Whitney U test was performed.

### IL-8 and Cxcl15 CRISPR/Cas9 Knock Outs

The 20 bp long gRNA, designed using Deskgen online software, for targeting *IL-8* and *Cxcl15* in exon 3 (5’- *TTCAGTGTAAAGCTTTCTGA* −3’ and 5’- *ACAGAGCAGTCCCAAAAAAT* −3’, respectively) were incorporated into two complementary 100-mer oligonucleotides and cloned into a gRNA containing plasmid containing the (NeoR/KanR) cassette (Addgene # 41824). The human codon optimized pCAGGS-Cas9-mCherry was used for gene-editing experiments (a gift from Stem Cell Core Facility at Columbia University). gRNA and Cas9 containing plasmids were introduced to prostate epithelial cells using the basic nucleofector kit (Amaxa, Lonza) following the manufacturer’s instructions for primary mammalian epithelial cells (program W001). Successfully transfected cells were selected by culturing in the presence of 400µg/ml of neomycin sulfate analog (G418; Sigma), and isolated based on their mCherry expression 24 hours after transfection. Knock out clones were screened for IL-8 and Cxcl15 expression by ELISA and gene-editing confirmed by PCR amplification and Sanger sequencing (GENEWIZ) using primers ∼200bp away from the cut site (IL-8 Forward: 5’- TTTGGACTTAGACTTTATGCCTGAC −3; IL-8 Reverse: 5’- TCCTGGGCAAACTATGTATGG −3; Cxcl15 Forward: 5’- GCTAGGCACACTGATATGTGTTAAA −3; Cxcl15 Reverse: 5’-ACATTTGGGGATGCTACTGG −3).

### Migration/Chemotaxis Assay

Cells and supernatants used in this assay were resuspended in culture media containing 0.5% or 2.5% FBS. Transwell plates of 3-mm pore size were coated with Fibronectin (Corning Costar) and loaded with 500 ml of medium or with different cell supernatants in triplicates (lower chamber). Cells were resuspended at 2×10^7^ cells/ml, and 200 ml of this suspension was placed in each of the inserts (upper chamber). After 2.5 hours of incubation at 37°C and 5% CO_2_, inserts were removed and 10,000 beads (Thermo Fisher) were added to each well. In some cases, either isotype or anti-CXCR2 (200 µg/ml) were added at the beginning of the experiment. The cells in the lower chamber were collected along with the starting cell population, stained with L/D, CD11b, Ly6C, and Ly6G and evaluated by flow cytometry in a BD FACSCelesta™ (BD Biosciences). The ratio of beads to cells was determined, allowing calculation of the number of cells that had migrated to the bottom well. *In vivo*, LD-PMN-MDSCs were collected as described below from splenocytes of CR-Myc-Cap tumor bearing mice and labeled with DiD (DiIC18(5) or 1,1’- Dioctadecyl-3,3,3’,3’-Tetramethylindodicarbocyanine, 4-Chlorobenzenesulfonate Salt; Invitrogen), a lipophilic membrane dye, as described previously^12^. DiD^+^ LD-PMN-MDSCs were adoptively transferred into FVB/NJ recipient 8-week male mice and their ability to migrate in response to 200ng of recombinant Cxcl15 was evaluated 4 hours after injection. Beads were also used to calculate absolute numbers of Ly6G^+^ PMNs and DiD^+^ LD-PMN-MDSCs *in vivo*.

### PMN-MDSC Enrichment

Animals were sacrificed and spleens were collected. After dissociating cell clumps, the cell suspension was centrifuged (740 g, 10 minutes, RT) and resuspended in 1 ml HBSS– EDTA containing 0.5% BSA. Cells were then resuspended in 50% Percoll solution and treated on a three-layer Percoll gradient (55%, 72%, and 81%) at (1500 g, 30 minutes, 10°C without break). LD-PMN-MDSCs were collected from the 50-55% and 55-72% interfaces. Red blood cells (RBCs) were eliminated with RBC lysis solution (Miltenyi).

### In vitro Suppression Assays

PMN-MDSCs were isolated from the spleen of CR-Myc-Cap-tumor bearing mice using the neutrophil isolation kit (Miltenyi) according to the manufacturer’s instructions; greater than 95% enrichment was confirmed by flow cytometry. Unless otherwise indicated, a density gradient separation was performed prior to column purification. OT-I (CD45.2) transgenic splenocytes were mixed at a 1:10 ratio with sex-matched CD45.1 splenocytes. Splenocytes containing CD8 T responder cells were stained with CellTrace Violet (5µM CTV; Thermo Fisher) and plated on a 96-well round-bottom plate at a density of 2×10^5^ cells per well. PMN-MDSCs cells were added at 2-fold dilutions starting from 2×10^5^ cells, in the presence of their cognate peptides (5pM OVA) and incubated for 60 hours. Proliferation of CD8 T responder cells (gated as L/D^-^CD8^+^CTV^+^) was quantified by flow cytometry based on the dilution of Cell Trace Violet (CTV). Percent suppression (% Suppression) was calculated by the following formula: % Suppression = [1-(% divided cells of the condition/ the average of % divided cells of T responder only conditions)] × 100.

### Z-score Analysis

IL-8 expression was evaluated in a publicly available data set (GSE8466)^13^ using z-score values of quantile-normalized microarray transcripts from benign prostate biopsies. Z-score values were obtained by scaling the data for each gene in each patient to: (expression - mean expression across all genes) / (standard deviation of expression across all genes).

### Statistical Analysis

Statistical analysis was performed using Prism 7 (GraphPad). Unpaired two-tailed t-tests, Mann-Whitney U test, Tukey’s multiple comparisons tests, or Wilcoxon rank sum tests were conducted and considered statistically significant at *p*-values ≤0.05 (*), 0.01 (**), 0.001 (***) and 0.0001 (****).

