## Extended Data Figures for "Castration-mediated IL-8 Promotes Myeloid Infiltration and Prostate Cancer Progression"

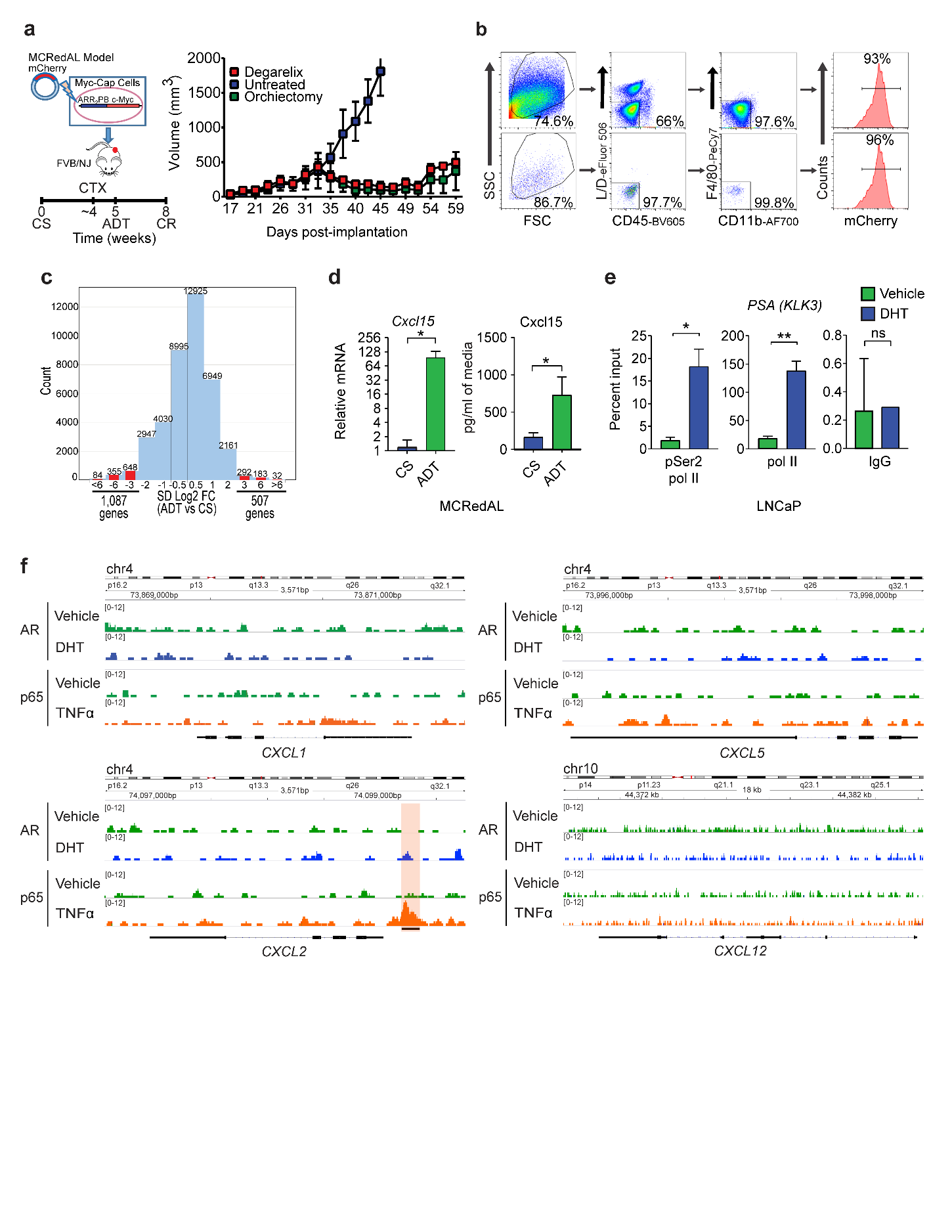
**Extended Data Figure 1 | Chemokine Regulation Upon AR signaling Stimulation and an Inflammatory Stimuli in Prostate Tumor Epithelial Cells. a,** Androgen responsive tumor epithelial cells progress from castration-sensitive (CS) to androgen responsive (ADT), and eventually developed castration-resistance (CR). CR was tumor size defined as ≥ 30% of nadir tumor volume. Left, fluorescent tagin strategy to generate mCherry^+^ Myc-Cap cells (MCRedAL cells). Right, tumor growth curve of MCRedAL tumors. CTX: Castration (n ≥ 3 per group, repeated x 2). **b,** Sorting strategy to isolate tumor epithelial cells from **a** based on their expression of mCherry and their CD45^-^CD11b^-^F4/80^-^ phenotype. **c,** Histogram of log2 fold change comparisons (SD Log2 FC) between ADT and CS groups among all the microarray transcripts (n = 3 per group). **d,** Gene and protein expression of Cxcl15 in indicated MCRedAL tumor cells *in vitro* by qRT-PCR and ELISA, respectively, replicate numbers as in **c**. **e,** Percentage input bound in ChIP–qRT-PCR assays assessing binding of pSer2 Pol II and pol II at *PSA* (*KLK3*) promoter loci in LNCaP cells treated for 24 h with or without DHT (100 nM; n = 3 per group). **f,** ChIP-Seq enrichment of AR at the *CXCR1, CXCR2, CXCR5,* and *CXCL12* promoters in LNCaP cells cultured in the presence of either vehicle (DMSO), AR signaling (DHT: 100 nM), or an inflammatory stimuli (TNFα: 1000 U/ml) (n = 2 per group; GSE83860). For **f**, loci with significant differential binding (black bar) were identified as in materials and methods. Unpaired t-tests were performed, *p*-values ≤ 0.05 (*) and 0.01 (**); *p*-values ≥ 0.05 (ns).

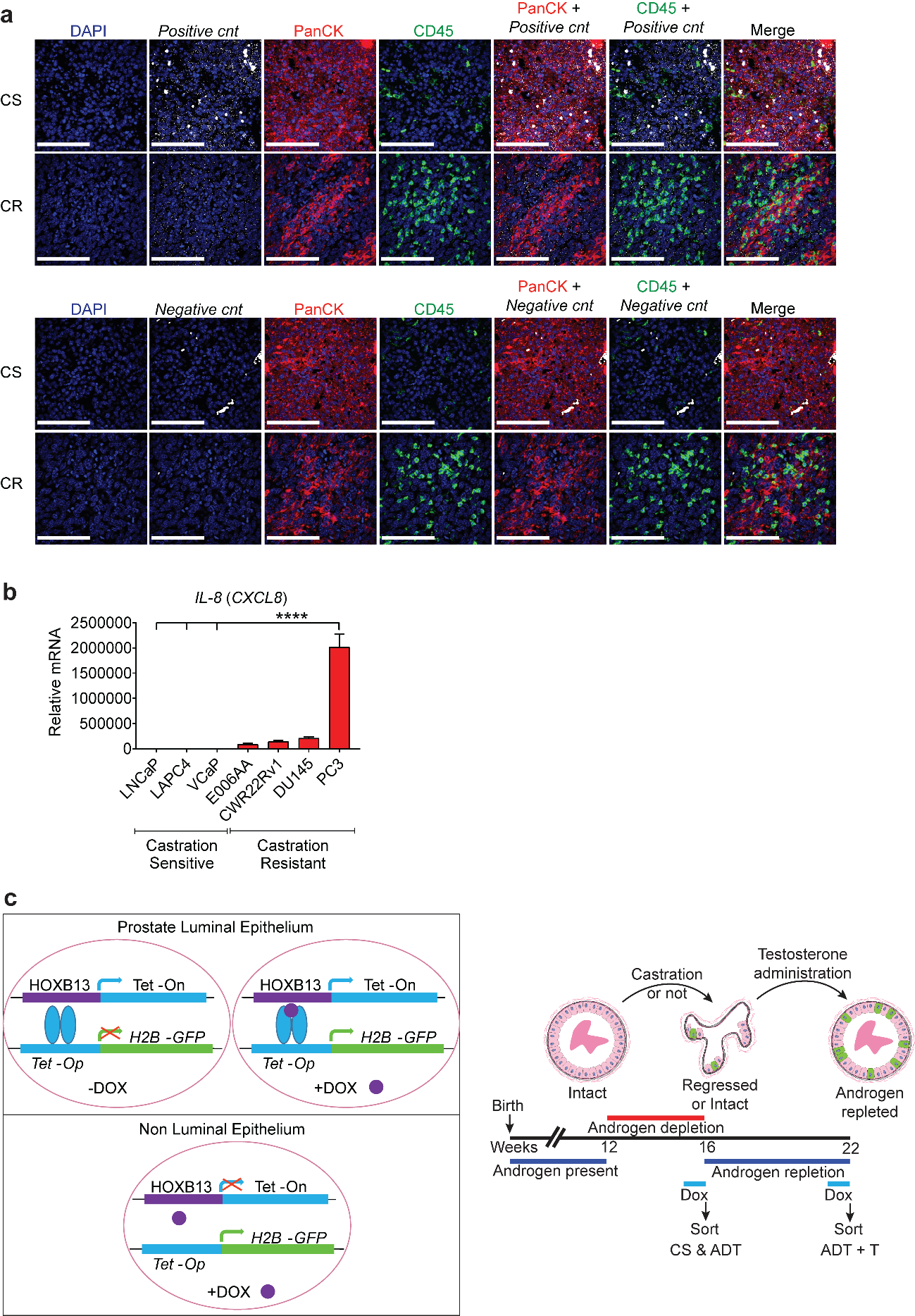
**Extended Data Figure 2 | IL-8 Expression in Mouse and Human Prostate Cancer.** **a,** Representative images of positive and negative controls (murine and bacteria probe, respectively) in Myc-Cap tumors from indicated treatment groups. Tumors were harvested when tumor volume reached ~500mm^3^ (CS group), 7 days after androgen-deprivation (ADT), or at the time of castration-resistance (CR). Prostate tumor tissue sections were hybridized with CF568-labeled probe sets (white) to *Cxcl15*, CF640-labeled anti-PanCK antibody (red), and CF488-labeled anti-CD45 antibody (green). Nuclei were counterstained with DAPI (blue). Repeated x 3. **b,** qRT-PCR quantification of *IL-8* in AR positive castration-sensitive (LNCaP, LAPC4, and VCaP) and AR independent castration-resistant (E006AA, CWR22Rv1, DU145, and PC3) human prostate cancer cell lines (n = 2 per group, repeated x 2). **c,** Non-Cancer Hoxb13-rtTA|TetO-H2BGFP mouse model to study the cells-of-origin of prostate cancer using luminal epithelial cells isolated from castration-sensitive (CS), androgen-deprivation treated (ADT) non-tumor bearing mice, and ADT treated mice that received testosterone repletion (ADT + T). Left, schematic representation of the effect of Doxycycline (DOX) on epithelial cells from a prostate duct. Right, schematic representation of the response of GFP^+^ murine luminal prostate epithelial cells to androgen-deprivation therapy (ADT) in a prostate duct. RISH images are 60X magnification; scale bar = 100 μm. Tukey’s multiple comparisons test with a single pooled variance was performed, *p*-values ≤ 0.0001 (****).

-
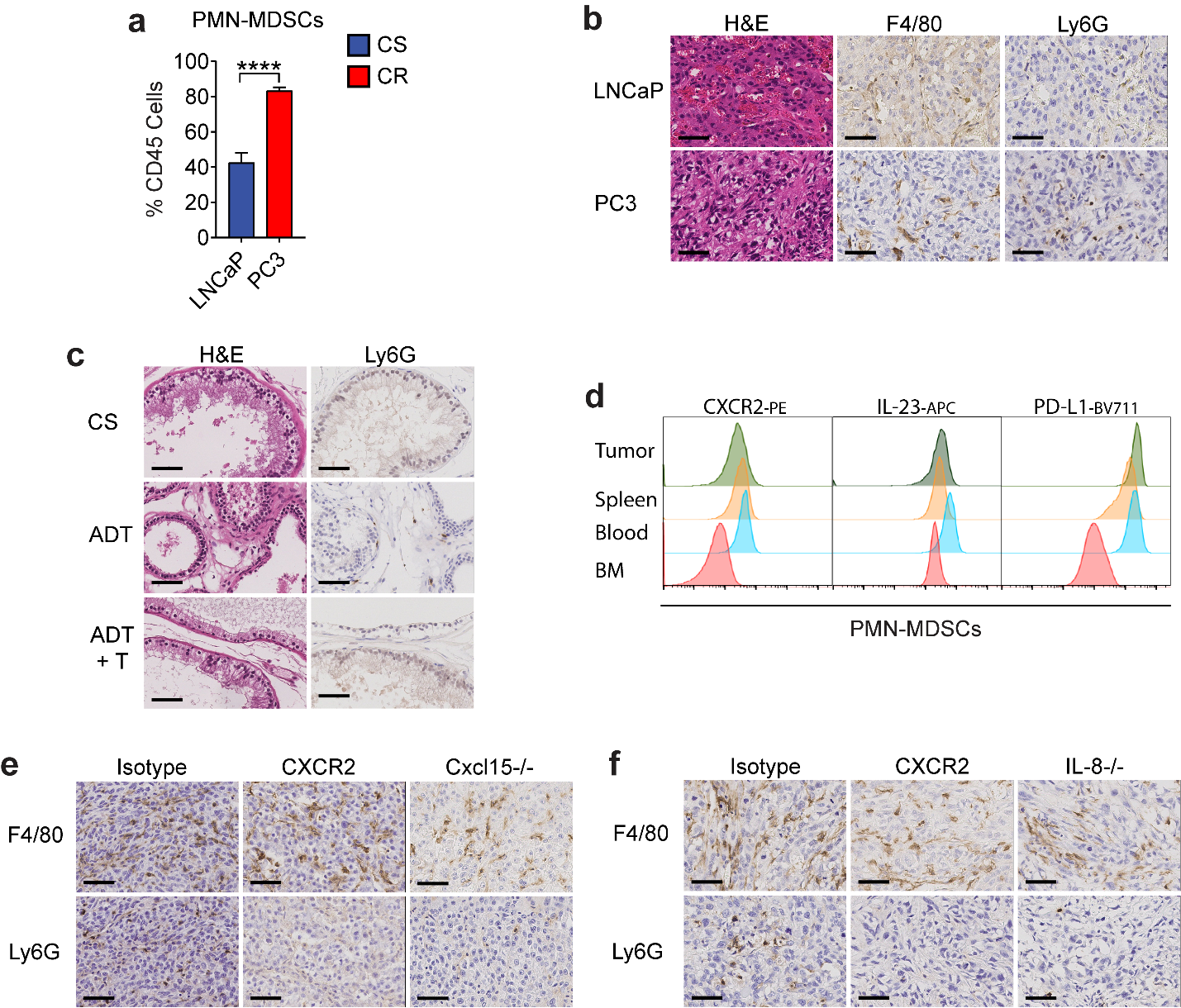
**Extended Data Figure 3 | PMN-MDSCs Infiltration Relays on IL-8 (Cxcl15) / CXCR2 Signaling Following ADT.** **a,** PMN-MDSCs as a percentage of CD45^+^ cells in the TME of indicated human prostate tumors as determined by flow cytometry (n = 3 per group, repeated x 2). **b,** Representative H&E and immunohistochemistry (Ly6G and F4/80) sections of the indicated human prostate xenografts (repeated x 3). **c,** Representative H&E and immunohistochemistry of Ly6G in non-cancerous murine prostate from castration-sensitive (CS), androgen-deprivation treated (ADT) non-tumor bearing mice, and ADT treated mice that received testosterone repletion (ADT + T). Repeated x 2. **d,** Representative histograms of protein expression determined by flow cytometry in PMN-MDSCs from indicated organs (repeated x 2). **e,** Representative H&E and immunohistochemistry (Ly6G and F4/80) on CR-Myc-Cap allografts treated as indicated (repeated x 3). **f,** Representative H&E and immunohistochemistry (Ly6G and F4/80) in PC3 tumor xenografts treated as indicated (repeated x 3). H&E and IHC images are 40X magnification; scale bar = 50 μm. Unpaired t-tests were performed, *p*-values ≤ 0.0001 (****).
-
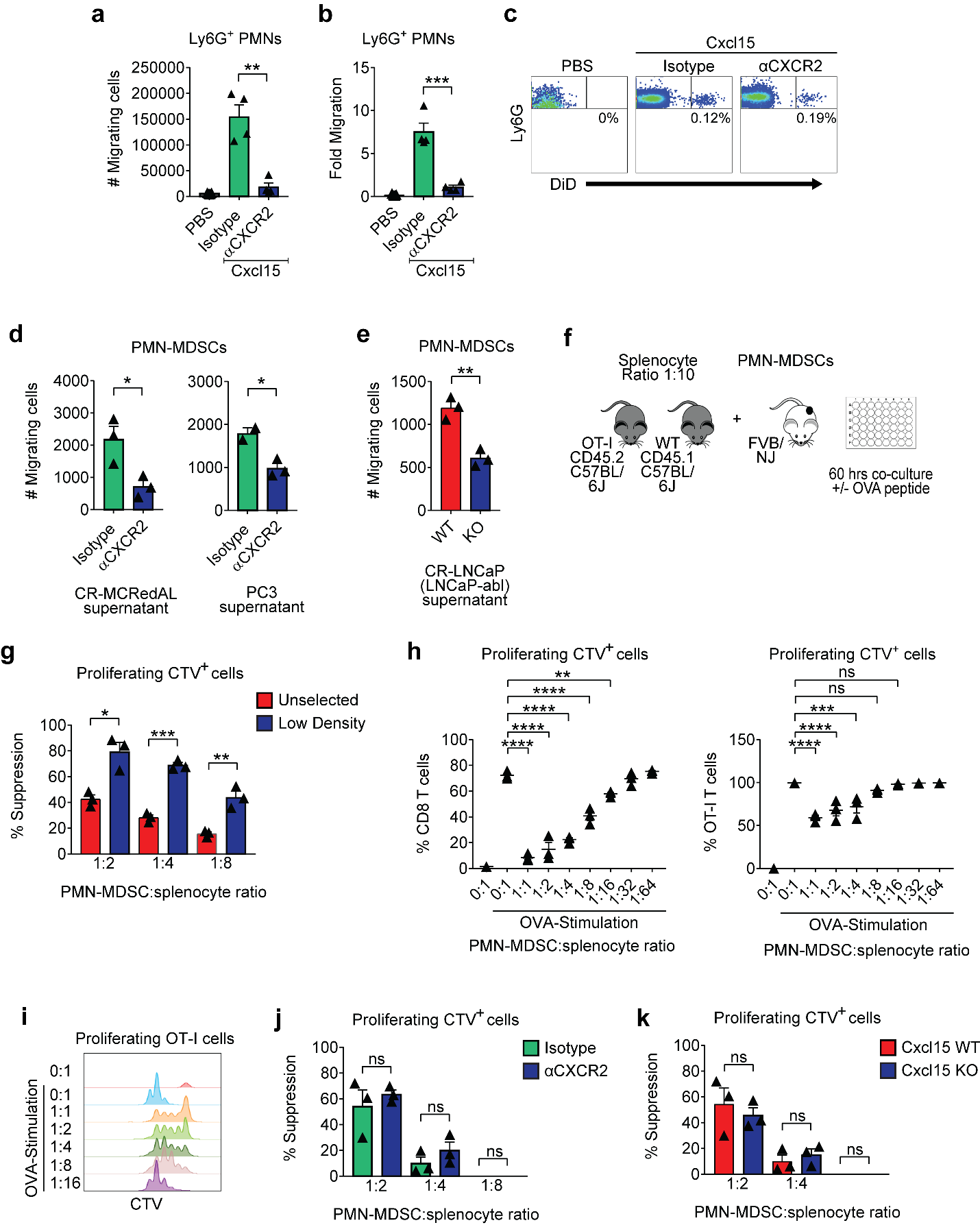
**Extended Data Figure 4 | PMN-MDSCs Migrate in Response to Cxcl15 and Suppress Antigen Specific T Cell Function.** **a,** Analysis of Ly6G^+^ PMNs in peritoneal washings receiving Cxcl15 (200ng/mouse, i.p.) in mice pre-treated with either isotype or αCXCR2 (n ≥ 4 per group, repeated x 2). **b,** Analysis of the fold change between the number of Ly6G^+^ PMNs in peritoneal washings from **a** in relation to PMNs’ numbers in peripheral blood of indicated treated mice. **c,** Representative plots of Ly6G^+^ PMNs in peritoneal washings from **a** of indicated treated mice (repeated x 2). **d,** PMN-MDSC *in vitro* migration towards tumor supernatants in the presence of either isotype or anti-CXCR2 (200 µg/ml). Antibodies were added at the beginning of the experiment (n ≥ 2 per group, repeated x 2). **e,** PMN-MDSC *in vitro* migration towards CR-LNCaP (LNCaP-abl) WT or IL-8 KO tumor supernatants (n = 3 per group, repeated x 2). **f,** Schematic representation of PMN-MDSC suppression assay. OT-I splenocytes (CD45.2) were mixed with naïve splenocytes (CD45.1) in a 1:10 ratio, labeled with CTV, and co-culture with PMN-MDSCs at the indicated ratios. T cell proliferation was stimulated (Stim) by OVA peptide (5pM) for 60 hours. **g,** Percent suppression when either unselected or low-density PMN-MDSCs were used for the experiment (n = 3 per group, repeated x 3). **h,** Percent of CD8 T cells (left) and antigen specific OT-I cells (CD45.2; right) proliferating at different proportions of PMN-MDSCs when stimulated with or without 5pM of OVA, replicate numbers as in **g**. **i,** Representative histograms of antigen specific OT-I cells proliferation based on the dilution of CTV dye when stimulated as in **h** (repeated x 2). **j,** Percent suppression in the presence of either isotype or anti-CXCR2 (200 µg/ml). Antibodies were added at the beginning of the experiment (n = 3 per group, repeated x 2). **k,** Percent suppression of PMN-MDSCs derived from spleens of WT or Cxcl15 KO Myc-Cap tumor bearing mice (n = 3 per group, repeated x 2). For **a-c**, PMNs were gated on CD45^+^Ly6G^+^ cells. Cell migration *in vivo* was evaluated 4 hours after PBS or cytokine treatment and normalized to 10,000 beads. PBS was injected as the control for these experiments. For **d-e**, PMN-MDSCs were isolated from spleens of mice bearing CR-Myc-Cap tumors and placed in the top chamber of a transwell. Culture supernatants were plated in the bottom chamber, and number of PMN-MDSCs migrating from the top to the bottom chamber after 2.5 hours was evaluated. For **g, j-k**, percent suppression (% Suppression) was calculated by the following formula: % Suppression = [1-(% divided cells of the condition / the average of % divided cells of T responder only conditions)] x 100. Unpaired t-tests were performed, *p*-values ≤ 0.05 (*), 0.01 (**), 0.001 (***) and 0.0001 (****); *p*-values ≥ 0.05 (ns).

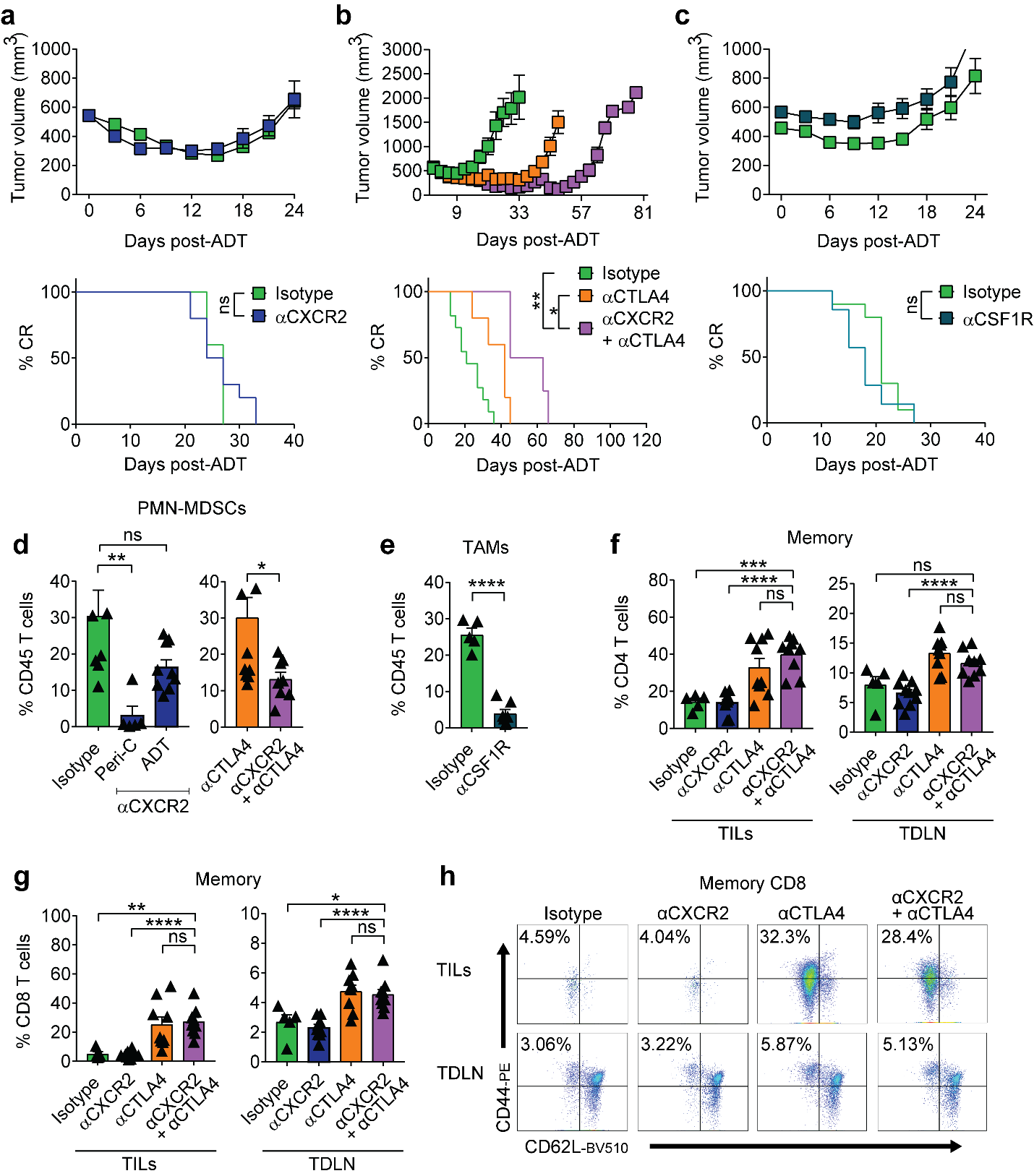
**Extended Data Figure 5 | The Therapeutic Effect of the Triple Combination is Associated with PMN-MDSC Reduction.** **a,** Tumor growth and survival curves of mice from isotype vs. αCXCR2 treatment groups (green vs. blue, respectively; n = 10 per group, repeated x 2). **b,** Tumor growth and survival curves of mice from isotype vs. αCTLA-4 vs. αCTLA-4 + αCXCR2 treatment groups (green vs. orange vs. purple, respectively; n ≥ 7 per group, repeated x 2). **c,** Tumor growth and survival curves of mice from isotype vs. αCSF1R treatment groups (green vs. purple, respectively; n ≥ 7 per group, repeated x 2). **d,** PMN-MDSCs as a percentage of CD45^+^ cells in the TME of indicated treatment groups, replicate numbers as in **b**. **e,** TAMs as a percentage of CD45^+^ cells in the TME of indicated treatment groups, replicate numbers as in **c**. **f,** Memory CD4 T cells as a percentage of CD45^+^CD4^+^ T cells in the tumor (tumor infiltrating lymphocytes: TILs) and tumor-draining lymph node (TDLN) of indicated treatment groups (n ≥ 5 per group, repeated x 2). **g,** Memory CD8 T cells as a percentage of CD45^+^CD8^+^ TILs and TDLN of indicated treatment groups, replicate numbers as in **f**. **h,** Representative plot of memory CD8^+^ TILs and TDLN of indicated treatment groups (repeated x 2). For **a-c**, treatment started when tumor volumes reached 400mm^3^. For **d-h**, treatment started when tumor volumes reached 200mm^3^. Average tumor volume (±s.e.m.) for each experimental group. Wilcoxon test was used for survival analysis. Flow cytometry as in materials and methods. Unpaired t-tests were performed, *p*-values ≤ 0.05 (*), 0.01 (**), 0.001 (***) and 0.0001 (****); *p*-values ≥ 0.05 (ns).
