## Extended Data Table 1 for "Castration-mediated IL-8 Promotes Myeloid Infiltration and Prostate Cancer Progression"

**Extended Data Table 1 | Amino Acid Sequence Homology Between human IL-8 (CXCL8) and the Murine Homologues.**

| RefSeq Accession # | Protein | Score | Expect | Identities | ELR/CXC motifs |
|---|---|---|---|---|---|
| NP_000575.1 | CXCL8 | 202 bits (514) | 1e-19 | 99/99 (100%) | **ELRCQCI** |
| EDL05301.1 | Cxcl1 | 78.6 bits (192) | 9e-26 | 35/66 (53%) | **ELRC**L**C**L |
| NP_033166.1 | Cxcl2 | 68.9 bits (167) | 6e-22 | 34/76 (45%) | **ELRC**L**C**L |
| NP_033167.2 | Cxcl5 | 63.5 bits (153) | 3e-19 | 27/66 (41%) | **ELR**VL**C**L |
| NP_035469.1 | Cxcl15 | 47.4 bits (111) | 8e-13 | 26/72 (36%) | **ELRC**L**CI** |
| AAH06640.1 | Cxcl12 | 39.7 bits (91) | 1e-10 | 26/82 (32%) | **SY**RC**P**C**-** |
| NP_705804.2 | Cxcl17 | No significant similarity found | | | |
