## Extended Data Table 2 for "Castration-mediated IL-8 Promotes Myeloid Infiltration and Prostate Cancer Progression"

**Extended Data Table 2 | NanoString nCounter Gene Expression Assay.**

| Gene Information | | | Immature TAMs | | | MHCII$^{hi}$ TAMs | | | MHCII$^{low}$ TAMs | | | PMN-MDSCs | | |
|---|---|---|---|---|---|---|---|---|---|---|---|---|---|---|
| **Gene Name** | **Accession #** | **Probe Name** | **A** | **B** | **C** | **A2** | **B2** | **C2** | **A3** | **B3** | **C3** | **A4** | **B4** | **C4** |
| A2m | NM_175628.3 | NM_175628.3:2128 | 0.39 | 0.08 | 2.13 | 0.11 | 0.08 | 0.7 | 0 | 0 | 1.61 | 1.75 | 0.78 | 0.02 |
| Abca1 | NM_013454.3 | NM_013454.3:6865 | 11.47 | 11.9 | 11.66 | 11.05 | 11.36 | 11.57 | 10.87 | 11.66 | 12.05 | 10.15 | 11.24 | 10.78 |
| Abcb1a | NM_011076.1 | NM_011076.1:2600 | 5.88 | 5.43 | 2.13 | 5.02 | 4.09 | 0.87 | 7.38 | 4.09 | 5.29 | 6.71 | 0.78 | 0.02 |
| Abcg1 | NM_009593.1 | NM_009593.1:295 | 0.39 | 0.08 | 2.13 | 1.2 | 4.26 | 0.7 | 1.32 | 0 | 4.58 | 2.75 | 0.78 | 0.02 |
| Abl1 | NM_009594.4 | NM_009594.4:1378 | 6.88 | 8.66 | 5.86 | 6.28 | 7.2 | 5.95 | 6.66 | 5.26 | 10.03 | 7.16 | 4.62 | 5.82 |
| Ada | NM_007398.3 | NM_007398.3:795 | 10.12 | 10.26 | 8.91 | 10.9 | 10.7 | 9.19 | 10.21 | 10.93 | 10.25 | 10.14 | 10.35 | 8.4 |
| Adora2a | NM_009630.2 | NM_009630.2:2306 | 3.56 | 0.08 | 2.13 | 2.16 | 0.08 | 0.7 | 0 | 0 | 1.61 | 0 | 0.78 | 0.02 |
| Aicda | NM_009645.2 | NM_009645.2:552 | 6.32 | 7.78 | 2.13 | 5.16 | 4.99 | 5.91 | 7 | 0 | 7.41 | 7.37 | 6.18 | 4.33 |
| Aire | NM_009646.1 | NM_009646.1:656 | 3.39 | 3.03 | 4.53 | 3.45 | 2.13 | 1.79 | 1.06 | 0.98 | 1.61 | 3.82 | 2.91 | 1.97 |
| Akt3 | NM_011785.3 | NM_011785.3:2494 | 8.49 | 8.99 | 9.1 | 7.79 | 8.31 | 8.53 | 7.83 | 8.11 | 9.1 | 8.02 | 8.13 | 7.91 |
| Alcam | NM_009655.1 | NM_009655.1:2605 | 7.83 | 8.35 | 8.25 | 7.53 | 8.38 | 7.86 | 7.58 | 8.59 | 8.57 | 6.83 | 6.79 | 7.74 |
| Ambp | NM_007443.3 | NM_007443.3:410 | 0.39 | 7.71 | 6.61 | 5.02 | 4 | 4.96 | 1.54 | 6.91 | 1.61 | 7.13 | 8.93 | 7.9 |
| Amica1 | NM_001005421.4 | NM_001005421.4:1002 | 6.25 | 6.78 | 5.18 | 5.47 | 6.15 | 6.29 | 3.64 | 5.8 | 3.56 | 7.77 | 9.31 | 9.82 |
| Angpt1 | NM_009640.3 | NM_009640.3:1350 | 0.39 | 0.08 | 2.13 | 0.11 | 0.08 | 0.7 | 0 | 0 | 1.61 | 0 | 0.78 | 0.02 |
| Angpt2 | NM_007426.3 | NM_007426.3:2020 | 3.71 | 4.87 | 6.4 | 6.85 | 5.61 | 5.17 | 7.54 | 7.2 | 7.67 | 5.5 | 4.72 | 8.54 |
| Anp32b | NM_130889.2 | NM_130889.2:480 | 12.1 | 12.11 | 12 | 12.04 | 11.78 | 11.68 | 12.1 | 11.97 | 12.35 | 12.52 | 12.11 | 12.73 |
| Anxa1 | NM_010730.2 | NM_010730.2:400 | 8.05 | 8.88 | 9.06 | 7.27 | 8.2 | 8.39 | 8.42 | 9.16 | 9.45 | 9.68 | 9.18 | 10.38 |
| Apoe | NM_009696.3 | NM_009696.3:129 | 13.92 | 14.14 | 13.76 | 13.28 | 14.07 | 13.18 | 13 | 13.57 | 14.2 | 12.25 | 11.38 | 12.18 |
| App | NM_007471.2 | NM_007471.2:511 | 9.72 | 9.29 | 8.78 | 9.52 | 9.15 | 8.73 | 9.93 | 9.87 | 9.46 | 9.22 | 8.95 | 9.15 |
| Arg1 | NM_007482.3 | NM_007482.3:626 | 10.31 | 9.95 | 10.65 | 9.69 | 10.07 | 10.27 | 9.44 | 10.24 | 10.81 | 9.34 | 9.36 | 9.35 |
| Arg2 | NM_009705.2 | NM_009705.2:249 | 6.54 | 6.36 | 5.86 | 4.37 | 2.7 | 4.96 | 4.73 | 7.22 | 7.98 | 8.26 | 9.53 | 10.34 |
| Atf1 | NM_007497.3 | NM_007497.3:1216 | 8.27 | 8.2 | 8.16 | 8.21 | 8.29 | 8.18 | 7.46 | 8.54 | 8.72 | 8.83 | 9.09 | 9.56 |
| Atf2 | NM_001284371.1 | NM_001284371.1:1392 | 5.09 | 7.53 | 6.24 | 5.73 | 6.8 | 6.26 | 6.18 | 6.33 | 6.69 | 8.4 | 7.98 | 8.42 |
| Atg10 | NM_025770.3 | NM_025770.3:46 | 5.34 | 2.6 | 2.13 | 4.58 | 0.08 | 0.7 | 5.94 | 4.02 | 1.61 | 0.43 | 0.78 | 5.33 |
| Atg12 | NM_026217.1 | NM_026217.1:1800 | 7.44 | 7 | 6.79 | 7.55 | 7.6 | 7.27 | 7.05 | 7.5 | 6.3 | 7.32 | 6.15 | 7.92 |
| Atg16l1 | NM_001205391.1 | NM_001205391.1:280 | 8.98 | 8.65 | 7.31 | 8.11 | 7.08 | 7.68 | 5.86 | 8.49 | 7.06 | 8.85 | 8.54 | 9.61 |
| Atg5 | NM_053069.5 | NM_053069.5:2142 | 11.88 | 11.37 | 11.79 | 11.98 | 11.34 | 12.28 | 10.63 | 10.95 | 11.65 | 10.89 | 11.65 | 11.59 |
| Atg7 | NM_028835.1 | NM_028835.1:855 | 0.39 | 0.08 | 2.13 | 0.11 | 0.08 | 0.7 | 0 | 0 | 1.61 | 4.24 | 0.78 | 0.02 |
| Atm | NM_007499.2 | NM_007499.2:5543 | 4.85 | 3.36 | 3.3 | 3.71 | 4.34 | 6.7 | 0.73 | 7.3 | 6.46 | 5.5 | 6 | 6.66 |
| Axl | NM_009465.3 | NM_009465.3:3820 | 12.14 | 12.29 | 12.38 | 12.46 | 12.04 | 12.28 | 10.96 | 11.49 | 11.91 | 10.57 | 10.15 | 10.9 |
| Batf | NM_016767.2 | NM_016767.2:750 | 8.92 | 7.37 | 7.54 | 8.16 | 8.05 | 7.29 | 7.57 | 8.03 | 9.03 | 8.99 | 8.54 | 10.03 |
| Bax | NM_007527.3 | NM_007527.3:735 | 11.65 | 11.83 | 11.07 | 11.45 | 11.14 | 10.87 | 11.18 | 11.61 | 11.48 | 11.55 | 11.32 | 12.07 |
| Bcl10 | NM_009740.1 | NM_009740.1:1168 | 11.49 | 11.57 | 11.25 | 11.25 | 11.23 | 10.95 | 11 | 11.25 | 11.39 | 11.45 | 11.5 | 12.32 |
| Bcl2 | NM_009741.3 | NM_009741.3:1844 | 11.72 | 12.14 | 11.4 | 11.79 | 11.32 | 11.57 | 10.86 | 11.57 | 11.62 | 11.03 | 11.39 | 11.5 |
| Bcl2l1 | NM_009743.4 | NM_009743.4:200 | 6.67 | 5.8 | 7.31 | 5.64 | 6.37 | 6.35 | 5.97 | 6.85 | 7.65 | 6.18 | 5.96 | 7.38 |
| Bcl6 | NM_009744.3 | NM_009744.3:185 | 9.71 | 8.14 | 9.5 | 7.95 | 7.78 | 7.86 | 8.04 | 7.94 | 9.41 | 8.92 | 10.58 | 10.14 |
| Bid | NM_007544.3 | NM_007544.3:1307 | 6.3 | 8.28 | 5.97 | 7.5 | 6.42 | 4.18 | 4.29 | 6.41 | 2.51 | 6.35 | 6.22 | 9.3 |
| Birc5 | NM_009689.2 | NM_009689.2:237 | 6.32 | 0.08 | 2.13 | 4.37 | 6.65 | 6.35 | 6.94 | 5.17 | 5.59 | 6.98 | 5.6 | 5.26 |
| Blk | NM_007549.2 | NM_007549.2:2198 | 5.25 | 0.08 | 4.22 | 0.11 | 0.08 | 0.7 | 0 | 0 | 1.61 | 0 | 0.78 | 0.02 |
| Blnk | NM_008528.4 | NM_008528.4:1546 | 7.2 | 7.9 | 6.47 | 8.77 | 8.12 | 7.99 | 7.41 | 7.56 | 6.89 | 7.55 | 0.78 | 6.74 |
| Bmi1 | NM_007552.4 | NM_007552.4:3354 | 6.85 | 8.21 | 6.07 | 8.44 | 7.54 | 8.17 | 7.81 | 7.81 | 7.03 | 8.33 | 8.43 | 7.53 |
| Bst1 | NM_009763.3 | NM_009763.3:854 | 9.17 | 9.79 | 9.88 | 9.05 | 9.29 | 9.78 | 8.94 | 9.15 | 10.18 | 10.76 | 11.81 | 12.35 |
| Bst2 | NM_198095.2 | NM_198095.2:468 | 6.58 | 6.98 | 7.15 | 7.02 | 6.33 | 6.1 | 6.01 | 6.46 | 7.36 | 6.72 | 5.44 | 5.6 |
| Btk | NM_013482.2 | NM_013482.2:2255 | 7.69 | 8.23 | 7.15 | 7.95 | 6.85 | 8.27 | 8.04 | 7.01 | 8.59 | 6.24 | 8.17 | 7.61 |
| Btla | NM_001037719.2 | NM_001037719.2:24 | 0.39 | 6.45 | 2.13 | 0.11 | 7.68 | 0.7 | 0 | 0 | 1.61 | 0 | 0.78 | 0.02 |
| C1qa | NM_007572.2 | NM_007572.2:566 | 9.2 | 9.74 | 8.02 | 7.8 | 10.19 | 9.65 | 9.71 | 9.19 | 9.3 | 8.58 | 2.03 | 8.93 |
| C1qb | NM_009777.2 | NM_009777.2:865 | 14.92 | 14.96 | 14.01 | 15.08 | 14.73 | 14.59 | 14.51 | 14.61 | 14.21 | 14.32 | 13.92 | 14.56 |
| C1qbp | NM_007573.2 | NM_007573.2:630 | 9.66 | 9.28 | 8.95 | 9.95 | 9.25 | 9.27 | 9.61 | 9.4 | 9.47 | 10.72 | 10 | 10.14 |
| C1ra | NM_023143.3 | NM_023143.3:1923 | 0.39 | 6.57 | 6.54 | 4.29 | 5.55 | 1.79 | 4.19 | 5.26 | 7.87 | 3.13 | 5.13 | 4.4 |
| C1s1 | NM_144938.2 | NM_144938.2:2490 | 0.39 | 0.08 | 2.13 | 1.76 | 0.08 | 0.7 | 0 | 0 | 4.76 | 0 | 0.78 | 0.02 |
| C2 | NM_013484.2 | NM_013484.2:2359 | 0.39 | 3.21 | 2.13 | 0.11 | 4.79 | 0.7 | 0.73 | 2.12 | 1.61 | 5.07 | 0.78 | 0.02 |
| C3 | NM_009778.2 | NM_009778.2:285 | 5.34 | 6.26 | 5.75 | 0.11 | 0.08 | 4.18 | 4.49 | 3.52 | 6.46 | 8.22 | 7.5 | 8.93 |
| C3ar1 | NM_009779.2 | NM_009779.2:555 | 11.05 | 11.36 | 10.57 | 10.42 | 10.7 | 10.61 | 10.74 | 11.01 | 11.26 | 9.75 | 9.35 | 9.79 |
| C4b | NM_009780.2 | NM_009780.2:491 | 11.04 | 9.17 | 9.6 | 10.8 | 10.5 | 10.06 | 9.63 | 9.82 | 8.58 | 9.32 | 6.59 | 9.1 |
| C5ar1 | NM_007577.3 | NM_007577.3:595 | 9.57 | 10.17 | 9.39 | 7.56 | 9.69 | 8.48 | 9.66 | 9.9 | 10.43 | 9.66 | 10.4 | 10.66 |
| C6 | NM_016704.2 | NM_016704.2:170 | 6.32 | 0.08 | 2.13 | 4.44 | 0.08 | 3.72 | 0 | 0 | 1.61 | 1.43 | 0.78 | 0.02 |
| C7 | XM_356827.6 | XM_356827.6:215 | 6.03 | 0.08 | 2.13 | 3.45 | 0.08 | 0.7 | 0 | 0 | 1.61 | 4.52 | 0.78 | 4 |
| C8a | NM_146148.2 | NM_146148.2:1022 | 0.39 | 0.08 | 2.13 | 0.11 | 0.08 | 0.7 | 0 | 0 | 1.61 | 0 | 0.78 | 4.25 |
| C8b | NM_133882.2 | NM_133882.2:1200 | 0.39 | 0.08 | 2.13 | 0.11 | 0.08 | 0.7 | 0 | 0 | 1.61 | 0 | 2.91 | 0.02 |
| C8g | NM_027062.1 | NM_027062.1:770 | 4.48 | 7.53 | 6.67 | 6.77 | 8.1 | 5.87 | 6.57 | 2.57 | 4.39 | 6.03 | 6.37 | 6.71 |
| C9 | NM_013485.1 | NM_013485.1:226 | 0.39 | 2.33 | 2.13 | 0.11 | 0.08 | 0.7 | 0 | 0 | 1.61 | 0 | 0.78 | 0.02 |
| CD209e | NM_130905.2 | NM_130905.2:696 | 0.39 | 0.08 | 2.13 | 0.11 | 0.08 | 0.7 | 0 | 0 | 1.61 | 0 | 0.78 | 0.02 |
| Camp | NM_009921.2 | NM_009921.2:355 | 0.39 | 0.08 | 2.13 | 0.11 | 0.08 | 0.7 | 0 | 0 | 1.61 | 8.59 | 8.48 | 9.14 |
| Card11 | NM_175362.2 | NM_175362.2:545 | 3.39 | 5.77 | 6.32 | 1.76 | 5.58 | 5.29 | 0 | 3.42 | 4.91 | 6.94 | 1.24 | 1.54 |
| Card9 | NM_001037747.1 | NM_001037747.1:210 | 0.39 | 0.08 | 2.13 | 0.11 | 2.91 | 0.7 | 2.06 | 1.47 | 1.61 | 0 | 0.78 | 0.02 |
| Casp1 | NM_009807.2 | NM_009807.2:259 | 9.87 | 10.16 | 9.09 | 9.32 | 9.93 | 9.63 | 9.17 | 9.5 | 9.42 | 9.4 | 9.58 | 8.41 |
| Casp3 | NM_009810.2 | NM_009810.2:630 | 5.52 | 0.89 | 2.13 | 6.07 | 5.17 | 6.2 | 1.06 | 4.28 | 5.05 | 5.13 | 0.78 | 3.91 |

| | | | | | | | | | | | | | | |
|---|---|---|---|---|---|---|---|---|---|---|---|---|---|---|
| Casp8 | NM_009812.2 | NM_009812.2:1463 | 8.69 | 8.84 | 8.27 | 8.37 | 8.08 | 7.77 | 7.15 | 8.2 | 8.73 | 7.83 | 6.74 | 7.38 |
| Ccl1 | NM_011329.2 | NM_011329.2:145 | 5.09 | 3.75 | 7.01 | 4.44 | 5.17 | 5.71 | 3.19 | 4.64 | 7.03 | 3.24 | 5.92 | 4.92 |
| Ccl11 | NM_011330.3 | NM_011330.3:430 | 0.39 | 5.56 | 2.13 | 0.11 | 0.08 | 0.7 | 4.13 | 4.15 | 3.56 | 0 | 0.78 | 3.7 |
| Ccl12 | NM_011331.2 | NM_011331.2:56 | 10.31 | 10.07 | 9.57 | 10.68 | 9.8 | 9.93 | 10.03 | 9.95 | 10.01 | 8.98 | 8.34 | 9.23 |
| Ccl17 | NM_011332.2 | NM_011332.2:247 | 4.91 | 6.69 | 7.31 | 6.56 | 7.87 | 7.5 | 0 | 0 | 1.61 | 0 | 0.78 | 0.02 |
| Ccl19 | NM_011888.2 | NM_011888.2:465 | 0.39 | 0.08 | 2.13 | 0.11 | 0.08 | 0.7 | 0 | 0 | 1.61 | 1.75 | 5.6 | 3.46 |
| Ccl2 | NM_011333.3 | NM_011333.3:415 | 13.25 | 13.27 | 13.08 | 13.4 | 12.55 | 12.66 | 12.72 | 12.87 | 13.36 | 12.29 | 12.14 | 11.92 |
| Ccl20 | NM_016960.1 | NM_016960.1:120 | 0.39 | 0.08 | 2.13 | 0.11 | 0.08 | 0.7 | 0 | 0 | 1.61 | 0 | 0.78 | 0.02 |
| Ccl21a | NM_011124.4 | NM_011124.4:170 | 0.39 | 0.08 | 2.13 | 0.11 | 0.08 | 0.7 | 0 | 0 | 1.61 | 0 | 0.78 | 0.02 |
| Ccl22 | NM_009137.2 | NM_009137.2:1096 | 0.39 | 0.08 | 2.13 | 1.2 | 0.08 | 1.79 | 0 | 0 | 1.61 | 0 | 0.78 | 0.02 |
| Ccl24 | NM_019577.4 | NM_019577.4:335 | 8.89 | 9.07 | 7.97 | 5.92 | 5.94 | 7.33 | 7.37 | 7.13 | 9.02 | 7.71 | 5.32 | 4.92 |
| Ccl25 | NM_009138.3 | NM_009138.3:316 | 0.39 | 5.2 | 2.13 | 0.11 | 0.08 | 0.7 | 0 | 2.57 | 1.61 | 0 | 0.78 | 0.02 |
| Ccl26 | NM_001013412.2 | NM_001013412.2:140 | 5.03 | 4.71 | 4.53 | 4.93 | 4.55 | 4.8 | 2.9 | 4.54 | 5.5 | 3.01 | 2.53 | 0.92 |
| Ccl27a | NM_001048179.1 | NM_001048179.1:265 | 0.39 | 0.08 | 2.13 | 0.11 | 0.08 | 0.7 | 3.44 | 0.98 | 4.58 | 3.6 | 0.78 | 1.97 |
| Ccl28 | NM_020279.3 | NM_020279.3:234 | 4.09 | 0.89 | 2.46 | 2.47 | 4.18 | 5.66 | 4.67 | 5.72 | 3.13 | 6.38 | 4.29 | 6.6 |
| Ccl3 | NM_011337.1 | NM_011337.1:60 | 9.91 | 9.72 | 8.97 | 9.26 | 8.35 | 8.39 | 9.27 | 8.65 | 9.36 | 11.11 | 11.13 | 11.86 |
| Ccl4 | NM_013652.1 | NM_013652.1:140 | 10.18 | 10.26 | 9.84 | 9.63 | 9.84 | 9.32 | 9.38 | 9.71 | 10.3 | 9.11 | 10.03 | 10.05 |
| Ccl5 | NM_013653.3 | NM_013653.3:240 | 8.68 | 9.25 | 8.21 | 8.82 | 8.48 | 8.76 | 5.81 | 7.16 | 6.73 | 7.55 | 5.44 | 4.25 |
| Ccl6 | NM_009139.2 | NM_009139.2:825 | 12.93 | 12.63 | 12.24 | 12.82 | 12.08 | 12.55 | 12.29 | 12.1 | 12.23 | 11.84 | 11.77 | 12.13 |
| Ccl7 | NM_013654.2 | NM_013654.2:215 | 8.2 | 7.79 | 7.81 | 7.39 | 7.3 | 7.33 | 7.87 | 8.64 | 8.45 | 6.4 | 7.55 | 7.1 |
| Ccl8 | NM_021443.2 | NM_021443.2:150 | 9.21 | 8.93 | 7.57 | 8.14 | 8.91 | 9.27 | 7.99 | 6.05 | 6.55 | 8.16 | 4.81 | 6.67 |
| Ccl9 | NM_011338.2 | NM_011338.2:1125 | 13.65 | 14 | 13.24 | 13.54 | 13.02 | 13.2 | 13.09 | 13.5 | 13.47 | 13.12 | 12.86 | 13.41 |
| Ccnd3 | NM_007632.2 | NM_007632.2:1311 | 10.44 | 11.48 | 10.55 | 9.08 | 9.75 | 9.9 | 9.49 | 9.43 | 9.96 | 10.3 | 10.99 | 12.31 |
| Ccr1 | NM_009912.4 | NM_009912.4:1526 | 8.02 | 8.99 | 8.35 | 8.58 | 7.91 | 8.67 | 7.86 | 8.56 | 8.32 | 10.23 | 10.07 | 10.92 |
| Ccr2 | NM_009915.2 | NM_009915.2:2965 | 9.21 | 9.04 | 10.04 | 9.14 | 8.89 | 9.6 | 9.04 | 8.96 | 10.36 | 7.92 | 7.85 | 8.22 |
| Ccr3 | NM_009914.4 | NM_009914.4:664 | 5.14 | 4.23 | 3.84 | 7.84 | 6.49 | 5.29 | 5.09 | 2.75 | 8.06 | 6.72 | 7.18 | 9.53 |
| Ccr4 | NM_009916.2 | NM_009916.2:394 | 0.39 | 0.08 | 2.13 | 0.11 | 0.08 | 0.7 | 0 | 0 | 2.51 | 1.43 | 4.98 | 3.32 |
| Ccr5 | NM_009917.5 | NM_009917.5:1340 | 11.53 | 11.04 | 11.18 | 10.18 | 10.46 | 10.82 | 10.57 | 10.87 | 11.63 | 9.68 | 8.79 | 9.33 |
| Ccr6 | NM_001190333.1 | NM_001190333.1:660 | 0.39 | 0.08 | 2.13 | 0.11 | 0.08 | 0.7 | 0 | 0 | 1.61 | 0 | 0.78 | 0.02 |
| Ccr7 | NM_007719.2 | NM_007719.2:755 | 6 | 6.88 | 2.13 | 4.12 | 5.92 | 5.99 | 2.73 | 2.57 | 4.39 | 4.82 | 0.78 | 4.92 |
| Ccr8 | NM_007720.2 | NM_007720.2:426 | 0.39 | 0.08 | 2.13 | 0.11 | 0.08 | 0.7 | 0 | 0 | 1.61 | 0 | 0.78 | 0.02 |
| Ccr9 | NM_009913.6 | NM_009913.6:820 | 0.39 | 0.08 | 2.13 | 0.11 | 0.08 | 0.7 | 0 | 0 | 1.61 | 0 | 0.78 | 0.02 |
| Ccrl2 | NM_017466.4 | NM_017466.4:655 | 12.49 | 12.12 | 11.6 | 11.04 | 11.51 | 11.45 | 11.57 | 11.75 | 12.01 | 14.15 | 13.85 | 15.32 |
| Cd14 | NM_009841.3 | NM_009841.3:235 | 11.17 | 11.35 | 11.07 | 8.73 | 10.37 | 9.7 | 10.2 | 10.72 | 11.52 | 12.15 | 12.3 | 13.48 |
| Cd160 | NM_001163496.1 | NM_001163496.1:416 | 0.39 | 3.03 | 2.13 | 0.11 | 0.08 | 0.7 | 0.73 | 6.56 | 1.61 | 0 | 0.78 | 0.02 |
| Cd163 | NM_053094.2 | NM_053094.2:3225 | 7.63 | 3.86 | 2.13 | 6.18 | 6.23 | 5.29 | 5.29 | 5.7 | 1.61 | 4.13 | 0.78 | 7.31 |
| Cd164 | NM_016898.2 | NM_016898.2:688 | 9.78 | 10.16 | 9.27 | 9.73 | 9.11 | 8.95 | 9.51 | 9.17 | 9.36 | 10.56 | 10.27 | 11.45 |
| Cd180 | NM_008533.2 | NM_008533.2:860 | 10.17 | 10.77 | 10.36 | 9.57 | 9.86 | 9.84 | 9.27 | 9.6 | 9.93 | 8.77 | 8.36 | 9.55 |
| Cd19 | NM_009844.2 | NM_009844.2:1118 | 0.39 | 0.08 | 2.13 | 1.76 | 3.68 | 0.7 | 0.32 | 0.25 | 1.61 | 0 | 0.78 | 0.02 |
| Cd1d1 | NM_007639.3 | NM_007639.3:1340 | 3.85 | 7.02 | 2.46 | 5.25 | 2.13 | 2.74 | 1.06 | 3.52 | 1.61 | 2.43 | 0.78 | 3.46 |
| Cd1d2 | NM_007640.2 | NM_007640.2:438 | 3.85 | 1.99 | 5.49 | 4.37 | 2.13 | 3.53 | 1.9 | 4.22 | 5.18 | 4.68 | 7.16 | 5.44 |
| Cd2 | XM_006500959.1 | XM_006500959.1:82 | 0.39 | 0.08 | 2.13 | 0.11 | 0.08 | 0.7 | 0 | 0 | 1.61 | 0 | 0.78 | 0.02 |
| Cd200 | NM_010818.3 | NM_010818.3:686 | 1.39 | 3.75 | 2.13 | 2.73 | 0.25 | 3.53 | 4.71 | 3.05 | 4.16 | 3.52 | 0.78 | 5.94 |
| Cd200r1 | NM_021325.3 | NM_021325.3:1090 | 8.04 | 8.38 | 8 | 8.67 | 8.5 | 8.64 | 8.1 | 6.18 | 8.06 | 6.87 | 7.83 | 6.12 |
| Cd207 | NM_144943.3 | NM_144943.3:605 | 0.39 | 0.08 | 2.13 | 2.94 | 0.08 | 0.7 | 0 | 2.12 | 1.61 | 1.01 | 0.78 | 0.02 |
| Cd22 | NM_001043317.2 | NM_001043317.2:865 | 9.5 | 8.25 | 6.96 | 8.32 | 8.75 | 8.47 | 9.12 | 5.32 | 7.98 | 4.98 | 0.78 | 7.77 |
| Cd244 | NM_018729.2 | NM_018729.2:262 | 0.39 | 0.08 | 2.13 | 0.11 | 0.08 | 0.7 | 0 | 0 | 1.61 | 0 | 0.78 | 0.02 |
| Cd247 | NM_001113391.2 | NM_001113391.2:215 | 0.39 | 0.08 | 2.13 | 0.11 | 0.08 | 0.7 | 0 | 0 | 1.61 | 0 | 0.78 | 6.91 |
| Cd27 | NM_001033126.2 | NM_001033126.2:235 | 2.39 | 0.08 | 3.84 | 0.11 | 4 | 3.31 | 2.54 | 3.31 | 3.13 | 3.43 | 0.78 | 4.96 |
| Cd274 | NM_021893.2 | NM_021893.2:515 | 8.37 | 7.34 | 8.23 | 6.97 | 7.07 | 7.9 | 6.36 | 7.08 | 8.03 | 8.16 | 8.74 | 9.48 |
| Cd276 | NM_133983.4 | NM_133983.4:1278 | 8.84 | 8.97 | 8.5 | 8.5 | 8.83 | 8.92 | 8.33 | 9.71 | 9.31 | 5.92 | 7.56 | 4.47 |
| Cd28 | NM_007642.4 | NM_007642.4:3304 | 4.91 | 3.21 | 2.13 | 4.29 | 3.42 | 0.87 | 5.71 | 6.4 | 3.13 | 3.13 | 0.78 | 2.8 |
| Cd33 | NM_001111058.1 | NM_001111058.1:405 | 7.76 | 7.22 | 7.1 | 6.93 | 7.18 | 5.75 | 6.71 | 7.1 | 1.61 | 9.8 | 9.91 | 10.65 |
| Cd34 | NM_001111059.1 | NM_001111059.1:560 | 0.39 | 0.08 | 2.13 | 0.11 | 0.08 | 0.7 | 0 | 0 | 1.61 | 0 | 0.78 | 0.02 |
| Cd36 | NM_007643.3 | NM_007643.3:1520 | 9.45 | 10.81 | 10.54 | 9.53 | 9.43 | 10.68 | 9.15 | 10.36 | 10.54 | 9.47 | 8.97 | 8.73 |
| Cd37 | NM_007645.4 | NM_007645.4:180 | 6.73 | 5.98 | 7.95 | 4.37 | 5.73 | 6.38 | 4.46 | 6.48 | 7.25 | 8.36 | 7.5 | 7.47 |
| Cd38 | NM_007646.4 | NM_007646.4:1490 | 12.46 | 12.51 | 11.53 | 12.79 | 12.13 | 11.98 | 11.96 | 11.55 | 11.38 | 11.32 | 9.91 | 11.49 |
| Cd3d | NM_013487.2 | NM_013487.2:289 | 0.39 | 0.08 | 2.13 | 0.11 | 0.08 | 0.7 | 0 | 0 | 1.61 | 6.95 | 6.72 | 6.6 |
| Cd3e | NM_007648.4 | NM_007648.4:380 | 0.39 | 0.08 | 2.13 | 0.11 | 0.08 | 2.74 | 0 | 0 | 1.61 | 4.6 | 0.78 | 0.02 |
| Cd3eap | NM_145822.2 | NM_145822.2:1858 | 4.78 | 4.97 | 5.18 | 2.16 | 4.18 | 3.89 | 4.8 | 4.39 | 6.18 | 5.84 | 0.78 | 4.53 |
| Cd3g | NM_009850.2 | NM_009850.2:430 | 0.39 | 0.08 | 2.13 | 0.11 | 0.08 | 0.7 | 0 | 0 | 6.46 | 2.01 | 0.78 | 3.81 |
| Cd4 | NM_013488.2 | NM_013488.2:950 | 0.39 | 0.08 | 2.13 | 5.4 | 4.99 | 0.7 | 0 | 5.17 | 1.61 | 0 | 0.78 | 0.02 |
| Cd40 | NM_011611.2 | NM_011611.2:1425 | 11.06 | 11.29 | 11.13 | 11.09 | 10.79 | 10.84 | 9.91 | 9.9 | 10.91 | 9.18 | 9.16 | 9.1 |
| Cd40lg | NM_011616.2 | NM_011616.2:600 | 3.56 | 3.86 | 2.13 | 2.16 | 0.08 | 3.89 | 3.13 | 1.83 | 1.61 | 4.18 | 0.78 | 4.71 |
| Cd44 | NM_009851.2 | NM_009851.2:112 | 4.2 | 6.69 | 2.13 | 0.11 | 4.09 | 3.72 | 4.76 | 5.17 | 5.4 | 3.34 | 2.53 | 4.53 |
| Cd46 | NM_010778.3 | NM_010778.3:115 | 0.39 | 0.08 | 2.13 | 0.11 | 0.08 | 0.7 | 0 | 0 | 1.61 | 0 | 0.78 | 0.02 |
| Cd47 | NM_010581.3 | NM_010581.3:165 | 10.14 | 10.61 | 10.22 | 9.97 | 10.04 | 10.5 | 9.81 | 10.6 | 10.66 | 10.19 | 10.52 | 11.02 |
| Cd48 | NM_007649.4 | NM_007649.4:370 | 9.58 | 9.28 | 9.62 | 10.45 | 9.75 | 9.85 | 9.06 | 10.46 | 9.92 | 8.6 | 6.99 | 9.34 |
| Cd5 | NM_007650.3 | NM_007650.3:1395 | 0.39 | 0.08 | 2.13 | 0.11 | 6.33 | 0.7 | 0 | 0 | 6.73 | 5.45 | 0.78 | 0.02 |
| Cd53 | NM_007651.3 | NM_007651.3:2300 | 6.17 | 3.51 | 7.35 | 6.97 | 5.33 | 7.75 | 5.45 | 6.06 | 5.99 | 6.95 | 7.41 | 9.28 |
| Cd55 | NM_010016.2 | NM_010016.2:1058 | 0.39 | 0.08 | 2.13 | 0.11 | 0.08 | 0.7 | 0 | 0 | 1.61 | 0 | 0.78 | 4.53 |

| | | | | | | | | | | | | | | |
|---|---|---|---|---|---|---|---|---|---|---|---|---|---|---|
| Cd59b | NM_181858.1 | NM_181858.1:322 | 0.39 | 0.08 | 2.13 | 0.11 | 0.08 | 0.7 | 0 | 0 | 1.61 | 2.6 | 0.78 | 0.02 |
| Cd6 | NM_001037801.2 | NM_001037801.2:1315 | 0.39 | 0.08 | 2.13 | 0.11 | 0.08 | 3.89 | 0 | 0 | 1.61 | 4.98 | 5.44 | 6.76 |
| Cd63 | NM_001042580.1 | NM_001042580.1:220 | 11.75 | 11.96 | 11.03 | 11.22 | 11.98 | 11.47 | 11.51 | 11.47 | 11.46 | 11.63 | 11.22 | 12.7 |
| Cd68 | NM_009853.1 | NM_009853.1:636 | 11.83 | 12.33 | 11.81 | 12.04 | 11.68 | 11.82 | 11.77 | 11.59 | 12.49 | 11.22 | 11.05 | 11.49 |
| Cd69 | NM_001033122.3 | NM_001033122.3:1370 | 9.92 | 10.19 | 9.86 | 10.29 | 9.92 | 9.7 | 8.95 | 10.07 | 9.71 | 10.21 | 10.63 | 10.98 |
| Cd7 | NM_009854.1 | NM_009854.1:234 | 0.39 | 2.33 | 3.84 | 0.11 | 0.08 | 0.7 | 0.73 | 0 | 3.13 | 3.52 | 2.53 | 4.33 |
| Cd70 | NM_011617.1 | NM_011617.1:650 | 0.39 | 0.08 | 2.13 | 2.94 | 0.08 | 0.7 | 0 | 0 | 1.61 | 4.43 | 7.64 | 0.02 |
| Cd74 | NM_001042605.1 | NM_001042605.1:391 | 12.01 | 12.41 | 12.62 | 11.31 | 12.05 | 12.14 | 8.48 | 9.43 | 10.54 | 8.63 | 8.41 | 8.24 |
| Cd79a | NM_007655.3 | NM_007655.3:1175 | 0.39 | 0.08 | 2.13 | 0.11 | 0.08 | 0.7 | 0 | 0 | 1.61 | 4.24 | 0.78 | 0.02 |
| Cd79b | NM_008339.2 | NM_008339.2:330 | 3.39 | 6.66 | 2.46 | 2.16 | 4.34 | 4.52 | 4.38 | 3.05 | 1.61 | 7.68 | 2.03 | 4.09 |
| Cd80 | NM_009855.2 | NM_009855.2:210 | 9.3 | 9.93 | 9.26 | 8.62 | 8.96 | 9.23 | 7.99 | 10.12 | 9.45 | 9.76 | 11.57 | 10.91 |
| Cd81 | NM_133655.2 | NM_133655.2:575 | 11.73 | 11.84 | 10.51 | 11.46 | 11.73 | 11.13 | 10.94 | 11.16 | 10.7 | 11.08 | 10.75 | 11.31 |
| Cd83 | NM_009856.2 | NM_009856.2:1624 | 12.31 | 12.6 | 12.34 | 12.26 | 11.9 | 12.04 | 11.68 | 12.1 | 12.41 | 10.91 | 10.47 | 11.17 |
| Cd84 | NM_001252472.1 | NM_001252472.1:188 | 12.01 | 12.1 | 11.48 | 11.71 | 11.51 | 11.75 | 11.5 | 11.73 | 11.73 | 11.42 | 11.68 | 12.06 |
| Cd86 | NM_019388.3 | NM_019388.3:251 | 10.53 | 11.09 | 10.02 | 10.05 | 10.1 | 10.08 | 9.08 | 10.25 | 10.08 | 8.32 | 9.28 | 9.61 |
| Cd8a | NM_001081110.2 | NM_001081110.2:355 | 0.39 | 0.08 | 2.13 | 0.11 | 0.08 | 0.7 | 0 | 0 | 1.61 | 0 | 0.78 | 0.02 |
| Cd8b1 | NM_009858.2 | NM_009858.2:1075 | 0.39 | 0.08 | 2.13 | 0.11 | 0.08 | 0.7 | 0 | 0 | 4.58 | 2.24 | 0.78 | 0.02 |
| Cd9 | NM_007657.3 | NM_007657.3:620 | 6.99 | 7.39 | 7.35 | 7.56 | 7.68 | 7.84 | 7.14 | 7.55 | 7.83 | 9.12 | 9.72 | 9.95 |
| Cd96 | NM_032465.2 | NM_032465.2:34 | 0.39 | 6.2 | 2.13 | 1.76 | 0.08 | 0.7 | 0 | 0 | 1.61 | 0 | 0.78 | 0.02 |
| Cd97 | NM_011925.1 | NM_011925.1:1975 | 12.32 | 12.42 | 11.89 | 11.59 | 11.67 | 11.54 | 11.45 | 11.9 | 12.05 | 12.88 | 12.82 | 13.71 |
| Cd99 | NM_025584.2 | NM_025584.2:716 | 10.01 | 9.11 | 8.8 | 7.59 | 9.12 | 9.74 | 9 | 9.67 | 8.44 | 7.97 | 8.3 | 8.96 |
| Cdh1 | NM_009864.2 | NM_009864.2:1610 | 7.34 | 7.44 | 7.01 | 5.61 | 7.52 | 7.02 | 0 | 6.33 | 6.77 | 4.52 | 0.78 | 5.22 |
| Cdh5 | NM_009868.3 | NM_009868.3:1615 | 1.39 | 3.03 | 2.13 | 0.11 | 0.08 | 0.7 | 0 | 0.98 | 1.61 | 6.37 | 0.78 | 0.02 |
| Cdk1 | NM_007659.3 | NM_007659.3:2538 | 0.39 | 0.08 | 2.13 | 0.28 | 0.08 | 0.7 | 0 | 0 | 1.61 | 0 | 0.78 | 0.02 |
| Cdkn1a | NM_007669.4 | NM_007669.4:1670 | 12.75 | 13.48 | 12.85 | 12.96 | 12.05 | 12.63 | 11.92 | 12.73 | 12.92 | 13.25 | 13.38 | 14.43 |
| Ceacam1 | NM_001039185.1 | NM_001039185.1:294 | 7.33 | 6.1 | 6.96 | 8.03 | 6.85 | 7.3 | 7.33 | 7.59 | 8.37 | 8.37 | 8.1 | 9.08 |
| Cebpb | NM_009883.3 | NM_009883.3:1147 | 10.32 | 10.13 | 10.14 | 10.23 | 9.31 | 9.4 | 9.3 | 9.65 | 10.42 | 11.44 | 11.74 | 12.55 |
| Cfb | NM_008198.2 | NM_008198.2:1685 | 9.43 | 8.44 | 8.84 | 7.91 | 8.81 | 8.52 | 8.24 | 8.62 | 8.53 | 8.27 | 7.68 | 8.64 |
| Cfd | NM_013459.1 | NM_013459.1:526 | 0.39 | 0.08 | 2.13 | 0.11 | 0.08 | 0.7 | 0 | 0 | 1.61 | 0 | 0.78 | 0.02 |
| Cfh | NM_009888.3 | NM_009888.3:807 | 8.81 | 7.31 | 6.9 | 7.96 | 7.7 | 7.79 | 7.21 | 6.82 | 7.16 | 7.07 | 0.78 | 6.2 |
| Cfi | NM_007686.2 | NM_007686.2:421 | 0.39 | 0.89 | 2.46 | 0.11 | 0.08 | 0.7 | 0.32 | 0 | 1.61 | 0 | 0.78 | 2.3 |
| Cfp | NM_008823.3 | NM_008823.3:1107 | 11.14 | 11.49 | 10.74 | 11.46 | 10.86 | 10.63 | 10.87 | 10.92 | 11.12 | 10.26 | 10 | 10.09 |
| Chil3 | NM_009892.2 | NM_009892.2:823 | 7.85 | 7.83 | 7.57 | 6.11 | 5.7 | 6.16 | 6.69 | 8.28 | 7.38 | 5.07 | 0.78 | 7.99 |
| Chit1 | NM_027979.1 | NM_027979.1:335 | 0.39 | 0.08 | 2.13 | 0.11 | 0.08 | 0.7 | 0 | 0 | 1.61 | 0 | 0.78 | 0.02 |
| Chuk | NM_001162410.1 | NM_001162410.1:222 | 5.03 | 5.24 | 5.63 | 6.67 | 7.67 | 6.87 | 7.21 | 7.64 | 3.56 | 6.4 | 6.11 | 5.66 |
| Ciita | NM_007575.2 | NM_007575.2:1358 | 4.48 | 3.75 | 5.34 | 3.93 | 3.8 | 4.52 | 2.82 | 3.52 | 4.76 | 3.13 | 4.62 | 2.3 |
| Cklf | NM_029295.2 | NM_029295.2:54 | 6.99 | 6.76 | 6.61 | 6.8 | 6.51 | 6.61 | 6.26 | 6.54 | 7.06 | 7.17 | 8.06 | 8.74 |
| Clec4a2 | NM_001170332.1 | NM_001170332.1:923 | 11.63 | 12.11 | 11.71 | 11.73 | 11.44 | 11.82 | 11.17 | 11.86 | 11.89 | 10.56 | 10.57 | 11 |
| Clec4n | NM_001190320.1 | NM_001190320.1:441 | 9.23 | 9.38 | 7.76 | 9.37 | 8.22 | 8.87 | 8.48 | 8.45 | 8.08 | 9.9 | 10.36 | 11.62 |
| Clec5a | NM_001038604.1 | NM_001038604.1:605 | 7.35 | 9.19 | 7.39 | 8.13 | 7.03 | 6.7 | 7.27 | 7.31 | 7.36 | 8.21 | 8.26 | 9.72 |
| Clec7a | NM_020008.2 | NM_020008.2:1008 | 10.31 | 10.08 | 10.55 | 9.73 | 9.89 | 10.06 | 9.15 | 8.69 | 9.71 | 10.95 | 11.44 | 11.99 |
| Clu | NM_013492.2 | NM_013492.2:354 | 0.39 | 0.08 | 2.13 | 0.11 | 0.08 | 0.7 | 0 | 2.91 | 4.91 | 3.89 | 0.78 | 0.02 |
| Cma1 | NM_010780.2 | NM_010780.2:860 | 0.39 | 0.08 | 2.13 | 4.87 | 0.08 | 0.7 | 0 | 0 | 1.61 | 0 | 5.6 | 0.02 |
| Cmah | NM_001111110.2 | NM_001111110.2:664 | 0.39 | 0.08 | 2.13 | 0.11 | 0.08 | 0.7 | 0 | 0 | 1.61 | 0 | 7.21 | 7.79 |
| Cmklr1 | NM_008153.3 | NM_008153.3:445 | 12.31 | 12.22 | 11.57 | 11.13 | 11.76 | 11.9 | 11.89 | 12.15 | 12.12 | 10.95 | 9.92 | 10.99 |
| Cmpk2 | NM_020557.4 | NM_020557.4:1600 | 9.6 | 10.15 | 9.42 | 8.76 | 9.13 | 9.71 | 8.49 | 9.61 | 10.43 | 9.58 | 8.72 | 10.07 |
| Col1a1 | NM_007742.3 | NM_007742.3:215 | 0.39 | 0.08 | 2.13 | 0.11 | 0.08 | 0.7 | 0 | 0 | 1.61 | 0 | 3.45 | 0.02 |
| Col3a1 | NM_009930.1 | NM_009930.1:4370 | 0.39 | 0.08 | 6.96 | 2.47 | 4.34 | 8.14 | 5.45 | 0 | 6.24 | 3.68 | 0.78 | 0.02 |
| Col4a1 | NM_009931.2 | NM_009931.2:4116 | 0.39 | 0.08 | 2.13 | 0.11 | 0.08 | 0.7 | 0.73 | 0 | 1.61 | 0.43 | 0.78 | 0.02 |
| Colec12 | NM_130449.2 | NM_130449.2:736 | 6.27 | 6.63 | 6.73 | 6.81 | 5.61 | 7.33 | 6.78 | 6.79 | 6.12 | 4.82 | 4.72 | 4.18 |
| Cr2 | NM_007758.2 | NM_007758.2:1650 | 0.39 | 1.54 | 2.13 | 0.11 | 1.73 | 0.7 | 0 | 0 | 1.61 | 0 | 0.78 | 0.02 |
| Creb1 | NM_001037726.1 | NM_001037726.1:2734 | 8.22 | 9.38 | 7.79 | 7.71 | 7.8 | 8.73 | 8.83 | 7.65 | 8.44 | 8.16 | 7.37 | 8.89 |
| Creb5 | NM_172728.2 | NM_172728.2:2415 | 5.34 | 8.53 | 4.22 | 7.22 | 5.73 | 7.14 | 5.67 | 3.87 | 3.13 | 6.24 | 0.78 | 5.68 |
| Crebbp | NM_001025432.1 | NM_001025432.1:3770 | 10.09 | 10.54 | 10.11 | 9.67 | 9.69 | 9.88 | 9.92 | 9.99 | 10.52 | 10.38 | 10.55 | 10.86 |
| Crp | NM_007768.4 | NM_007768.4:163 | 0.39 | 0.08 | 3.3 | 0.11 | 0.08 | 0.7 | 3.64 | 1.47 | 4.16 | 1.43 | 0.78 | 3.32 |
| Csf1 | NM_001113530.1 | NM_001113530.1:833 | 6.88 | 1.54 | 5.18 | 4.29 | 2.91 | 5.17 | 4.62 | 6.49 | 6.24 | 9.88 | 10.84 | 11.6 |
| Csf1r | NM_001037859.2 | NM_001037859.2:1892 | 9.15 | 9.99 | 9.29 | 9.72 | 8.87 | 8.92 | 9.01 | 8.86 | 9.49 | 8.87 | 6.99 | 9.47 |
| Csf2 | NM_009969.4 | NM_009969.4:732 | 0.39 | 5.2 | 2.13 | 0.11 | 0.08 | 0.7 | 0 | 0 | 5.18 | 0 | 0.78 | 3.46 |
| Csf2rb | NM_007780.4 | NM_007780.4:4185 | 11.9 | 12 | 12.21 | 11.67 | 11.26 | 11.84 | 10.46 | 11.3 | 11.89 | 12.45 | 13.01 | 13.72 |
| Csf3 | NM_009971.1 | NM_009971.1:830 | 0.39 | 0.08 | 2.13 | 0.11 | 0.08 | 0.7 | 0 | 0 | 1.61 | 0 | 0.78 | 1.97 |
| Csf3r | NM_001252651.1 | NM_001252651.1:1294 | 7.53 | 7.36 | 6.9 | 3.71 | 3.9 | 0.7 | 5.61 | 3.71 | 5.59 | 10.43 | 10.99 | 11.3 |
| Cspg4 | NM_139001.2 | NM_139001.2:1530 | 0.39 | 0.08 | 2.13 | 0.28 | 1.17 | 5.79 | 3.94 | 4.15 | 1.61 | 0 | 0.78 | 0.02 |
| Ctla4 | NM_009843.3 | NM_009843.3:1475 | 0.39 | 4.52 | 2.13 | 5.84 | 7.09 | 7.48 | 0 | 0 | 6.46 | 3.52 | 0.78 | 7.06 |
| Ctsg | NM_007800.1 | NM_007800.1:785 | 0.39 | 0.08 | 2.13 | 0.11 | 0.08 | 0.7 | 0 | 0 | 1.61 | 0 | 0.78 | 0.02 |
| Ctsh | NM_007801.2 | NM_007801.2:1090 | 10.7 | 11.17 | 10.65 | 11.22 | 10.7 | 10.53 | 9.91 | 10.46 | 10.81 | 9.74 | 9.17 | 9.85 |
| Ctsl | NM_009984.3 | NM_009984.3:45 | 7.87 | 9.39 | 8.78 | 8.06 | 7.78 | 8.83 | 8.25 | 8.3 | 9.74 | 7.16 | 7.91 | 7.78 |
| Ctss | NM_021281.2 | NM_021281.2:740 | 14.03 | 14.3 | 13.7 | 13.92 | 13.78 | 13.8 | 13.42 | 13.85 | 13.85 | 13.6 | 13.52 | 14.29 |
| Ctsw | NM_009985.4 | NM_009985.4:190 | 2.71 | 0.08 | 5.34 | 0.11 | 3.56 | 1.79 | 0.73 | 0.98 | 3.13 | 0 | 0.78 | 0.02 |
| Cx3cl1 | NM_009142.3 | NM_009142.3:125 | 5.43 | 6.07 | 6.32 | 6.09 | 4 | 5.83 | 3.59 | 4.93 | 5.99 | 6.21 | 7.23 | 7.43 |
| Cx3cr1 | NM_009987.3 | NM_009987.3:2696 | 10.78 | 11.08 | 10.31 | 11.98 | 11.33 | 10.89 | 10.8 | 10.95 | 11.16 | 10.03 | 8.1 | 9.27 |
| Cxcl1 | NM_008176.1 | NM_008176.1:560 | 12.54 | 12.85 | 12.19 | 12.09 | 11.99 | 12.09 | 11.97 | 12.51 | 12.79 | 12.61 | 12.71 | 13.85 |

| | | | | | | | | | | | | | | |
|---|---|---|---|---|---|---|---|---|---|---|---|---|---|---|
| Cxcl10 | NM_021274.1 | NM_021274.1:115 | 8.95 | 9.57 | 8.21 | 7.41 | 8.99 | 8.32 | 7.53 | 8.91 | 8.32 | 7.89 | 7.32 | 8.26 |
| Cxcl11 | NM_019494.1 | NM_019494.1:345 | 7.53 | 6.75 | 7.35 | 5.7 | 4.09 | 3.72 | 4.8 | 4.64 | 6.81 | 5.82 | 0.78 | 5.26 |
| Cxcl12 | NM_021704.3 | NM_021704.3:259 | 0.39 | 0.08 | 2.13 | 0.11 | 0.08 | 0.7 | 0 | 0 | 1.61 | 0 | 0.78 | 0.02 |
| Cxcl13 | NM_018866.2 | NM_018866.2:551 | 0.39 | 0.08 | 2.13 | 0.11 | 0.25 | 0.7 | 0 | 0 | 1.61 | 0 | 0.78 | 0.02 |
| Cxcl14 | XM_006517307.1 | XM_006517307.1:446 | 6.5 | 6.24 | 6.47 | 7.15 | 6.46 | 6.4 | 4.86 | 6.34 | 6.55 | 4.75 | 6.04 | 2.57 |
| Cxcl15 | NM_011339.2 | NM_011339.2:419 | 0.39 | 0.08 | 2.13 | 0.11 | 0.08 | 0.7 | 0 | 0 | 1.61 | 0 | 0.78 | 6.46 |
| Cxcl16 | NM_023158.6 | NM_023158.6:679 | 10.52 | 11.59 | 9.97 | 10.22 | 10.38 | 10.15 | 10.22 | 10.35 | 10.47 | 9.42 | 8.91 | 8.59 |
| Cxcl2 | NM_009140.2 | NM_009140.2:765 | 14.12 | 14.38 | 13.68 | 14.05 | 13.19 | 13.5 | 13.49 | 13.87 | 14 | 15.12 | 15.1 | 16.36 |
| Cxcl3 | NM_203320.2 | NM_203320.2:275 | 0.39 | 5.16 | 2.13 | 4.37 | 0.08 | 0.7 | 5.75 | 2.12 | 4.39 | 8.38 | 7.26 | 9.69 |
| Cxcl5 | NM_009141.2 | NM_009141.2:565 | 0.39 | 0.08 | 2.13 | 4.7 | 0.08 | 0.7 | 0 | 7.04 | 4.58 | 0 | 0.78 | 0.02 |
| Cxcl9 | NM_008599.2 | NM_008599.2:40 | 5.39 | 4.52 | 7.6 | 5.33 | 2.44 | 6.91 | 1.06 | 1.83 | 5.18 | 0 | 0.78 | 2.57 |
| Cxcr1 | NM_178241.4 | NM_178241.4:860 | 0.39 | 0.08 | 2.13 | 3.93 | 0.08 | 0.7 | 0 | 0 | 1.61 | 6.29 | 5.38 | 7.93 |
| Cxcr2 | NM_009909.3 | NM_009909.3:440 | 4.71 | 3.21 | 2.13 | 0.28 | 0.08 | 0.7 | 1.9 | 0.25 | 1.61 | 9.97 | 10.58 | 11.43 |
| Cxcr3 | NM_009910.2 | NM_009910.2:605 | 0.39 | 0.08 | 2.13 | 0.11 | 0.08 | 0.7 | 0 | 0 | 1.61 | 0 | 0.78 | 0.02 |
| Cxcr4 | NM_009911.3 | NM_009911.3:704 | 12 | 12.32 | 12.12 | 11.18 | 11.84 | 11.54 | 11.23 | 12.05 | 12.76 | 12.42 | 12.76 | 13.44 |
| Cxcr5 | NM_007551.2 | NM_007551.2:1648 | 0.39 | 0.08 | 2.13 | 0.11 | 0.08 | 0.7 | 0 | 0 | 1.61 | 0 | 0.78 | 1.54 |
| Cxcr6 | NM_030712.4 | NM_030712.4:650 | 0.39 | 5.32 | 2.13 | 3.13 | 0.08 | 5.83 | 3.38 | 3.42 | 7.03 | 6.91 | 0.78 | 0.02 |
| Cybb | NM_007807.2 | NM_007807.2:1535 | 11.08 | 10.9 | 10.76 | 10.55 | 10.9 | 11.02 | 10.35 | 10.48 | 11.33 | 9.88 | 9.96 | 10.58 |
| Cyfip2 | NM_001252459.1 | NM_001252459.1:1136 | 3.97 | 5.11 | 6.32 | 5.25 | 6.37 | 5.71 | 4.59 | 6.22 | 7.83 | 7.16 | 6.72 | 6.86 |
| Cyld | NM_001276279.1 | NM_001276279.1:2200 | 6.71 | 6.41 | 6.79 | 6.34 | 7.33 | 7.4 | 6.43 | 6.28 | 6.46 | 7.05 | 7.01 | 8.47 |
| Ddx58 | NM_172689.3 | NM_172689.3:1751 | 10.47 | 11.2 | 11.09 | 10.46 | 10.42 | 9.87 | 9.17 | 11.19 | 10.56 | 10.24 | 12.06 | 10.58 |
| Ddx60 | NM_001081215.1 | NM_001081215.1:2965 | 9.05 | 9.56 | 9.6 | 8.88 | 9.23 | 9.2 | 8.34 | 9.55 | 9.63 | 8.7 | 9.31 | 7.74 |
| Defb1 | NM_007843.3 | NM_007843.3:157 | 2.39 | 0.08 | 2.13 | 1.76 | 1.17 | 4.18 | 0.32 | 3.19 | 4.91 | 5.95 | 3.45 | 0.02 |
| Dll4 | NM_019454.2 | NM_019454.2:542 | 0.39 | 0.08 | 2.13 | 0.11 | 0.08 | 0.7 | 0 | 0 | 1.61 | 4.18 | 0.78 | 4.4 |
| Dmbt1 | XM_006507298.1 | XM_006507298.1:2054 | 0.39 | 3.75 | 2.13 | 2.47 | 1.73 | 0.7 | 1.06 | 2.36 | 1.61 | 0 | 0.78 | 0.02 |
| Dock9 | XM_976068.1 | XM_976068.1:360 | 4.2 | 0.08 | 8.07 | 6.7 | 4.9 | 5.41 | 4.09 | 4.15 | 7.93 | 8.08 | 6.79 | 8.85 |
| Dpp4 | NM_001159543.1 | NM_001159543.1:1303 | 7.81 | 7.71 | 8 | 8.96 | 8.6 | 8.18 | 0 | 0 | 7.27 | 1.01 | 8.06 | 1.97 |
| Dusp4 | NM_176933.4 | NM_176933.4:2200 | 7.03 | 8.53 | 7.15 | 7.15 | 5.7 | 7.16 | 7.16 | 8.22 | 9.05 | 8.59 | 9.11 | 8.3 |
| Dusp6 | NM_026268.2 | NM_026268.2:1648 | 7.43 | 5.28 | 6.24 | 6.4 | 6.97 | 6.16 | 5.87 | 6.63 | 7.36 | 6.65 | 5.7 | 7.26 |
| Ebi3 | NM_015766.2 | NM_015766.2:1015 | 9.45 | 9.69 | 8.09 | 9.91 | 8.23 | 9.07 | 8.17 | 9.46 | 7.83 | 9.6 | 10.97 | 11.79 |
| Ecsit | NM_001253897.1 | NM_001253897.1:348 | 0.39 | 0.08 | 3.3 | 0.11 | 0.08 | 0.7 | 0 | 0 | 1.61 | 4.95 | 5.2 | 4.82 |
| Egfr | NM_207655.2 | NM_207655.2:1335 | 0.39 | 0.08 | 2.13 | 0.11 | 0.08 | 0.7 | 4.64 | 0 | 1.61 | 5.21 | 0.78 | 0.02 |
| Egr1 | NM_007913.5 | NM_007913.5:515 | 11.44 | 12.51 | 11.63 | 11.76 | 11.53 | 11.1 | 10.37 | 11.75 | 11.36 | 13.18 | 13.31 | 14.7 |
| Egr2 | NM_010118.2 | NM_010118.2:1785 | 0.39 | 0.08 | 2.13 | 0.11 | 0.08 | 0.7 | 0 | 0 | 1.61 | 0 | 0.78 | 4.4 |
| Egr3 | NM_018781.2 | NM_018781.2:518 | 2.39 | 6.87 | 2.13 | 2.94 | 5.94 | 0.7 | 3.19 | 2.91 | 1.61 | 7.79 | 8.32 | 9.07 |
| Elane | NM_015779.2 | NM_015779.2:491 | 0.39 | 0.08 | 2.13 | 0.11 | 0.08 | 0.7 | 0 | 0 | 1.61 | 0 | 0.78 | 0.02 |
| Elk1 | NM_007922.4 | NM_007922.4:3070 | 3.97 | 6.12 | 5.18 | 3.13 | 5.92 | 4.41 | 5.42 | 5.84 | 5.99 | 5.31 | 6.25 | 5.33 |
| Emr1 | NM_010130.1 | NM_010130.1:995 | 10.99 | 10.85 | 11.07 | 10.83 | 11.01 | 10.8 | 10.91 | 10.87 | 11.83 | 9.55 | 9.21 | 9.37 |
| Eng | NM_001146350.1 | NM_001146350.1:574 | 4.48 | 0.08 | 2.13 | 5.73 | 0.08 | 4.71 | 5 | 2.57 | 1.61 | 4.64 | 0.78 | 0.02 |
| Entpd1 | NM_009848.3 | NM_009848.3:2062 | 8.14 | 7.94 | 9.16 | 9.37 | 8.88 | 8.83 | 6.79 | 6.91 | 7.75 | 9.14 | 9.69 | 10.56 |
| Eomes | NM_010136.2 | NM_010136.2:2665 | 2.97 | 0.08 | 4.78 | 4.12 | 0.08 | 0.7 | 1.73 | 1.47 | 1.61 | 1.43 | 0.78 | 0.02 |
| Ep300 | NM_177821.6 | NM_177821.6:4305 | 8.07 | 8.21 | 7.67 | 6.88 | 7.9 | 7.63 | 7.34 | 7.29 | 8.08 | 8.23 | 8.12 | 8.11 |
| Epcam | NM_008532.2 | NM_008532.2:1550 | 5.43 | 7.68 | 2.13 | 6.3 | 7 | 7.37 | 1.73 | 4.64 | 1.61 | 7.54 | 2.53 | 6.79 |
| Epsti1 | NM_029495.2 | NM_029495.2:26 | 0.39 | 0.08 | 2.13 | 2.73 | 0.25 | 0.7 | 0 | 0.25 | 4.16 | 0 | 0.78 | 0.02 |
| Erbb2 | NM_001003817.1 | NM_001003817.1:1395 | 0.39 | 0.08 | 2.13 | 0.11 | 0.08 | 0.7 | 0.73 | 0 | 1.61 | 4.95 | 4.62 | 0.02 |
| Ets1 | NM_001038642.1 | NM_001038642.1:740 | 7.17 | 9.85 | 8.6 | 7.8 | 7.64 | 8.45 | 8.47 | 8.91 | 8.77 | 6.98 | 9.03 | 7.28 |
| Ewsr1 | NM_007968.3 | NM_007968.3:150 | 11.32 | 11.41 | 11.05 | 10.6 | 11.23 | 10.83 | 10.79 | 11.58 | 11.21 | 11.35 | 11.73 | 11.84 |
| F12 | NM_021489.2 | NM_021489.2:405 | 0.39 | 0.08 | 2.13 | 0.11 | 0.08 | 0.7 | 0 | 0 | 1.61 | 2.6 | 0.78 | 0.02 |
| F13a1 | NM_001166391.1 | NM_001166391.1:2790 | 11.52 | 11.34 | 10.83 | 11.25 | 11.32 | 10.85 | 11.32 | 11.43 | 11.26 | 10.74 | 10.03 | 11.1 |
| F2rl1 | NM_007974.4 | NM_007974.4:994 | 4.09 | 0.89 | 6.16 | 3.71 | 4.61 | 5.29 | 0 | 2.75 | 5.05 | 3.89 | 0.78 | 2.3 |
| Fadd | NM_010175.5 | NM_010175.5:2641 | 7.89 | 7.96 | 7.7 | 7.2 | 7.26 | 8.58 | 7.67 | 7.65 | 8.05 | 7.96 | 6.43 | 5.1 |
| Fap | NM_007986.2 | NM_007986.2:530 | 4.97 | 4.31 | 3.3 | 5.29 | 4.26 | 4.52 | 3.44 | 3.52 | 1.61 | 2.24 | 0.78 | 0.02 |
| Fas | NM_007987.2 | NM_007987.2:596 | 7.07 | 7.14 | 6.54 | 6.09 | 6.62 | 5.46 | 6.54 | 6.05 | 7.71 | 8.19 | 8.86 | 9.41 |
| Fasl | NM_010177.3 | NM_010177.3:645 | 0.39 | 0.08 | 2.13 | 0.11 | 4.18 | 0.7 | 0 | 0 | 1.61 | 2.6 | 0.78 | 0.02 |
| Fcer1a | NM_010184.1 | NM_010184.1:114 | 0.39 | 0.89 | 2.13 | 0.11 | 0.08 | 0.7 | 1.9 | 0 | 1.61 | 0.43 | 0.78 | 0.02 |
| Fcer1g | NM_010185.4 | NM_010185.4:264 | 10.32 | 10.82 | 10.35 | 10.45 | 10.03 | 10.08 | 9.84 | 10.27 | 10.57 | 10.26 | 10.32 | 11.08 |
| Fcer2a | NM_001253737.1 | NM_001253737.1:1770 | 1.97 | 1.54 | 4.78 | 3.13 | 4.18 | 4.41 | 0 | 1.83 | 1.61 | 3.13 | 4.52 | 5.26 |
| Fcgr1 | NM_010186.5 | NM_010186.5:185 | 9.24 | 9.28 | 9.68 | 7.45 | 9.21 | 8.92 | 9.44 | 9.89 | 10.49 | 7.57 | 7.99 | 8.12 |
| Fcgr2b | NM_001077189.1 | NM_001077189.1:1225 | 12.79 | 13.37 | 13.31 | 13.27 | 12.95 | 13.12 | 11.46 | 12.76 | 13.1 | 11.57 | 11.58 | 12.34 |
| Fcgr3 | NM_010188.5 | NM_010188.5:1175 | 10.49 | 10.7 | 10.14 | 10.84 | 9.94 | 10.02 | 9.76 | 9.84 | 10.31 | 10.19 | 9.93 | 10.93 |
| Fcgr4 | NM_144559.1 | NM_144559.1:608 | 9.02 | 8.71 | 9.61 | 9.08 | 8.76 | 8.98 | 8.58 | 8.15 | 9.3 | 8.1 | 8.23 | 8.89 |
| Fez1 | NM_183171.4 | NM_183171.4:426 | 0.39 | 1.99 | 2.13 | 2.94 | 0.08 | 3.31 | 0 | 0 | 1.61 | 0 | 0.78 | 0.02 |
| Flt3 | NM_010229.2 | NM_010229.2:3302 | 0.39 | 4.77 | 2.13 | 1.76 | 4 | 0.7 | 0 | 0 | 1.61 | 0 | 0.78 | 1.97 |
| Flt3l | NM_013520.2 | NM_013520.2:529 | 0.39 | 0.08 | 2.13 | 0.11 | 0.25 | 0.7 | 0 | 3.95 | 1.61 | 2.6 | 0.78 | 0.02 |
| Fn1 | NM_010233.1 | NM_010233.1:2627 | 13.47 | 13.91 | 13.51 | 13.28 | 13.47 | 13.32 | 13.2 | 13.81 | 13.95 | 12.8 | 13.33 | 13.53 |
| Fos | NM_010234.2 | NM_010234.2:1330 | 13.52 | 13.62 | 13.49 | 13.46 | 12.95 | 13.32 | 12.43 | 13.29 | 13.6 | 13.21 | 13.68 | 14.1 |
| Foxj1 | NM_008240.3 | NM_008240.3:2179 | 0.39 | 0.08 | 2.13 | 4.03 | 0.08 | 0.7 | 0 | 0 | 1.61 | 5.16 | 0.78 | 3.32 |
| Foxp3 | NM_054039.2 | NM_054039.2:194 | 0.39 | 0.08 | 2.13 | 0.11 | 0.08 | 0.7 | 0 | 0 | 1.61 | 0 | 0.78 | 0.02 |
| Fpr2 | NM_008039.2 | NM_008039.2:1015 | 12.46 | 12.38 | 12.01 | 11.75 | 10.69 | 11.29 | 11.16 | 12.02 | 12.03 | 13.84 | 14.02 | 15.1 |
| Fut7 | NM_001177367.1 | NM_001177367.1:1151 | 0.39 | 0.08 | 2.13 | 0.11 | 0.08 | 0.7 | 0 | 0 | 1.61 | 2.43 | 0.78 | 2.8 |
| Fyn | NM_008054.2 | NM_008054.2:1030 | 8.16 | 7.98 | 8.44 | 8.76 | 8.74 | 8.38 | 7.92 | 7.02 | 8.28 | 7.63 | 8.33 | 5.99 |

| | | | | | | | | | | | | | | |
|---|---|---|---|---|---|---|---|---|---|---|---|---|---|---|
| Gata3 | NM_008091.3 | NM_008091.3:1943 | 2.39 | 1.99 | 2.13 | 5.82 | 4.09 | 2.34 | 3.44 | 2.57 | 4.58 | 6.17 | 4.15 | 4.25 |
| Gbp2b | NM_010259.2 | NM_010259.2:1808 | 7.97 | 7.85 | 8.16 | 6.8 | 4.41 | 7.86 | 4.38 | 5.53 | 5.77 | 5.91 | 3.2 | 6.35 |
| Gbp5 | NM_153564.2 | NM_153564.2:1176 | 0.39 | 8.34 | 2.13 | 0.11 | 0.08 | 0.7 | 5.86 | 0 | 1.61 | 6.16 | 5.32 | 7.74 |
| Gfi1 | NM_010278.2 | NM_010278.2:1875 | 0.39 | 0.08 | 2.13 | 0.11 | 0.08 | 0.7 | 0.32 | 0.25 | 1.61 | 3.24 | 0.78 | 0.02 |
| Glycam1 | NM_008134.2 | NM_008134.2:124 | 1.97 | 3.63 | 5.75 | 0.28 | 2.13 | 5.1 | 0.32 | 3.05 | 3.89 | 2.24 | 5.55 | 3.46 |
| Gpi1 | NM_008155.4 | NM_008155.4:1540 | 11.26 | 11.6 | 11.6 | 11.3 | 11.25 | 11.02 | 10.83 | 11.12 | 11.49 | 11.34 | 11.03 | 11.61 |
| Gpr183 | NM_183031.2 | NM_183031.2:238 | 9.43 | 9.78 | 10.57 | 9.7 | 9.59 | 10.25 | 8.62 | 8.77 | 10.04 | 7.92 | 8.53 | 9.19 |
| Gpr44 | XM_006526695.1 | XM_006526695.1:358 | 0.39 | 1.99 | 2.13 | 0.11 | 4.09 | 0.7 | 0 | 2.12 | 1.61 | 1.01 | 0.78 | 0.92 |
| Gtf3c1 | NM_207239.1 | NM_207239.1:4586 | 4.56 | 5.91 | 2.46 | 4.51 | 5.44 | 4.04 | 2.19 | 4.97 | 5.59 | 5.24 | 0.78 | 0.02 |
| Gzma | NM_010370.2 | NM_010370.2:188 | 1.97 | 3.03 | 2.13 | 2.94 | 3.1 | 0.7 | 2.44 | 0 | 3.13 | 0 | 0.78 | 0.02 |
| Gzmb | NM_013542.2 | NM_013542.2:1020 | 0.39 | 4.45 | 6.24 | 6.87 | 4.95 | 4.62 | 0 | 2.75 | 3.56 | 2.75 | 0.78 | 7.22 |
| Gzmk | NM_008196.1 | NM_008196.1:302 | 4.09 | 3.75 | 2.13 | 5.33 | 0.08 | 3.31 | 2.44 | 3.95 | 5.92 | 1.01 | 0.78 | 0.02 |
| Gzmm | NM_008504.2 | NM_008504.2:610 | 0.39 | 0.08 | 2.13 | 0.11 | 0.08 | 0.7 | 0 | 0 | 1.61 | 4.47 | 0.78 | 6.64 |
| H2-Aa | NM_010378.2 | NM_010378.2:450 | 13.01 | 13.2 | 12.91 | 11.69 | 12.85 | 13.04 | 8.69 | 9.42 | 10.42 | 8.97 | 9.47 | 8.8 |
| H2-Ab1 | NM_207105.2 | NM_207105.2:164 | 9.12 | 9.74 | 8.77 | 6.97 | 9.83 | 8.9 | 4.13 | 1.83 | 1.61 | 7.24 | 0.78 | 0.02 |
| H2-D1 | NM_010380.3 | NM_010380.3:1133 | 14.16 | 14.26 | 13.99 | 13.96 | 13.69 | 13.9 | 13.51 | 13.81 | 14.03 | 14 | 14.04 | 14.9 |
| H2-DMb1 | NM_010387.2 | NM_010387.2:1089 | 5.94 | 5.66 | 6.07 | 6.07 | 6.31 | 7.11 | 4.19 | 5.35 | 2.51 | 4.47 | 4.9 | 3.16 |
| H2-DMb2 | NM_010388.4 | NM_010388.4:195 | 10.09 | 10.1 | 9.68 | 9.27 | 10.52 | 10.2 | 5.63 | 6.24 | 6.85 | 4.72 | 0.78 | 5.63 |
| H2-Dma | NM_010386.3 | NM_010386.3:530 | 9.82 | 10.5 | 9.89 | 8.58 | 10.58 | 10.38 | 8.17 | 8.99 | 9.79 | 6.77 | 4.9 | 5.89 |
| H2-Ea-ps | NM_010381.2 | NM_010381.2:735 | 11.84 | 12 | 11.6 | 12.17 | 11.66 | 11.99 | 6.24 | 3.62 | 8.16 | 8.32 | 7.4 | 4.09 |
| H2-Eb1 | NM_010382.2 | NM_010382.2:935 | 8.42 | 8.51 | 9.72 | 9.26 | 8.93 | 9.58 | 0 | 4.93 | 5.77 | 7.14 | 7.74 | 10.24 |
| H2-K1 | NM_001001892.2 | NM_001001892.2:1370 | 14.49 | 14.62 | 14.3 | 14.39 | 13.85 | 14.19 | 13.95 | 14.09 | 14.43 | 14.6 | 14.55 | 15.54 |
| H2-M3 | NM_013819.2 | NM_013819.2:1180 | 13.15 | 13.33 | 13.18 | 13.07 | 13.06 | 13.09 | 12.59 | 12.92 | 13.13 | 11.78 | 11.97 | 11.91 |
| H2-Ob | NM_010389.3 | NM_010389.3:1606 | 7.79 | 8.19 | 8.25 | 7.11 | 7.7 | 8.32 | 1.06 | 0.25 | 4.91 | 3.13 | 0.78 | 7.31 |
| H2-Q1 | NM_010390.3 | NM_010390.3:316 | 0.39 | 0.08 | 2.13 | 0.11 | 0.08 | 0.7 | 0 | 0 | 1.61 | 4.64 | 0.78 | 0.02 |
| H2-Q10 | NM_010391.4 | NM_010391.4:890 | 0.39 | 0.08 | 2.13 | 0.11 | 0.08 | 0.7 | 0 | 0 | 1.61 | 3.82 | 0.78 | 0.02 |
| H2-Q2 | NM_010392.2 | NM_010392.2:472 | 0.39 | 0.08 | 2.13 | 0.11 | 0.08 | 0.7 | 0 | 0 | 1.61 | 0 | 0.78 | 0.02 |
| H2-T23 | NM_010398.3 | NM_010398.3:441 | 7.68 | 9.71 | 10.04 | 8.7 | 8.91 | 8.16 | 7.36 | 6.81 | 9.06 | 7.85 | 5.2 | 9.79 |
| H60a | NM_010400.2 | NM_010400.2:2020 | 11.11 | 10.98 | 10.75 | 11.22 | 10.18 | 10.94 | 10.07 | 10.95 | 11.02 | 10.86 | 10.08 | 11.95 |
| Hamp | NM_032541.1 | NM_032541.1:202 | 0.39 | 0.08 | 2.13 | 0.11 | 0.08 | 0.7 | 0 | 0 | 1.61 | 0 | 5.7 | 0.02 |
| Havcr2 | NM_134250.2 | NM_134250.2:134 | 9.26 | 10.01 | 9.93 | 8.54 | 9.31 | 9.5 | 7.34 | 9.92 | 9.32 | 8.49 | 9.45 | 10.12 |
| Hc | NM_010406.1 | NM_010406.1:1065 | 0.39 | 0.08 | 2.13 | 0.11 | 0.08 | 0.7 | 0 | 0 | 1.61 | 0 | 0.78 | 0.02 |
| Hck | NM_010407.3 | NM_010407.3:1874 | 13.66 | 13.8 | 13.38 | 13.94 | 13.1 | 14.08 | 13.51 | 13.58 | 12.94 | 13.26 | 14.14 | 14.32 |
| Hcst | NM_011827.3 | NM_011827.3:166 | 6.12 | 5.53 | 4.53 | 5.47 | 4.73 | 4.18 | 2.9 | 6.03 | 6.55 | 6.17 | 6.28 | 7.6 |
| Herc6 | NM_025992.2 | NM_025992.2:876 | 9.63 | 10.53 | 9.96 | 9.55 | 9.11 | 9.52 | 8.95 | 9.33 | 10.76 | 9.62 | 10.33 | 9.97 |
| Hif1a | NM_010431.2 | NM_010431.2:1294 | 10.77 | 10.79 | 10.8 | 10.5 | 9.85 | 10.32 | 10.16 | 10.87 | 11.22 | 10.83 | 10.93 | 11.77 |
| Hmgb1 | NM_010439.3 | NM_010439.3:1574 | 9.92 | 9.72 | 9.54 | 9.25 | 9.68 | 9.99 | 9.63 | 9.43 | 9.68 | 9.35 | 10.05 | 9.91 |
| Hras | NM_008284.2 | NM_008284.2:1890 | 0.39 | 0.08 | 2.13 | 0.11 | 0.08 | 0.7 | 0 | 0 | 1.61 | 0 | 0.78 | 0.02 |
| Hsd11b1 | NM_008288.2 | NM_008288.2:110 | 8.32 | 8.03 | 8.02 | 6.67 | 7.67 | 7.06 | 4.29 | 5.46 | 7.85 | 11.02 | 10.68 | 11.07 |
| Hspb2 | NM_024441.3 | NM_024441.3:667 | 0.39 | 0.08 | 2.13 | 0.11 | 0.08 | 0.7 | 0 | 0 | 1.61 | 0.43 | 0.78 | 0.02 |
| Icam1 | NM_010493.2 | NM_010493.2:2195 | 12.86 | 12.95 | 12.49 | 12.64 | 12.35 | 12.15 | 11.76 | 12.63 | 12.3 | 13.59 | 13.7 | 15.06 |
| Icam2 | NM_010494.1 | NM_010494.1:375 | 0.39 | 3.51 | 2.13 | 4.37 | 5.51 | 3.72 | 0 | 6.05 | 5.29 | 5.18 | 0.78 | 3.46 |
| Icam4 | NM_023892.2 | NM_023892.2:540 | 4.56 | 5.72 | 3.84 | 5.02 | 5.22 | 4.96 | 4.16 | 3.71 | 1.61 | 4.18 | 5.55 | 7.2 |
| Icos | NM_017480.1 | NM_017480.1:142 | 5.25 | 4.06 | 7.7 | 6.16 | 6.78 | 6.29 | 0.32 | 5.2 | 6.99 | 0.43 | 4.01 | 0.02 |
| Icosl | NM_015790.3 | NM_015790.3:349 | 8.92 | 8.47 | 8.81 | 7.57 | 8.51 | 7.73 | 6.85 | 8.24 | 8.73 | 7.86 | 9.48 | 9.03 |
| Ido1 | NM_008324.1 | NM_008324.1:521 | 2.97 | 3.36 | 6.61 | 1.76 | 4.18 | 3.72 | 2.73 | 4.09 | 3.89 | 3.82 | 5.7 | 4.47 |
| Ifi27 | NM_026790.2 | NM_026790.2:166 | 7.48 | 7.93 | 2.13 | 7.21 | 7.48 | 6.89 | 6.24 | 7.64 | 8.67 | 8.63 | 7.26 | 7.77 |
| Ifi35 | NM_027320.4 | NM_027320.4:820 | 10.36 | 10.35 | 10.6 | 9.9 | 10.4 | 10.12 | 9.71 | 10.79 | 11.86 | 10.19 | 10.22 | 10.29 |
| Ifi44 | NM_133871.2 | NM_133871.2:990 | 3.71 | 1.54 | 7.1 | 5.4 | 5.67 | 5.29 | 3.69 | 3.87 | 7.3 | 4.38 | 0.78 | 5.77 |
| Ifi44l | NM_031367.1 | NM_031367.1:1410 | 0.39 | 0.08 | 2.13 | 0.11 | 0.08 | 0.7 | 0 | 0 | 1.61 | 0 | 0.78 | 0.02 |
| Ifih1 | NM_027835.2 | NM_027835.2:1997 | 9.02 | 9.17 | 9.96 | 8.87 | 8.86 | 9.04 | 8.21 | 8.77 | 9.68 | 7.52 | 7.82 | 8.44 |
| Ifit1 | NM_008331.2 | NM_008331.2:890 | 4.85 | 7.4 | 5.49 | 5.02 | 6.39 | 5.75 | 4.23 | 4.73 | 5.18 | 5.95 | 4.15 | 5.5 |
| Ifit2 | NM_008332.2 | NM_008332.2:230 | 7.95 | 8.2 | 7.6 | 6.62 | 7.83 | 7.61 | 6.14 | 7.83 | 8.28 | 7.18 | 8.77 | 7.4 |
| Ifit3 | NM_010501.1 | NM_010501.1:1290 | 9.84 | 10.61 | 10.53 | 9.69 | 9.75 | 9.67 | 8.95 | 9.92 | 10.41 | 8.94 | 9.47 | 9.72 |
| Ifitm1 | NM_001112715.1 | NM_001112715.1:412 | 10.04 | 10.42 | 9.24 | 9.11 | 9.64 | 9.85 | 8.13 | 8.56 | 8.08 | 13.44 | 13.72 | 14.31 |
| Ifitm2 | NM_030694.1 | NM_030694.1:87 | 11.35 | 11.34 | 10.85 | 8.77 | 10.68 | 10.18 | 11.13 | 10.79 | 11.88 | 11.82 | 11.92 | 12.65 |
| Ifna1 | NM_010502.2 | NM_010502.2:30 | 0.39 | 0.08 | 2.13 | 0.11 | 0.08 | 0.7 | 0 | 0 | 1.61 | 0 | 0.78 | 1.54 |
| Ifna2 | NM_010503.2 | NM_010503.2:89 | 0.39 | 0.08 | 2.13 | 0.11 | 0.08 | 0.7 | 0 | 0 | 1.61 | 0 | 0.78 | 0.02 |
| Ifna4 | NM_010504.2 | NM_010504.2:261 | 0.39 | 0.08 | 2.13 | 0.11 | 0.08 | 0.7 | 0 | 0 | 1.61 | 0 | 0.78 | 0.02 |
| Ifnar1 | NM_010508.1 | NM_010508.1:1195 | 8.42 | 7.24 | 8.33 | 8 | 7.9 | 8.65 | 7.54 | 9.12 | 9.69 | 7.41 | 9.32 | 8.85 |
| Ifnar2 | NM_001110498.1 | NM_001110498.1:725 | 13.51 | 13.65 | 13.79 | 13.5 | 13.31 | 13.44 | 12.91 | 13.5 | 13.99 | 12.84 | 13.34 | 13.76 |
| Ifnb1 | NM_010510.1 | NM_010510.1:335 | 0.39 | 6.34 | 2.13 | 4.82 | 0.08 | 0.7 | 4.54 | 5.38 | 4.39 | 2.6 | 0.78 | 3.59 |
| Ifng | NM_008337.1 | NM_008337.1:95 | 0.39 | 0.08 | 2.13 | 0.11 | 0.08 | 0.7 | 0 | 0 | 1.61 | 0 | 0.78 | 0.02 |
| Ifngr1 | NM_010511.2 | NM_010511.2:985 | 10.49 | 10.77 | 10.96 | 9.84 | 10.46 | 10.3 | 9.34 | 10.09 | 10.5 | 9.69 | 10.35 | 11.24 |
| Ifnl2 | NM_001024673.2 | NM_001024673.2:72 | 0.39 | 0.08 | 2.13 | 0.11 | 0.08 | 0.7 | 0 | 0 | 1.61 | 0 | 0.78 | 0.02 |
| Igf1r | NM_010513.2 | NM_010513.2:3390 | 0.39 | 0.08 | 2.13 | 0.11 | 0.08 | 0.7 | 0 | 0 | 1.61 | 5.77 | 8.16 | 8.65 |
| Igf2r | NM_010515.1 | NM_010515.1:2585 | 5.14 | 4.31 | 2.46 | 3.13 | 4.26 | 4.3 | 0.32 | 4.5 | 4.91 | 5.97 | 2.53 | 5.01 |
| Igll1 | NM_001190325.1 | NM_001190325.1:690 | 0.39 | 0.08 | 2.13 | 0.11 | 0.08 | 0.7 | 0 | 0 | 1.61 | 0 | 0.78 | 0.02 |
| Ikbkb | NM_010546.2 | NM_010546.2:498 | 8.04 | 7.35 | 7.39 | 6.95 | 6.35 | 6.4 | 6.31 | 6.11 | 7.79 | 6.52 | 6.25 | 6.83 |
| Ikbke | NM_019777.3 | NM_019777.3:618 | 8.9 | 8.96 | 8.27 | 7.29 | 7.69 | 7.43 | 7.01 | 8.77 | 8.71 | 8.14 | 7.05 | 9.95 |
| Ikbkg | NM_178590.2 | NM_178590.2:525 | 3.71 | 4.52 | 6.4 | 5.36 | 4.26 | 5.56 | 0.73 | 3.79 | 6.12 | 4.01 | 4.52 | 1.97 |

| | | | | | | | | | | | | | | |
|---|---|---|---|---|---|---|---|---|---|---|---|---|---|---|
| Ikzf1 | NM_001025597.1 | NM_001025597.1:4420 | 10.38 | 10.83 | 11.04 | 10.57 | 10.58 | 10.46 | 9.21 | 9.63 | 10.61 | 9.47 | 11.07 | 10.55 |
| Ikzf2 | NM_011770.4 | NM_011770.4:7230 | 7.21 | 6.07 | 6.79 | 8.27 | 4 | 7.53 | 6.15 | 0 | 7.46 | 8.46 | 4.62 | 9.37 |
| Il10 | NM_010548.1 | NM_010548.1:985 | 11.58 | 10.85 | 10.45 | 9.75 | 10.63 | 10.33 | 10.65 | 11.06 | 11.3 | 9.93 | 10.2 | 11.45 |
| Il10ra | NM_008348.2 | NM_008348.2:75 | 7.13 | 8.9 | 7.15 | 7.38 | 6.71 | 6.66 | 7.56 | 7 | 7.44 | 6.53 | 7.09 | 7.88 |
| Il11 | NM_008350.2 | NM_008350.2:285 | 0.39 | 5.11 | 2.13 | 2.16 | 4 | 0.7 | 1.9 | 0.25 | 3.56 | 1.01 | 3.45 | 2.3 |
| Il11ra1 | NM_010549.3 | NM_010549.3:22 | 0.39 | 0.08 | 5.18 | 0.11 | 0.08 | 0.7 | 0 | 0 | 1.61 | 0 | 0.78 | 4.71 |
| Il12a | NM_008351.1 | NM_008351.1:355 | 0.39 | 0.08 | 2.13 | 0.11 | 0.08 | 0.7 | 0 | 0 | 1.61 | 0 | 0.78 | 0.02 |
| Il12b | NM_008352.1 | NM_008352.1:1045 | 0.39 | 6.24 | 4.22 | 4.76 | 6.78 | 5.29 | 0 | 0 | 1.61 | 2.24 | 5.5 | 2.99 |
| Il12rb1 | NM_008353.2 | NM_008353.2:1088 | 0.39 | 0.08 | 2.13 | 1.76 | 0.08 | 0.7 | 4.13 | 0 | 1.61 | 0 | 1.24 | 0.02 |
| Il12rb2 | NM_008354.3 | NM_008354.3:1395 | 0.39 | 3.03 | 2.13 | 0.11 | 0.08 | 0.7 | 0 | 0 | 1.61 | 2.89 | 0.78 | 6.91 |
| Il13 | NM_008355.2 | NM_008355.2:425 | 0.39 | 2.83 | 2.13 | 0.11 | 0.08 | 0.7 | 0 | 0 | 1.61 | 0 | 0.78 | 0.02 |
| Il13ra1 | NM_133990.4 | NM_133990.4:845 | 7.89 | 5.53 | 6.79 | 6.54 | 6.9 | 6.16 | 6.91 | 7.21 | 7.69 | 8.25 | 8.37 | 9.12 |
| Il13ra2 | NM_008356.3 | NM_008356.3:1010 | 0.39 | 0.08 | 2.13 | 0.11 | 0.08 | 0.7 | 0 | 0 | 1.61 | 0 | 0.78 | 0.02 |
| Il15 | NM_008357.2 | NM_008357.2:854 | 6.81 | 5.77 | 6.4 | 5.87 | 5.26 | 6.16 | 5.04 | 6.22 | 6.46 | 7.26 | 7.11 | 8.47 |
| Il15ra | NM_008358.2 | NM_008358.2:800 | 0.39 | 0.08 | 2.13 | 0.11 | 0.08 | 0.7 | 0 | 0 | 1.61 | 0 | 0.78 | 3.59 |
| Il16 | NM_010551.3 | NM_010551.3:3095 | 0.39 | 0.08 | 2.13 | 0.11 | 0.08 | 0.7 | 3.59 | 0 | 1.61 | 2.89 | 7.46 | 0.02 |
| Il17a | NM_010552.3 | NM_010552.3:205 | 0.39 | 0.08 | 2.13 | 3.59 | 0.08 | 0.7 | 0 | 0 | 1.61 | 5.56 | 5.26 | 0.02 |
| Il17b | NM_019508.1 | NM_019508.1:346 | 0.39 | 0.08 | 2.13 | 2.16 | 0.08 | 0.87 | 0 | 0.98 | 4.58 | 3.43 | 6.18 | 4.92 |
| Il17f | NM_145856.2 | NM_145856.2:625 | 4.39 | 5.24 | 4.53 | 4.76 | 4.61 | 4.41 | 3.69 | 4.73 | 5.92 | 6.13 | 4.98 | 4.82 |
| Il17ra | NM_008359.1 | NM_008359.1:312 | 9.57 | 9.63 | 8.57 | 6.42 | 8.18 | 7.44 | 7.71 | 8.44 | 9.27 | 9.57 | 9.88 | 10.5 |
| Il17rb | NM_019583.3 | NM_019583.3:1338 | 0.39 | 0.08 | 2.13 | 0.11 | 0.08 | 0.7 | 0 | 0 | 1.61 | 0 | 0.78 | 0.02 |
| Il18 | NM_008360.1 | NM_008360.1:270 | 6.56 | 7.29 | 7.73 | 6.87 | 6.21 | 6.63 | 5.99 | 6.67 | 6.64 | 6.76 | 8.3 | 8.83 |
| Il18r1 | NM_001161842.1 | NM_001161842.1:620 | 0.39 | 7.1 | 4.22 | 3.59 | 0.08 | 2.74 | 6.6 | 0 | 1.61 | 7.75 | 6.72 | 7.97 |
| Il18rap | NM_010553.2 | NM_010553.2:2055 | 5.48 | 4.52 | 4.53 | 4.76 | 2.91 | 0.7 | 3.86 | 3.19 | 6.06 | 9.17 | 9.45 | 10.9 |
| Il19 | NM_001009940.1 | NM_001009940.1:464 | 0.39 | 0.89 | 3.84 | 3.59 | 0.08 | 0.7 | 0 | 0 | 5.4 | 0 | 0.78 | 8 |
| Il1a | NM_010554.4 | NM_010554.4:225 | 9.98 | 10.48 | 10.02 | 9.52 | 9.48 | 9.38 | 8.71 | 9.39 | 10.42 | 10.54 | 10.72 | 12.44 |
| Il1b | NM_008361.3 | NM_008361.3:1120 | 15.78 | 16.4 | 15.96 | 15.93 | 14.91 | 15.75 | 14.81 | 15.53 | 15.94 | 17.15 | 17.65 | 18.53 |
| Il1r1 | NM_001123382.1 | NM_001123382.1:820 | 3.71 | 0.08 | 5.34 | 0.11 | 0.08 | 4.96 | 2.44 | 5.46 | 1.61 | 6.17 | 8.29 | 7.12 |
| Il1r2 | NM_010555.4 | NM_010555.4:458 | 9.81 | 10.4 | 9.57 | 8.27 | 9.18 | 9.24 | 8.38 | 6.88 | 8.53 | 13.66 | 13.95 | 14.94 |
| Il1rap | NM_008364.2 | NM_008364.2:2415 | 4.78 | 5.6 | 2.13 | 4.29 | 2.44 | 2.34 | 4.84 | 2.57 | 1.61 | 8.55 | 8.44 | 9.18 |
| Il1rapl2 | NM_030688.1 | NM_030688.1:202 | 5.03 | 5.43 | 7.01 | 5.64 | 5.7 | 5.87 | 3.98 | 5.32 | 6.99 | 5.43 | 5.92 | 5.84 |
| Il1rl1 | NM_001025602.2 | NM_001025602.2:815 | 0.39 | 0.08 | 2.13 | 3.93 | 4.95 | 3.89 | 2.19 | 3.52 | 4.58 | 5.62 | 0.78 | 5.53 |
| Il1rl2 | NM_133193.3 | NM_133193.3:860 | 0.39 | 4.06 | 2.13 | 0.11 | 0.08 | 0.7 | 3.69 | 0 | 1.61 | 3.68 | 0.78 | 0.02 |
| Il1rn | NM_031167.5 | NM_031167.5:224 | 11.42 | 11.89 | 12.14 | 9.78 | 10.64 | 10.56 | 10.37 | 11.23 | 12.17 | 12.32 | 12.28 | 13.62 |
| Il2 | NM_008366.3 | NM_008366.3:314 | 0.39 | 0.08 | 2.13 | 0.11 | 0.08 | 0.7 | 0 | 0 | 1.61 | 3.01 | 0.78 | 0.02 |
| Il21 | NM_021782.2 | NM_021782.2:1762 | 0.39 | 0.08 | 2.13 | 0.11 | 0.08 | 0.7 | 0 | 0 | 1.61 | 0 | 0.78 | 0.02 |
| Il21r | NM_021887.1 | NM_021887.1:619 | 7.13 | 7.4 | 6.24 | 7.41 | 7.37 | 6.81 | 7.56 | 7.52 | 7.87 | 4.24 | 0.78 | 4.09 |
| Il22 | NM_016971.1 | NM_016971.1:477 | 0.39 | 0.08 | 2.13 | 0.11 | 0.08 | 0.7 | 0 | 0 | 1.61 | 0 | 0.78 | 0.02 |
| Il22ra1 | NM_178257.1 | NM_178257.1:685 | 0.39 | 0.08 | 2.13 | 0.28 | 0.08 | 0.7 | 0 | 0.98 | 1.61 | 3.75 | 0.78 | 0.02 |
| Il22ra2 | NM_178258.5 | NM_178258.5:20 | 0.39 | 0.08 | 2.13 | 0.28 | 0.08 | 0.7 | 0.73 | 0 | 1.61 | 5.34 | 0.78 | 3.46 |
| Il23a | NM_031252.1 | NM_031252.1:360 | 5.64 | 7.97 | 5.34 | 2.94 | 5.92 | 6.26 | 1.73 | 4.64 | 5.5 | 10.19 | 10.04 | 12.61 |
| Il23r | NM_144548.1 | NM_144548.1:690 | 0.39 | 4.97 | 2.13 | 1.76 | 0.08 | 0.7 | 0 | 0 | 1.61 | 3.89 | 0.78 | 0.02 |
| Il24 | NM_053095.2 | NM_053095.2:545 | 0.39 | 0.08 | 2.13 | 0.11 | 0.08 | 0.7 | 0 | 0 | 1.61 | 0 | 0.78 | 0.02 |
| Il25 | NM_080729.2 | NM_080729.2:649 | 0.39 | 0.08 | 2.13 | 0.11 | 0.08 | 0.7 | 0 | 0 | 1.61 | 0 | 0.78 | 5.4 |
| Il27 | NM_145636.1 | NM_145636.1:402 | 0.39 | 0.08 | 2.13 | 0.11 | 0.08 | 0.7 | 0 | 0 | 1.61 | 0 | 0.78 | 0.02 |
| Il2ra | NM_008367.2 | NM_008367.2:325 | 6.54 | 8.24 | 6.07 | 6.84 | 4.99 | 5.41 | 0 | 4.02 | 1.61 | 7.11 | 0.78 | 0.02 |
| Il2rb | NM_008368.3 | NM_008368.3:2365 | 0.39 | 3.86 | 3.84 | 5.2 | 5.13 | 3.53 | 0.73 | 6.6 | 2.51 | 6.95 | 0.78 | 0.02 |
| Il2rg | NM_013563.3 | NM_013563.3:1226 | 11.88 | 11.84 | 11.46 | 11.24 | 11.16 | 11.57 | 10.79 | 11.29 | 11.93 | 10.32 | 11.44 | 11.99 |
| Il3 | NM_010556.4 | NM_010556.4:155 | 0.39 | 2.33 | 2.13 | 2.16 | 0.08 | 0.7 | 0 | 0 | 1.61 | 2.75 | 0.78 | 2.57 |
| Il34 | NM_001135100.1 | NM_001135100.1:848 | 0.39 | 0.08 | 2.13 | 0.11 | 0.08 | 0.7 | 0 | 0 | 1.61 | 2.01 | 0.78 | 1.54 |
| Il3ra | NM_008369.1 | NM_008369.1:567 | 0.39 | 0.08 | 2.13 | 0.11 | 0.08 | 0.7 | 0 | 0 | 1.61 | 2.6 | 0.78 | 0.02 |
| Il4 | NM_021283.1 | NM_021283.1:345 | 0.39 | 0.08 | 2.13 | 3.45 | 0.25 | 0.7 | 0.73 | 1.47 | 3.56 | 0 | 0.78 | 0.02 |
| Il4ra | NM_010557.1 | NM_010557.1:3017 | 10.79 | 11.15 | 10.28 | 9.7 | 9.98 | 9.87 | 10 | 10.44 | 10.66 | 10.32 | 10.6 | 11.18 |
| Il5 | NM_010558.1 | NM_010558.1:177 | 2.97 | 0.08 | 2.13 | 0.11 | 2.44 | 2.74 | 1.54 | 2.57 | 4.58 | 2.6 | 0.78 | 0.02 |
| Il5ra | NM_008370.2 | NM_008370.2:2290 | 0.39 | 0.08 | 2.13 | 0.11 | 0.08 | 0.7 | 0 | 0 | 1.61 | 0 | 0.78 | 4.92 |
| Il6 | NM_031168.1 | NM_031168.1:40 | 4.39 | 4.45 | 6.24 | 5.47 | 5.17 | 5.1 | 4.32 | 3.62 | 6.96 | 5.18 | 0.78 | 0.92 |
| Il6ra | NM_010559.2 | NM_010559.2:2825 | 10.29 | 10.47 | 9.91 | 9.57 | 9.27 | 9.61 | 9.25 | 9.56 | 10.39 | 9.97 | 11.04 | 11.57 |
| Il6st | NM_010560.2 | NM_010560.2:2325 | 0.39 | 0.08 | 2.13 | 0.11 | 0.08 | 0.7 | 2.44 | 0 | 1.61 | 0 | 0.78 | 0.02 |
| Il7 | NM_008371.2 | NM_008371.2:1055 | 5.09 | 3.51 | 2.13 | 3.13 | 0.08 | 0.7 | 2.73 | 3.05 | 5.29 | 4.18 | 2.03 | 2.3 |
| Il7r | NM_008372.3 | NM_008372.3:1020 | 10.64 | 11.51 | 11.33 | 9.55 | 10.42 | 11.19 | 9.01 | 10.98 | 12.21 | 8.95 | 11.04 | 9.42 |
| Il9 | NM_008373.1 | NM_008373.1:39 | 0.39 | 0.08 | 2.13 | 1.2 | 0.08 | 0.7 | 0 | 0 | 1.61 | 0 | 0.78 | 0.02 |
| Ilf3 | NM_010561.2 | NM_010561.2:1902 | 10.42 | 10.67 | 10.04 | 10.19 | 9.66 | 9.33 | 10.34 | 9.87 | 10.67 | 9.61 | 10.48 | 10.62 |
| Inpp5d | NM_001110192.1 | NM_001110192.1:2186 | 7.92 | 7.21 | 7.76 | 7.03 | 7.28 | 6.89 | 7.14 | 8.65 | 8.74 | 7.37 | 7.9 | 8.19 |
| Irak1 | NM_008363.2 | NM_008363.2:951 | 6.56 | 7.79 | 8.31 | 7.7 | 7.98 | 7.61 | 7.5 | 8.47 | 9.33 | 7.29 | 9.44 | 7.68 |
| Irak2 | NM_001113553.1 | NM_001113553.1:485 | 7.99 | 8.01 | 7.5 | 6.36 | 7.03 | 7.51 | 6.36 | 7.98 | 7.53 | 8.29 | 8.45 | 9.51 |
| Irak3 | NM_028679.3 | NM_028679.3:2608 | 8.09 | 7.53 | 8.27 | 7.54 | 6.21 | 7.39 | 7.18 | 7.86 | 8 | 8.2 | 7.79 | 9.33 |
| Irak4 | NM_029926.5 | NM_029926.5:250 | 7.63 | 7.74 | 8.05 | 6.36 | 6.53 | 7.23 | 7.07 | 6.63 | 7.51 | 7.82 | 9.41 | 8.14 |
| Irf1 | NM_008390.1 | NM_008390.1:365 | 6.2 | 8.23 | 5.34 | 6.34 | 6.98 | 6.4 | 5.71 | 6.86 | 5.5 | 6.61 | 7.24 | 8.47 |
| Irf2 | NM_008391.2 | NM_008391.2:440 | 10.28 | 10.16 | 10.47 | 10.03 | 9.71 | 9.6 | 9.22 | 9.46 | 10.24 | 8.88 | 9.43 | 9.73 |
| Irf3 | NM_016849.4 | NM_016849.4:526 | 5.78 | 7.36 | 4.99 | 5.84 | 4.73 | 5.23 | 5.76 | 6.38 | 6.35 | 5.36 | 0.78 | 6.2 |
| Irf4 | NM_013674.1 | NM_013674.1:1878 | 6 | 4.15 | 5.49 | 5.54 | 3.27 | 5.41 | 4.16 | 1.47 | 1.61 | 3.01 | 0.78 | 1.54 |

| | | | | | | | | | | | | | | |
|---|---|---|---|---|---|---|---|---|---|---|---|---|---|---|
| Irf5 | NM_012057.3 | NM_012057.3:1826 | 12.95 | 12.94 | 12.57 | 12.74 | 12.51 | 12.46 | 12.37 | 12.74 | 12.91 | 11.66 | 11.79 | 12.6 |
| Irf7 | NM_016850.2 | NM_016850.2:705 | 9.63 | 9.24 | 9.24 | 6.51 | 8.26 | 7.53 | 8.98 | 9.33 | 9.61 | 8.39 | 5.32 | 7.18 |
| Irf8 | NM_008320.3 | NM_008320.3:2274 | 9.62 | 9.2 | 9.94 | 10.02 | 10.17 | 9.53 | 8.68 | 9.35 | 10.04 | 7.97 | 7.68 | 7.87 |
| Irgm2 | NM_019440.2 | NM_019440.2:2555 | 0.39 | 0.08 | 2.13 | 0.11 | 0.08 | 3.53 | 0 | 0 | 1.61 | 5.18 | 2.03 | 5.37 |
| Isg15 | NM_015783.1 | NM_015783.1:395 | 10.24 | 10.16 | 10.31 | 9.91 | 9.37 | 9.59 | 8.84 | 9.07 | 10.37 | 10.26 | 10.45 | 10.74 |
| Isg20 | NM_020583.5 | NM_020583.5:552 | 9.46 | 8.71 | 9.34 | 7.51 | 8.45 | 8.68 | 9.02 | 9.27 | 9.79 | 11.06 | 10.67 | 11.09 |
| Itch | NM_008395.2 | NM_008395.2:575 | 9.7 | 9.34 | 9.7 | 9.12 | 10.1 | 9.04 | 8.6 | 9.79 | 10.13 | 9.14 | 10.05 | 10.53 |
| Itga1 | NM_001033228.3 | NM_001033228.3:2550 | 0.39 | 0.08 | 2.13 | 0.11 | 0.08 | 0.7 | 0 | 0 | 1.61 | 0 | 5.2 | 1.54 |
| Itga2 | NM_008396.2 | NM_008396.2:1860 | 0.39 | 0.08 | 2.13 | 0.11 | 0.08 | 0.7 | 0 | 0 | 1.61 | 0 | 0.78 | 0.02 |
| Itga2b | NM_010575.2 | NM_010575.2:461 | 0.39 | 0.08 | 2.13 | 0.11 | 0.08 | 0.7 | 0 | 0 | 1.61 | 0 | 0.78 | 0.02 |
| Itga4 | NM_010576.3 | NM_010576.3:6600 | 6.25 | 8.79 | 7.01 | 7.59 | 6.17 | 6.13 | 5.41 | 5.98 | 6.77 | 5.5 | 7.07 | 4.53 |
| Itga5 | NM_010577.3 | NM_010577.3:2446 | 8.99 | 10.61 | 9.39 | 9.29 | 9.39 | 9.43 | 8.6 | 10.21 | 10.82 | 8.52 | 9.62 | 10.15 |
| Itga6 | NM_008397.3 | NM_008397.3:910 | 9.13 | 9.27 | 9.13 | 8.98 | 8.78 | 8.53 | 9.05 | 9.29 | 9.95 | 9.21 | 8.24 | 8.34 |
| Itgae | NM_008399.1 | NM_008399.1:775 | 4.3 | 6.41 | 6.47 | 7.17 | 7.57 | 6.85 | 5.5 | 0 | 1.61 | 3.6 | 0.78 | 6.2 |
| Itgal | NM_008400.2 | NM_008400.2:950 | 7.77 | 8.09 | 8.02 | 6.44 | 7.17 | 6.43 | 7.21 | 7.86 | 7.69 | 9.75 | 9.65 | 9.64 |
| Itgam | NM_008401.2 | NM_008401.2:155 | 10.46 | 10.57 | 9.42 | 9.23 | 10.08 | 9.56 | 10.14 | 11.02 | 10.94 | 10.41 | 10.77 | 11.12 |
| Itgax | NM_021334.2 | NM_021334.2:327 | 4.64 | 6.84 | 2.13 | 4.51 | 5.79 | 4.96 | 5.3 | 4.97 | 2.51 | 6.56 | 5.13 | 6.14 |
| Itgb1 | NM_010578.1 | NM_010578.1:1855 | 11.16 | 10.94 | 10.16 | 10.29 | 10.76 | 10.93 | 11.19 | 10.98 | 10.84 | 11.37 | 10.89 | 11.25 |
| Itgb2 | NM_008404.4 | NM_008404.4:2542 | 12.14 | 12.46 | 12.32 | 12.27 | 12.06 | 12.21 | 11.26 | 11.95 | 12.4 | 11.69 | 12.18 | 12.59 |
| Itgb3 | NM_016780.2 | NM_016780.2:2595 | 0.39 | 2.83 | 5.18 | 2.94 | 5.79 | 3.89 | 6.98 | 0 | 3.56 | 6.81 | 6.15 | 7.46 |
| Itgb4 | NM_001005608.2 | NM_001005608.2:3355 | 0.39 | 2.6 | 2.13 | 3.13 | 0.08 | 0.7 | 1.06 | 2.12 | 1.61 | 0 | 0.78 | 1.54 |
| Itk | NM_010583.3 | NM_010583.3:404 | 0.39 | 0.08 | 3.3 | 0.11 | 0.08 | 0.7 | 1.73 | 1.47 | 1.61 | 2.6 | 1.24 | 3.7 |
| Jak1 | NM_146145.2 | NM_146145.2:4080 | 10.66 | 11.04 | 10.78 | 10.7 | 9.96 | 10.63 | 9.9 | 10 | 10.64 | 10.63 | 11.53 | 11.98 |
| Jak2 | NM_008413.2 | NM_008413.2:1049 | 10.14 | 10.96 | 10.23 | 9.18 | 9.79 | 10.42 | 9.66 | 10.26 | 10.54 | 10.67 | 11.12 | 12.5 |
| Jak3 | NM_010589.5 | NM_010589.5:145 | 0.39 | 0.08 | 2.13 | 0.11 | 0.08 | 0.7 | 1.54 | 8.15 | 1.61 | 4.01 | 0.78 | 5.26 |
| Jam3 | NM_023277.4 | NM_023277.4:145 | 0.39 | 0.08 | 2.13 | 0.11 | 0.08 | 0.7 | 0 | 0 | 1.61 | 0 | 0.78 | 0.02 |
| Jun | NM_010591.2 | NM_010591.2:2212 | 10.85 | 10.76 | 10.75 | 10.88 | 10.58 | 10.08 | 10.59 | 10.92 | 11.27 | 10.79 | 10.71 | 11.37 |
| Kdr | NM_010612.2 | NM_010612.2:995 | 4.97 | 7.21 | 5.86 | 4.87 | 4.18 | 5.66 | 5.62 | 5.99 | 6.6 | 0 | 4.15 | 4.09 |
| Kit | NM_001122733.1 | NM_001122733.1:4275 | 0.39 | 0.08 | 2.13 | 4.44 | 5.61 | 0.7 | 0 | 0 | 1.61 | 0 | 0.78 | 6.44 |
| Klra1 | NM_016659.3 | NM_016659.3:105 | 0.39 | 0.08 | 2.13 | 0.11 | 0.08 | 0.7 | 0 | 0 | 1.61 | 0 | 0.78 | 0.02 |
| Klra15 | NM_013793.2 | NM_013793.2:736 | 0.39 | 0.08 | 2.13 | 0.11 | 0.08 | 0.7 | 0 | 0 | 1.61 | 0 | 0.78 | 0.02 |
| Klra17 | NM_133203.4 | NM_133203.4:58 | 5.71 | 3.75 | 3.84 | 5.79 | 4.99 | 3.89 | 2.64 | 0 | 6.12 | 5.73 | 7.65 | 7.44 |
| Klra2 | NM_001170851.1 | NM_001170851.1:420 | 9.74 | 10.06 | 9.86 | 9.2 | 9.06 | 9.36 | 9 | 9.62 | 10.24 | 9.54 | 10.15 | 10.67 |
| Klra20 | NM_053150.2 | NM_053150.2:176 | 0.39 | 0.08 | 2.13 | 0.11 | 0.08 | 0.7 | 0 | 0 | 1.61 | 0 | 0.78 | 0.02 |
| Klra21 | NM_053151.1 | NM_053151.1:41 | 0.39 | 0.08 | 2.13 | 1.2 | 0.08 | 0.7 | 0 | 0 | 1.61 | 0 | 0.78 | 0.02 |
| Klra27 | XM_916590.3 | XM_916590.3:496 | 0.39 | 0.08 | 2.13 | 0.11 | 0.08 | 0.7 | 0 | 0 | 1.61 | 0 | 0.78 | 0.02 |
| Klra3 | NM_010648.2 | NM_010648.2:7 | 0.39 | 0.08 | 2.13 | 0.11 | 0.08 | 0.7 | 0 | 0 | 1.61 | 0 | 0.78 | 0.02 |
| Klra4 | NM_010649.3 | NM_010649.3:169 | 0.39 | 3.86 | 2.13 | 0.11 | 0.08 | 0.7 | 0 | 0 | 4.76 | 2.89 | 0.78 | 4.4 |
| Klra5 | NM_008463.2 | NM_008463.2:174 | 0.39 | 0.08 | 2.13 | 2.73 | 0.08 | 0.7 | 0 | 0 | 1.61 | 0 | 0.78 | 0.02 |
| Klra6 | NM_008464.2 | NM_008464.2:880 | 2.39 | 1.99 | 2.13 | 0.11 | 0.08 | 0.7 | 2.54 | 0 | 1.61 | 2.43 | 0.78 | 0.02 |
| Klra7 | NM_001110323.1 | NM_001110323.1:250 | 0.39 | 4.97 | 2.13 | 4.58 | 0.08 | 0.7 | 0 | 0 | 1.61 | 5.38 | 3.84 | 6.18 |
| Klrb1 | NM_001099918.1 | NM_001099918.1:327 | 0.39 | 0.08 | 2.13 | 0.11 | 0.08 | 0.7 | 0 | 0 | 1.61 | 0 | 0.78 | 0.02 |
| Klrb1c | NM_008527.2 | NM_008527.2:47 | 0.39 | 0.08 | 2.13 | 0.11 | 0.08 | 0.7 | 0 | 0 | 1.61 | 2.24 | 0.78 | 0.92 |
| Klrc1 | NM_001136068.1 | NM_001136068.1:68 | 0.39 | 0.08 | 5.49 | 0.11 | 0.08 | 0.7 | 0 | 0 | 3.13 | 4.38 | 0.78 | 3.81 |
| Klrc2 | NM_010653.4 | NM_010653.4:259 | 0.39 | 0.08 | 2.13 | 0.11 | 0.08 | 0.7 | 0 | 0 | 1.61 | 0 | 0.78 | 0.02 |
| Klrd1 | NM_010654.2 | NM_010654.2:434 | 4.97 | 6.61 | 7.84 | 7.31 | 6.53 | 8.28 | 1.9 | 0.25 | 5.18 | 3.75 | 5.05 | 0.02 |
| Klrg1 | NM_016970.1 | NM_016970.1:825 | 4.64 | 0.08 | 5.75 | 4.03 | 5.33 | 3.72 | 1.54 | 3.31 | 5.77 | 1.75 | 5.13 | 3.46 |
| Klrk1 | NM_001083322.1 | NM_001083322.1:144 | 8.17 | 7.22 | 9.13 | 5.97 | 8.84 | 8.13 | 7.55 | 0 | 6.99 | 3.82 | 1.24 | 5.22 |
| Lag3 | NM_008479.1 | NM_008479.1:1700 | 0.39 | 0.08 | 2.13 | 0.11 | 0.08 | 0.7 | 0 | 0 | 1.61 | 2.43 | 0.78 | 0.02 |
| Lamp1 | NM_010684.2 | NM_010684.2:2080 | 14.59 | 14.67 | 14.05 | 14.97 | 14.09 | 14.56 | 14.03 | 14.08 | 14.07 | 14.01 | 13.8 | 14.33 |
| Lamp2 | NM_001017959.1 | NM_001017959.1:908 | 10.07 | 9.7 | 9.62 | 9.59 | 9.54 | 9.87 | 9.54 | 9.62 | 9.74 | 10.26 | 10.76 | 11.32 |
| Lamp3 | NM_177356.3 | NM_177356.3:1074 | 3.56 | 0.08 | 5.97 | 7.21 | 4.09 | 3.06 | 2.73 | 3.71 | 3.89 | 6.86 | 2.91 | 3.46 |
| Lbp | NM_008489.2 | NM_008489.2:1015 | 7.21 | 5.56 | 2.13 | 5.82 | 3.1 | 5.23 | 7.76 | 6.65 | 7.03 | 8.59 | 7.55 | 8 |
| Lck | NM_010693.2 | NM_010693.2:1180 | 0.39 | 0.08 | 2.13 | 0.11 | 2.44 | 0.7 | 0.73 | 0 | 1.61 | 0 | 0.78 | 0.02 |
| Lcn2 | NM_008491.1 | NM_008491.1:190 | 0.39 | 0.08 | 2.13 | 0.11 | 0.08 | 0.7 | 5.53 | 0 | 5.5 | 9.71 | 9.63 | 10.38 |
| Lcp1 | NM_001247984.1 | NM_001247984.1:3344 | 13.46 | 13.58 | 12.97 | 13.43 | 12.91 | 13.09 | 13.13 | 13.19 | 13.33 | 13.79 | 13.83 | 14.89 |
| Lgals3 | NM_001145953.1 | NM_001145953.1:665 | 12.83 | 13.34 | 12.84 | 12.9 | 12.7 | 12.79 | 12.21 | 12.89 | 13.06 | 12.29 | 12.49 | 13.11 |
| Lif | NM_008501.2 | NM_008501.2:3435 | 5.39 | 7.55 | 7.76 | 6.71 | 5.67 | 5.17 | 7.11 | 5.6 | 8.49 | 4.72 | 6.72 | 5.37 |
| Lilra5 | NM_001081239.2 | NM_001081239.2:994 | 10.52 | 10.05 | 7.63 | 10.53 | 8.71 | 8.41 | 8.77 | 7.03 | 8.51 | 7.34 | 6.69 | 8.47 |
| Litaf | NM_019980.1 | NM_019980.1:1100 | 11.85 | 11.46 | 11.39 | 11.05 | 11.1 | 11.38 | 11.08 | 11.3 | 11.24 | 12.43 | 12.63 | 13.48 |
| Lrp1 | NM_008512.2 | NM_008512.2:1310 | 9.66 | 9.75 | 9.15 | 8.06 | 9.36 | 9.51 | 9.28 | 9.68 | 10.39 | 7.85 | 10.26 | 8.35 |
| Lrrn3 | NM_010733.2 | NM_010733.2:1765 | 2.97 | 3.63 | 2.13 | 1.2 | 3.1 | 3.53 | 0 | 0 | 2.51 | 1.43 | 0.78 | 2.8 |
| Lta | NM_010735.1 | NM_010735.1:1115 | 5.2 | 3.86 | 6.9 | 6.38 | 6.48 | 5.87 | 3.69 | 5.72 | 8.32 | 4.56 | 7.01 | 6.63 |
| Ltb | NM_008518.2 | NM_008518.2:163 | 7.2 | 6.8 | 2.13 | 0.11 | 6.08 | 5.61 | 0 | 5.86 | 6.3 | 7.09 | 9.33 | 9.37 |
| Ltbr | NM_010736.3 | NM_010736.3:1962 | 14.59 | 14.71 | 14.44 | 14.86 | 14.19 | 14.14 | 14.2 | 14.38 | 14.62 | 14.75 | 14.29 | 15.39 |
| Ltf | NM_008522.3 | NM_008522.3:2545 | 0.39 | 5.11 | 2.13 | 0.11 | 4.34 | 4.8 | 3.32 | 0 | 1.61 | 3.68 | 7.34 | 8.91 |
| Ltk | NM_008523.2 | NM_008523.2:788 | 0.39 | 0.08 | 2.13 | 1.76 | 0.08 | 0.7 | 0 | 0 | 1.61 | 1.75 | 0.78 | 3.7 |
| Ly86 | NM_010745.2 | NM_010745.2:725 | 10.68 | 11.45 | 10.85 | 11.28 | 10.88 | 11 | 10.25 | 9.89 | 10.47 | 9.95 | 8.34 | 9.68 |
| Ly9 | NM_008534.2 | NM_008534.2:1190 | 8.72 | 8.3 | 8.41 | 8.31 | 8.67 | 8.68 | 8.44 | 9.17 | 8.58 | 9.01 | 8.97 | 9.89 |
| Ly96 | NM_001159711.1 | NM_001159711.1:64 | 9.26 | 9.51 | 8.6 | 9.74 | 9.02 | 9.16 | 8.96 | 9.41 | 8.68 | 8.49 | 8.94 | 9.02 |
| Lyn | NM_010747.1 | NM_010747.1:1725 | 12.52 | 12.68 | 12.13 | 12.34 | 12.2 | 11.87 | 12.15 | 12.48 | 12.51 | 12.86 | 12.95 | 13.84 |

| | | | | | | | | | | | | | | |
|---|---|---|---|---|---|---|---|---|---|---|---|---|---|---|
| Lyve1 | NM_053247.4 | NM_053247.4:150 | 7.51 | 7 | 6.16 | 6.32 | 6.86 | 6.99 | 6.93 | 6.68 | 5.68 | 5.24 | 6.59 | 4.33 |
| Lyz2 | NM_017372.3 | NM_017372.3:98 | 12.08 | 12.1 | 11.15 | 10.71 | 11.96 | 11.5 | 11.15 | 11.57 | 11.64 | 10.99 | 10.47 | 10.32 |
| Maf | NM_001025577.2 | NM_001025577.2:43 | 3.39 | 5.56 | 2.13 | 0.11 | 4.95 | 0.7 | 4.86 | 2.57 | 5.68 | 2.24 | 0.78 | 0.02 |
| Map2k1 | NM_008927.3 | NM_008927.3:1695 | 10.22 | 10.59 | 10.16 | 10.18 | 9.99 | 10.51 | 9.7 | 10.04 | 10.72 | 9.91 | 10.1 | 10.88 |
| Map2k2 | NM_023138.4 | NM_023138.4:1440 | 9.62 | 9.45 | 10.1 | 10.4 | 8.71 | 9.51 | 9.54 | 9.09 | 9.56 | 9.87 | 9.41 | 9.92 |
| Map2k4 | NM_009157.4 | NM_009157.4:1335 | 8.84 | 8.91 | 8.53 | 8.93 | 8.96 | 8.46 | 9.01 | 8.31 | 9.25 | 9.91 | 9.87 | 10.58 |
| Map3k1 | NM_011945.2 | NM_011945.2:1640 | 4.91 | 4.59 | 2.13 | 0.11 | 0.08 | 2.34 | 3.13 | 3.62 | 1.61 | 0.43 | 0.78 | 2.8 |
| Map3k5 | NM_008580.4 | NM_008580.4:640 | 6.56 | 7.43 | 2.13 | 0.28 | 5.3 | 0.7 | 5.42 | 7.13 | 1.61 | 8 | 8 | 6.92 |
| Map3k7 | NM_009316.1 | NM_009316.1:822 | 9.68 | 10.2 | 10.1 | 9.88 | 9.94 | 10 | 9.3 | 9.71 | 10.28 | 9.73 | 8.88 | 9.67 |
| Map4k2 | NM_009006.2 | NM_009006.2:666 | 6.34 | 4.82 | 2.13 | 0.11 | 5.26 | 0.7 | 5.63 | 0 | 1.61 | 7.15 | 9.7 | 8.93 |
| Mapk1 | NM_011949.3 | NM_011949.3:2640 | 11.97 | 11.99 | 12.02 | 12.36 | 11.79 | 12.02 | 11.48 | 12.13 | 11.98 | 11.94 | 11.9 | 12.28 |
| Mapk11 | NM_011161.5 | NM_011161.5:396 | 0.39 | 3.75 | 2.13 | 3.93 | 0.08 | 0.7 | 2.98 | 3.42 | 3.89 | 4.6 | 0.78 | 4.25 |
| Mapk14 | NM_001168513.1 | NM_001168513.1:114 | 8.57 | 7.79 | 8.05 | 8.06 | 8.77 | 8.14 | 7.83 | 7.38 | 8.79 | 8.5 | 8.51 | 8.87 |
| Mapk3 | NM_011952.2 | NM_011952.2:825 | 8.17 | 8 | 7.01 | 8.05 | 7.95 | 8.28 | 6.67 | 8.34 | 8.35 | 7.93 | 7.23 | 7.3 |
| Mapk8 | NM_016700.3 | NM_016700.3:970 | 9.24 | 9.26 | 8.74 | 8.49 | 8.28 | 8.24 | 9.09 | 8.82 | 9.2 | 8.99 | 8.42 | 9.51 |
| Mapkapk2 | NM_008551.1 | NM_008551.1:1991 | 8.92 | 9.12 | 8.02 | 6.24 | 8.15 | 7.95 | 8.31 | 8.39 | 8.64 | 10.16 | 10.22 | 10.36 |
| Marco | NM_010766.2 | NM_010766.2:350 | 6.8 | 6.93 | 6.16 | 6.52 | 5.97 | 6.56 | 5.7 | 5.94 | 6.3 | 5.21 | 1.24 | 5.26 |
| Masp1 | NM_008555.2 | NM_008555.2:210 | 0.39 | 0.08 | 2.13 | 0.11 | 0.08 | 0.7 | 0 | 0 | 1.61 | 0 | 0.78 | 0.02 |
| Masp2 | NM_010767.3 | NM_010767.3:363 | 0.39 | 0.08 | 2.13 | 0.11 | 0.08 | 0.7 | 0 | 0 | 3.13 | 0 | 0.78 | 0.02 |
| Mavs | NM_144888.1 | NM_144888.1:2510 | 0.39 | 5.91 | 6.32 | 5.9 | 5.79 | 4.62 | 3.06 | 5.4 | 2.51 | 4.95 | 0.78 | 0.02 |
| Mbl2 | NM_010776.1 | NM_010776.1:525 | 2.97 | 0.89 | 2.13 | 0.28 | 2.13 | 2.74 | 1.9 | 3.19 | 1.61 | 1.01 | 0.78 | 0.02 |
| Mcam | NM_023061.2 | NM_023061.2:630 | 0.39 | 0.08 | 2.13 | 0.11 | 0.08 | 0.7 | 5.19 | 0 | 6.24 | 5.72 | 0.78 | 6.39 |
| Mef2c | NM_001170537.1 | NM_001170537.1:4341 | 6.09 | 6.92 | 5.86 | 6.09 | 6.58 | 7.44 | 6.64 | 7.55 | 6.46 | 5.8 | 1.24 | 4.71 |
| Mefv | NM_001161790.1 | NM_001161790.1:928 | 7.8 | 9.01 | 8.02 | 4.12 | 6.06 | 6.38 | 5.06 | 6.85 | 8.62 | 9.11 | 8.64 | 10.51 |
| Mertk | NM_008587.1 | NM_008587.1:1320 | 8.76 | 8.29 | 5.49 | 6.81 | 8.33 | 7.57 | 5.36 | 7.77 | 7.96 | 0 | 5.7 | 6.2 |
| Mfge8 | NM_008594.2 | NM_008594.2:1357 | 6.64 | 5.5 | 2.13 | 1.76 | 5.09 | 5.17 | 5.14 | 2.75 | 6.24 | 7.47 | 6.95 | 5.3 |
| Mif | NM_010798.2 | NM_010798.2:373 | 12.42 | 12.68 | 12.49 | 12.34 | 12.01 | 12.52 | 11.88 | 12.01 | 12.96 | 12.62 | 12.95 | 13.13 |
| Mill2 | NM_153760.2 | NM_153760.2:22 | 4.39 | 0.08 | 2.13 | 3.71 | 0.08 | 4.41 | 0 | 2.12 | 3.56 | 5.77 | 0.78 | 0.02 |
| Mme | NM_008604.3 | NM_008604.3:285 | 6.71 | 5.24 | 2.13 | 5.92 | 3.8 | 0.7 | 7.63 | 5.48 | 5.59 | 9 | 7.05 | 0.02 |
| Mmp9 | NM_013599.2 | NM_013599.2:1570 | 5.64 | 5.69 | 2.13 | 0.11 | 3.68 | 4.3 | 5.56 | 2.91 | 5.18 | 8.21 | 8.17 | 10.12 |
| Mnx1 | NM_019944.1 | NM_019944.1:1740 | 0.39 | 0.08 | 2.13 | 0.11 | 0.08 | 0.7 | 0 | 0 | 1.61 | 0 | 0.78 | 0.02 |
| Mpo | NM_010824.2 | NM_010824.2:1648 | 4.78 | 4.87 | 6.85 | 5.54 | 7.03 | 6.48 | 4.96 | 5.7 | 5.77 | 5.36 | 4.01 | 8.04 |
| Mpped1 | NM_172610.3 | NM_172610.3:424 | 0.39 | 0.08 | 2.13 | 0.11 | 0.08 | 0.7 | 0 | 0 | 1.61 | 6.7 | 0.78 | 0.02 |
| Mr1 | NM_008209.4 | NM_008209.4:1360 | 7.89 | 8.37 | 7.05 | 7.85 | 8.04 | 8.04 | 4.9 | 5.7 | 4.16 | 5.34 | 6.59 | 6.06 |
| Mrc1 | NM_008625.1 | NM_008625.1:3992 | 11.52 | 11.21 | 10.39 | 10.61 | 10.64 | 10.31 | 10.88 | 10.54 | 11.02 | 9.82 | 9.42 | 10.33 |
| Ms4a1 | NM_007641.5 | NM_007641.5:166 | 0.39 | 0.08 | 2.13 | 0.11 | 0.08 | 0.7 | 0 | 0 | 1.61 | 0 | 0.78 | 0.02 |
| Ms4a2 | NM_001276330.1 | NM_001276330.1:104 | 3.39 | 0.08 | 2.13 | 0.11 | 1.73 | 0.7 | 0 | 0 | 4.91 | 0 | 0.78 | 0.02 |
| Msln | NM_018857.1 | NM_018857.1:346 | 0.39 | 0.08 | 2.13 | 0.11 | 0.08 | 0.7 | 0 | 0 | 1.61 | 0 | 0.78 | 0.02 |
| Msr1 | NM_001113326.1 | NM_001113326.1:555 | 12.37 | 12.45 | 12.17 | 11.86 | 12.07 | 12.12 | 11.88 | 12.34 | 12.6 | 11.26 | 11.08 | 11.74 |
| Mst1r | NM_009074.1 | NM_009074.1:3135 | 0.39 | 0.08 | 2.13 | 0.11 | 0.08 | 0.7 | 0 | 0 | 1.61 | 0 | 0.78 | 0.02 |
| Muc1 | NM_013605.1 | NM_013605.1:1445 | 0.39 | 1.99 | 2.13 | 0.11 | 0.08 | 0.7 | 0 | 0 | 1.61 | 0 | 0.78 | 0.02 |
| Mx1 | NM_010846.1 | NM_010846.1:1078 | 3.85 | 3.36 | 5.18 | 4.03 | 4.34 | 4.52 | 2.06 | 4.02 | 5.05 | 0 | 2.03 | 0.02 |
| Mx2 | NM_013606.1 | NM_013606.1:2095 | 6.37 | 8.96 | 8.93 | 6.73 | 8.21 | 6.06 | 7.72 | 8.83 | 9.14 | 6.69 | 0.78 | 6.71 |
| Myc | NM_010849.4 | NM_010849.4:630 | 6.45 | 0.08 | 4.22 | 4.29 | 5.41 | 7.35 | 3.06 | 5.76 | 5.05 | 5.31 | 0.78 | 0.02 |
| Myd88 | NM_010851.2 | NM_010851.2:1595 | 8.3 | 8.72 | 8.88 | 8.67 | 8.19 | 8.51 | 8.06 | 8.96 | 9.34 | 8.83 | 9.68 | 10.25 |
| Ncam1 | NM_001113204.1 | NM_001113204.1:740 | 0.39 | 0.08 | 5.34 | 4.44 | 0.08 | 2.74 | 1.9 | 3.71 | 3.89 | 5.77 | 7.23 | 0.02 |
| Ncf4 | NM_008677.2 | NM_008677.2:741 | 6.76 | 7.49 | 6.24 | 5.67 | 6.54 | 6.16 | 6.27 | 6.76 | 7.16 | 7.88 | 7.6 | 8.98 |
| Ncr1 | NM_010746.3 | NM_010746.3:391 | 0.39 | 6 | 2.13 | 0.11 | 3.1 | 3.06 | 5.06 | 5.9 | 1.61 | 5.54 | 5.05 | 5.89 |
| Nefl | NM_010910.1 | NM_010910.1:1303 | 4.85 | 3.51 | 2.13 | 4.37 | 0.08 | 0.7 | 1.32 | 2.12 | 1.61 | 3.01 | 0.78 | 0.02 |
| Nfatc1 | NM_016791.4 | NM_016791.4:1570 | 7.21 | 4.06 | 2.13 | 5.02 | 5.33 | 2.74 | 5.02 | 4.64 | 5.68 | 6.09 | 7.71 | 4.47 |
| Nfatc2 | NM_001037177.1 | NM_001037177.1:1559 | 5.78 | 5.32 | 2.13 | 5.61 | 6.11 | 4.8 | 5.86 | 4.68 | 5.05 | 3.89 | 4.01 | 6.26 |
| Nfatc3 | NM_010901.2 | NM_010901.2:2260 | 7.73 | 7.95 | 8.37 | 7.85 | 7.79 | 8 | 7.86 | 7.98 | 7.16 | 9.18 | 5.13 | 8.81 |
| Nfatc4 | NM_023699.3 | NM_023699.3:2205 | 0.39 | 0.08 | 2.13 | 0.11 | 0.08 | 0.7 | 0.32 | 0 | 1.61 | 3.43 | 0.78 | 5.22 |
| Nfkb1 | NM_008689.2 | NM_008689.2:2125 | 9.09 | 10.3 | 8.21 | 9.11 | 9.08 | 8.93 | 7.35 | 9.56 | 9.46 | 7.45 | 7.31 | 10.25 |
| Nfkb2 | NM_019408.2 | NM_019408.2:1150 | 8.05 | 8.87 | 7.84 | 7.11 | 8.77 | 7.87 | 8.95 | 7.63 | 8.89 | 9.78 | 8.47 | 10.39 |
| Nfkbia | NM_010907.2 | NM_010907.2:646 | 9.93 | 9.85 | 9.77 | 8.81 | 9.24 | 9.03 | 8.42 | 9.93 | 9.95 | 10.56 | 11.21 | 12.24 |
| Nlrc5 | NM_001033207.3 | NM_001033207.3:6665 | 10.85 | 11.47 | 11.32 | 11.47 | 10.17 | 11.21 | 9.23 | 10.36 | 10.79 | 10.31 | 10.42 | 11.21 |
| Nlrp3 | NM_145827.3 | NM_145827.3:508 | 9.02 | 9.9 | 9.15 | 7.82 | 8.66 | 8.34 | 7.36 | 8.56 | 8.57 | 10.98 | 12.05 | 12.88 |
| Nod1 | NM_172729.2 | NM_172729.2:1446 | 0.39 | 0.08 | 2.13 | 0.11 | 0.08 | 3.06 | 0.73 | 0 | 3.56 | 3.82 | 0.78 | 2.8 |
| Nod2 | NM_145857.2 | NM_145857.2:1086 | 3.39 | 5.8 | 5.49 | 0.11 | 4.84 | 2.34 | 3.82 | 4.02 | 5.05 | 5.45 | 4.52 | 4.71 |
| Nos2 | NM_010927.3 | NM_010927.3:3715 | 6.69 | 0.08 | 8.96 | 4.29 | 0.08 | 7.06 | 0 | 5.29 | 5.5 | 3.89 | 0.78 | 5.4 |
| Notch1 | NM_008714.2 | NM_008714.2:1425 | 7.53 | 7.46 | 7.19 | 6.65 | 6.62 | 6.91 | 4.86 | 5.98 | 6.46 | 6.59 | 7.59 | 7.58 |
| Nrp1 | NM_008737.2 | NM_008737.2:1190 | 9.21 | 9.39 | 8.91 | 9.03 | 9.21 | 9.4 | 9.14 | 8.99 | 8.98 | 7.63 | 8.21 | 7.58 |
| Nt5e | NM_011851.3 | NM_011851.3:1600 | 7.1 | 8.17 | 7.87 | 5.82 | 7.07 | 5.95 | 6.64 | 7.76 | 8.06 | 9 | 7.64 | 9.34 |
| Nup107 | NM_134010.2 | NM_134010.2:200 | 6.58 | 6.22 | 3.3 | 4.87 | 5.84 | 5.1 | 6.61 | 6.49 | 1.61 | 6.12 | 6.59 | 4.18 |
| Oas2 | NM_145227.3 | NM_145227.3:414 | 8.16 | 7.91 | 8.12 | 6.57 | 7.58 | 7.66 | 6.77 | 7.33 | 8.53 | 7.32 | 6.85 | 8.4 |
| Oas3 | NM_145226.2 | NM_145226.2:365 | 5.78 | 7.22 | 6.61 | 4.29 | 5.04 | 5.61 | 6.22 | 6.65 | 6.3 | 4.68 | 6.43 | 7.22 |
| Oasl1 | NM_145209.3 | NM_145209.3:626 | 10.12 | 9.45 | 9.46 | 7.02 | 9.44 | 9.53 | 9.54 | 8.99 | 9.61 | 7.86 | 8.81 | 9.3 |
| Osm | NM_001013365.2 | NM_001013365.2:579 | 0.39 | 4.92 | 2.13 | 0.11 | 0.08 | 0.7 | 2.19 | 0 | 1.61 | 4.89 | 0.78 | 6.01 |
| Pax5 | NM_008782.2 | NM_008782.2:90 | 0.39 | 2.83 | 2.13 | 3.83 | 2.91 | 2.74 | 1.32 | 0 | 1.61 | 0 | 4.81 | 0.02 |
| Pdcd1 | NM_008798.1 | NM_008798.1:1134 | 0.39 | 0.08 | 2.13 | 1.76 | 1.73 | 0.7 | 0 | 0 | 3.56 | 0 | 0.78 | 0.02 |

| | | | | | | | | | | | | | | |
|---|---|---|---|---|---|---|---|---|---|---|---|---|---|---|
| Pdcd1lg2 | NM_021396.2 | NM_021396.2:1870 | 8.23 | 8.13 | 9.31 | 7.64 | 6.77 | 9.14 | 7.41 | 3.87 | 7.62 | 8.61 | 9.62 | 12.2 |
| Pdgfc | NM_019971.2 | NM_019971.2:1165 | 6.32 | 6.03 | 4.99 | 6.18 | 8.13 | 7.73 | 6.81 | 8.01 | 6.77 | 6.84 | 7.26 | 6.33 |
| Pdgfrb | NM_008809.1 | NM_008809.1:1185 | 0.39 | 0.08 | 2.13 | 0.11 | 4.26 | 0.7 | 0 | 0 | 1.61 | 0 | 0.78 | 0.02 |
| Pecam1 | NM_008816.2 | NM_008816.2:1100 | 4.09 | 6.3 | 2.13 | 0.11 | 0.08 | 0.7 | 6.27 | 1.83 | 5.05 | 4.18 | 4.72 | 0.02 |
| Pik3cd | XM_003945690.1 | XM_003945690.1:4648 | 10.28 | 9.21 | 10.38 | 10.06 | 9.53 | 10.14 | 8.87 | 9.52 | 10.26 | 9.69 | 10.57 | 10.69 |
| Pik3cg | NM_020272.2 | NM_020272.2:2890 | 7.13 | 6.98 | 6.47 | 7.09 | 6.11 | 6.43 | 6.57 | 6.86 | 7.46 | 8.16 | 7.29 | 8.56 |
| Pin1 | NM_023371.3 | NM_023371.3:2480 | 0.39 | 0.08 | 2.13 | 5.25 | 3.27 | 0.7 | 4.9 | 5.48 | 1.61 | 0 | 0.78 | 0.02 |
| Pla2g1b | NM_011107.1 | NM_011107.1:360 | 5.85 | 2.6 | 2.13 | 6.38 | 1.73 | 0.7 | 1.54 | 0 | 1.61 | 3.24 | 0.78 | 5.53 |
| Pla2g6 | NM_001199023.1 | NM_001199023.1:768 | 0.39 | 3.36 | 2.13 | 0.11 | 0.08 | 0.7 | 0 | 0 | 1.61 | 2.6 | 0.78 | 5.26 |
| Plau | NM_008873.2 | NM_008873.2:1950 | 13.3 | 13.47 | 13.22 | 13.39 | 13.41 | 13.21 | 13.08 | 13.13 | 13.45 | 12.48 | 12.02 | 13.02 |
| Plaur | NM_011113.3 | NM_011113.3:1085 | 12.43 | 12.94 | 12.5 | 12.1 | 11.87 | 11.89 | 11.34 | 12.14 | 12.77 | 13.08 | 13.13 | 14.19 |
| Pmch | NM_029971.2 | NM_029971.2:215 | 0.39 | 0.08 | 2.13 | 0.11 | 0.08 | 0.7 | 0 | 0 | 1.61 | 0 | 0.78 | 0.02 |
| Pml | NM_008884.5 | NM_008884.5:616 | 5.71 | 2.33 | 2.13 | 0.11 | 0.25 | 3.53 | 6.56 | 5.6 | 1.61 | 0 | 0.78 | 0.02 |
| Pnma1 | NM_027438.3 | NM_027438.3:906 | 0.39 | 0.08 | 2.13 | 2.73 | 1.17 | 0.87 | 4.38 | 0 | 1.61 | 0 | 0.78 | 0.02 |
| Pou2af1 | NM_011136.2 | NM_011136.2:1865 | 8.37 | 7.54 | 9.67 | 7.29 | 8.81 | 9.04 | 0 | 0.25 | 1.61 | 6.77 | 6.25 | 0.02 |
| Pou2f2 | NM_001163556.1 | NM_001163556.1:52 | 8.95 | 4.23 | 4.78 | 5.07 | 7.14 | 5.75 | 8.17 | 6.55 | 7.27 | 6.12 | 0.78 | 2.57 |
| Pparg | NM_011146.1 | NM_011146.1:1060 | 6.2 | 6.1 | 5.63 | 6.67 | 6.84 | 5.71 | 5.35 | 6.96 | 6.92 | 6.24 | 0.78 | 3.81 |
| Ppbp | NM_023785.2 | NM_023785.2:225 | 6.37 | 8.13 | 6.32 | 6.65 | 2.7 | 5.91 | 8.66 | 7.81 | 7.73 | 7.04 | 5.55 | 7.28 |
| Prdm1 | NM_007548.3 | NM_007548.3:1440 | 7.32 | 7.94 | 8.21 | 8.91 | 7.58 | 7.63 | 7.21 | 7.95 | 7.85 | 8.21 | 9.06 | 7.53 |
| Prf1 | NM_011073.2 | NM_011073.2:1350 | 0.39 | 0.08 | 2.13 | 0.11 | 0.08 | 0.7 | 0 | 0 | 1.61 | 1.75 | 0.78 | 4 |
| Prg2 | NM_008920.4 | NM_008920.4:119 | 0.39 | 0.08 | 2.13 | 0.11 | 0.08 | 0.7 | 0 | 0 | 1.61 | 0 | 0.78 | 0.02 |
| Prkcd | NM_011103.2 | NM_011103.2:1265 | 10.42 | 9.72 | 9.37 | 9.53 | 9.56 | 9.66 | 9.73 | 9.39 | 10.31 | 9.63 | 8.97 | 10.81 |
| Prkce | NM_011104.2 | NM_011104.2:1510 | 3.97 | 6.05 | 5.86 | 3.59 | 4.61 | 0.7 | 0.32 | 0 | 1.61 | 7.46 | 5.92 | 7.19 |
| Psen1 | NM_008943.2 | NM_008943.2:2770 | 10.03 | 8.6 | 9.62 | 9.62 | 9.22 | 9.16 | 8.4 | 8.79 | 9.37 | 9.02 | 9.07 | 9.33 |
| Psen2 | NM_001128605.1 | NM_001128605.1:560 | 0.39 | 0.08 | 2.13 | 1.76 | 0.08 | 0.7 | 0 | 0 | 1.61 | 0 | 0.78 | 0.02 |
| Psma2 | NM_008944.2 | NM_008944.2:136 | 9.4 | 9.41 | 9.45 | 9.57 | 9.01 | 9 | 8.98 | 9.06 | 9.33 | 9.05 | 8.78 | 8.74 |
| Psmb10 | NM_013640.3 | NM_013640.3:401 | 0.39 | 0.08 | 2.13 | 0.11 | 2.13 | 0.7 | 1.73 | 5 | 1.61 | 1.01 | 0.78 | 0.02 |
| Psmb7 | NM_011187.1 | NM_011187.1:184 | 8.88 | 9.54 | 8.58 | 8.94 | 9.24 | 9.46 | 9.69 | 9.45 | 9.29 | 9.26 | 8.85 | 9.54 |
| Psmb8 | NM_010724.2 | NM_010724.2:362 | 8.17 | 7.85 | 7.19 | 0.28 | 7.93 | 7.63 | 8.05 | 6.88 | 8.4 | 7.38 | 6.54 | 8.61 |
| Psmb9 | NM_013585.2 | NM_013585.2:540 | 12.29 | 12.55 | 12.01 | 10.92 | 12.06 | 11.73 | 11.66 | 11.62 | 12.29 | 11.26 | 11.46 | 11.5 |
| Psmd7 | NM_010817.2 | NM_010817.2:1424 | 12.04 | 12.43 | 11.67 | 12.57 | 11.52 | 12 | 11.83 | 11.87 | 11.83 | 12.61 | 11.87 | 12.14 |
| Ptgdr2 | NM_009962.2 | NM_009962.2:270 | 0.39 | 0.08 | 2.13 | 0.28 | 0.08 | 0.7 | 0 | 0 | 1.61 | 0 | 0.78 | 0.02 |
| Ptgs2 | NM_011198.3 | NM_011198.3:675 | 11.02 | 11.26 | 11.2 | 8.89 | 9.78 | 10.37 | 9.57 | 10.49 | 11.54 | 12.61 | 12.93 | 13.98 |
| Ptprc | NM_011210.3 | NM_011210.3:2320 | 10.89 | 11.13 | 11.03 | 10.38 | 10.99 | 10.61 | 9.98 | 10.71 | 11.34 | 11.49 | 11.78 | 12.67 |
| Pvr | NM_027514.2 | NM_027514.2:962 | 9.52 | 9.91 | 9.01 | 8.06 | 8.64 | 8.6 | 8.3 | 9.54 | 9.76 | 6.93 | 7.58 | 7.52 |
| Pvrl2 | NM_001159724.1 | NM_001159724.1:984 | 0.39 | 0.08 | 2.13 | 0.11 | 0.08 | 0.7 | 0 | 0 | 1.61 | 2.6 | 0.78 | 2.3 |
| Pycard | NM_023258.4 | NM_023258.4:1654 | 3.85 | 4.77 | 2.13 | 2.47 | 3.56 | 4.71 | 5.49 | 7.45 | 1.61 | 2.89 | 4.15 | 5.66 |
| Raet1 | NM_009018.1 | NM_009018.1:548 | 5.64 | 7.49 | 5.86 | 5.73 | 7.51 | 6.83 | 5.45 | 7.47 | 7.16 | 6.44 | 7.38 | 6.18 |
| Rag1 | NM_009019.2 | NM_009019.2:1945 | 0.39 | 0.08 | 2.13 | 0.11 | 0.08 | 0.7 | 2.06 | 0 | 1.61 | 0 | 0.78 | 0.02 |
| Rel | NM_009044.2 | NM_009044.2:1290 | 13.24 | 13.44 | 12.75 | 12.69 | 12.62 | 12.87 | 12.6 | 12.98 | 13.03 | 13.17 | 13.08 | 14.33 |
| Rela | NM_009045.4 | NM_009045.4:645 | 7.39 | 8.41 | 8.16 | 7.85 | 7.78 | 7.14 | 6.52 | 7.35 | 8.66 | 8.07 | 8.9 | 9.06 |
| Relb | NM_009046.2 | NM_009046.2:2013 | 10.07 | 11.12 | 10.33 | 9.73 | 10.13 | 10.03 | 9.43 | 10.7 | 9.98 | 10.94 | 11.34 | 11.96 |
| Reps1 | NM_001111065.1 | NM_001111065.1:294 | 8.34 | 7.33 | 8.63 | 5.36 | 7.06 | 8.2 | 8.95 | 7.81 | 7.96 | 7.56 | 9.44 | 6.69 |
| Ripk2 | NM_138952.3 | NM_138952.3:830 | 7.38 | 7.99 | 5.75 | 6.64 | 5.92 | 6.29 | 5.69 | 6.73 | 8.1 | 5.29 | 4.52 | 6.7 |
| Rora | NM_013646.1 | NM_013646.1:1045 | 4.56 | 6.12 | 2.13 | 5.44 | 5.55 | 4.41 | 5.07 | 3.87 | 3.89 | 5.07 | 0.78 | 0.02 |
| Rorc | NM_011281.2 | NM_011281.2:648 | 0.39 | 0.08 | 2.13 | 0.11 | 0.08 | 0.7 | 0.32 | 5.14 | 1.61 | 0 | 0.78 | 0.02 |
| Rps6 | NM_009096.3 | NM_009096.3:1195 | 9.93 | 9.96 | 9.25 | 10.25 | 9.67 | 9.51 | 10.19 | 9.64 | 10.02 | 10.52 | 10.04 | 10.38 |
| Rrad | NM_019662.2 | NM_019662.2:1439 | 6.43 | 8.73 | 6.4 | 7.85 | 8.16 | 8.55 | 6.35 | 3.31 | 7.87 | 5.31 | 4.29 | 6.99 |
| Rsad2 | NM_021384.4 | NM_021384.4:510 | 9.4 | 10.86 | 10.04 | 8.51 | 8.86 | 8.78 | 8.09 | 9.81 | 9.54 | 8.42 | 10.34 | 10.07 |
| Runx1 | NM_001111021.1 | NM_001111021.1:3055 | 9.16 | 10.05 | 8.92 | 9.2 | 8.9 | 9.04 | 8.35 | 9.21 | 9.2 | 6.4 | 8.5 | 9.33 |
| Runx3 | NM_019732.2 | NM_019732.2:803 | 9.53 | 9.63 | 9.65 | 7.96 | 9.61 | 9.19 | 9.18 | 9.94 | 10.32 | 8.09 | 9.19 | 7.63 |
| S100a8 | NM_013650.2 | NM_013650.2:227 | 10.59 | 10.83 | 10.28 | 9.06 | 7.19 | 8.65 | 8.56 | 7.95 | 5.68 | 14.22 | 14.25 | 15.2 |
| S100b | NM_009115.3 | NM_009115.3:1090 | 0.39 | 0.08 | 2.13 | 1.2 | 0.08 | 0.7 | 0.73 | 0 | 1.61 | 0 | 0.78 | 0.02 |
| Saa1 | NM_009117.3 | NM_009117.3:351 | 0.39 | 0.08 | 4.22 | 3.3 | 3.42 | 0.7 | 0 | 0 | 3.13 | 6.09 | 3.84 | 3.16 |
| Sbno2 | NM_183426.1 | NM_183426.1:2656 | 7.02 | 4.87 | 2.13 | 4.44 | 5.87 | 5.35 | 5.19 | 5.6 | 6.64 | 7.37 | 7.98 | 8.26 |
| Sele | NM_011345.2 | NM_011345.2:2575 | 0.39 | 5.53 | 2.13 | 3.13 | 1.73 | 0.7 | 5.13 | 5.96 | 1.61 | 7.58 | 4.41 | 8.23 |
| Sell | NM_001164059.1 | NM_001164059.1:664 | 9.84 | 9.56 | 9.63 | 6.09 | 7.64 | 7.19 | 8.1 | 8.54 | 8.97 | 13.03 | 13.25 | 13.93 |
| Selplg | NM_009151.3 | NM_009151.3:1712 | 12.64 | 12.81 | 13.07 | 13.35 | 12.51 | 13.17 | 12.04 | 12.21 | 12.57 | 13.39 | 14.49 | 14.68 |
| Serpinb2 | NM_001174170.1 | NM_001174170.1:715 | 0.39 | 0.08 | 2.13 | 0.11 | 0.08 | 0.7 | 0 | 3.19 | 6.12 | 0 | 2.53 | 2.8 |
| Serping1 | NM_009776.3 | NM_009776.3:1480 | 6.6 | 0.08 | 2.13 | 6.38 | 0.08 | 0.7 | 5.54 | 0.25 | 1.61 | 8.04 | 7.23 | 0.02 |
| Sh2b2 | NM_018825.3 | NM_018825.3:1074 | 0.39 | 0.08 | 2.13 | 0.28 | 0.08 | 0.7 | 0 | 0.25 | 1.61 | 0 | 4.01 | 3.32 |
| Sh2d1a | NM_011364.3 | NM_011364.3:250 | 0.39 | 1.54 | 2.13 | 0.11 | 1.73 | 0.7 | 0 | 0 | 1.61 | 0 | 0.78 | 0.92 |
| Sh2d1b1 | NM_012009.4 | NM_012009.4:973 | 5.94 | 9.26 | 7.84 | 10.27 | 9.12 | 9.97 | 7.89 | 6.77 | 9.12 | 6.42 | 4.81 | 3.7 |
| Sigirr | NM_023059.3 | NM_023059.3:800 | 0.39 | 0.08 | 2.13 | 0.11 | 0.08 | 0.7 | 0 | 0 | 1.61 | 0 | 0.78 | 0.02 |
| Siglec1 | NM_011426.3 | NM_011426.3:4550 | 6.73 | 7.41 | 6.9 | 7.39 | 6.68 | 5.99 | 6.16 | 6.48 | 6.6 | 7.26 | 7.59 | 5.01 |
| Slamf1 | NM_013730.4 | NM_013730.4:1770 | 0.39 | 2.33 | 2.13 | 0.11 | 0.08 | 0.7 | 0 | 0.25 | 1.61 | 0 | 0.78 | 2.57 |
| Slamf6 | NM_030710.2 | NM_030710.2:805 | 3.56 | 4.15 | 2.13 | 4.7 | 5.26 | 3.31 | 3.49 | 4.59 | 1.61 | 0 | 0.78 | 0.02 |
| Slamf7 | NM_144539.5 | NM_144539.5:750 | 8.08 | 7.53 | 9.33 | 7.18 | 7.17 | 7.42 | 6.82 | 7.11 | 9.3 | 4.01 | 7.8 | 0.02 |
| Slc11a1 | NM_013612.2 | NM_013612.2:945 | 10.05 | 9.5 | 10.01 | 8.35 | 9.97 | 10.23 | 9.07 | 9.8 | 10.76 | 11.35 | 11.1 | 11.51 |
| Slc7a11 | NM_011990.2 | NM_011990.2:1700 | 6.34 | 6.91 | 6.32 | 6.26 | 5.04 | 5.35 | 6.39 | 7.02 | 5.92 | 10.12 | 9.79 | 11.05 |
| Smad2 | NM_010754.4 | NM_010754.4:880 | 8.79 | 8.23 | 8.46 | 7.64 | 8.19 | 7.79 | 8.38 | 7.39 | 8.78 | 6.24 | 8.29 | 7.28 |

| | | | | | | | | | | | | | | |
|---|---|---|---|---|---|---|---|---|---|---|---|---|---|---|
| Smad3 | NM_016769.3 | NM_016769.3:1845 | 5.03 | 6.12 | 2.13 | 5.29 | 5.55 | 5.29 | 5.02 | 7.57 | 6.12 | 7.61 | 8.06 | 7.24 |
| Smad4 | NM_008540.2 | NM_008540.2:2885 | 6.76 | 5.69 | 6.4 | 7.44 | 7.67 | 7.32 | 6.31 | 6.61 | 7.58 | 7.72 | 8.81 | 8.5 |
| Smn1 | NM_011420.2 | NM_011420.2:390 | 8.58 | 8.59 | 9.14 | 6.67 | 8.68 | 8.12 | 9.1 | 9.47 | 9.39 | 8.53 | 6.57 | 9.62 |
| Smpd3 | NM_021491.3 | NM_021491.3:1578 | 0.39 | 0.08 | 2.13 | 0.11 | 0.08 | 0.7 | 0 | 0 | 1.61 | 0 | 0.78 | 0.02 |
| Snai1 | NM_011427.2 | NM_011427.2:910 | 0.39 | 0.08 | 2.13 | 0.11 | 0.08 | 0.7 | 0 | 0 | 1.61 | 5.95 | 6.43 | 7.77 |
| Socs1 | NM_009896.2 | NM_009896.2:1020 | 7.47 | 8.29 | 7.35 | 5.84 | 5.76 | 5.75 | 6.99 | 5.96 | 8.83 | 7.4 | 9.91 | 9.26 |
| Socs3 | NM_007707.2 | NM_007707.2:585 | 6.41 | 7.53 | 7.1 | 6.36 | 6.54 | 5.95 | 5.77 | 7.24 | 8.13 | 8.84 | 7.83 | 9.75 |
| Spink5 | NM_001081180.1 | NM_001081180.1:2863 | 0.39 | 0.08 | 2.13 | 0.11 | 0.08 | 0.7 | 0 | 0.25 | 1.61 | 5.54 | 4.72 | 4.71 |
| Spn | NM_001037810.1 | NM_001037810.1:726 | 3.2 | 0.08 | 3.3 | 2.73 | 0.08 | 1.79 | 1.73 | 0 | 2.51 | 0 | 0.78 | 0.02 |
| Spp1 | NM_009263.3 | NM_009263.3:420 | 11.71 | 12.43 | 12.4 | 11.97 | 11.88 | 11.77 | 11.28 | 12.24 | 12.84 | 11.29 | 11.67 | 12 |
| St6gal1 | NM_001252506.1 | NM_001252506.1:1564 | 10.26 | 9.93 | 10.33 | 9.52 | 10.57 | 9.56 | 9.82 | 9.82 | 10.99 | 9.86 | 8.5 | 8.78 |
| Stat1 | NM_009283.3 | NM_009283.3:1590 | 8.64 | 9.21 | 10.04 | 7.87 | 8.84 | 9.04 | 6.7 | 8.71 | 9.64 | 8.22 | 9.31 | 10.09 |
| Stat2 | NM_019963.1 | NM_019963.1:362 | 8.05 | 6.8 | 7.67 | 7.01 | 7.27 | 6.87 | 6.47 | 5.92 | 8.33 | 6.82 | 7.75 | 6.81 |
| Stat3 | NM_213659.2 | NM_213659.2:2130 | 6.64 | 6.3 | 6.54 | 5.29 | 6.23 | 5.56 | 6.77 | 7.26 | 6.64 | 8.57 | 6.69 | 9.05 |
| Stat4 | NM_011487.4 | NM_011487.4:1816 | 7.45 | 6.66 | 7.87 | 6.07 | 6.56 | 7.9 | 5.53 | 6.85 | 5.4 | 6.06 | 7.32 | 7.76 |
| Stat5b | NM_011489.3 | NM_011489.3:4855 | 9.46 | 10.35 | 9.36 | 9.37 | 9.13 | 9.28 | 8.72 | 9.24 | 9.63 | 10.33 | 10.9 | 10.72 |
| Stat6 | NM_009284.2 | NM_009284.2:3465 | 11.63 | 11.81 | 11.13 | 11.37 | 11.02 | 11.03 | 10.67 | 11.09 | 11.55 | 11.76 | 12.15 | 12.66 |
| Syk | NM_001198977.1 | NM_001198977.1:2064 | 10.58 | 8.24 | 9.46 | 8.97 | 8.76 | 9.17 | 8.16 | 9.25 | 10.26 | 10.63 | 11.15 | 11.45 |
| Syt17 | NM_138649.1 | NM_138649.1:1390 | 0.39 | 0.08 | 2.13 | 0.11 | 0.08 | 0.7 | 0 | 0 | 1.61 | 0 | 0.78 | 0.02 |
| Tab1 | NM_025609.2 | NM_025609.2:570 | 0.39 | 4.45 | 2.13 | 0.11 | 0.08 | 0.7 | 3.86 | 0 | 1.61 | 3.82 | 0.78 | 0.02 |
| Tal1 | NM_011527.2 | NM_011527.2:2490 | 0.39 | 0.08 | 2.13 | 0.11 | 0.08 | 0.7 | 0 | 0 | 1.61 | 0 | 0.78 | 0.02 |
| Tank | NM_011529.1 | NM_011529.1:491 | 10.5 | 10.3 | 10.38 | 9.39 | 9.64 | 9.73 | 10.12 | 10.86 | 10.7 | 9.83 | 9.76 | 10.74 |
| Tap1 | NM_001161730.1 | NM_001161730.1:856 | 6.17 | 6.2 | 6.47 | 0.11 | 6.35 | 6.99 | 6.6 | 5.63 | 1.61 | 5.87 | 1.24 | 6.3 |
| Tap2 | NM_011530.2 | NM_011530.2:495 | 5.3 | 0.08 | 2.13 | 0.11 | 0.25 | 2.34 | 1.73 | 5.48 | 1.61 | 5.43 | 0.78 | 0.02 |
| Tapbp | NM_009318.2 | NM_009318.2:2195 | 12.26 | 12.54 | 12.27 | 12.5 | 11.74 | 12.09 | 11.75 | 12.06 | 12.75 | 11.88 | 12.33 | 12.95 |
| Tbk1 | NM_019786.4 | NM_019786.4:440 | 6.2 | 6.26 | 2.13 | 5.25 | 3.68 | 5.61 | 5.83 | 4.68 | 5.29 | 4.01 | 0.78 | 6.52 |
| Tbx21 | NM_019507.1 | NM_019507.1:625 | 0.39 | 0.08 | 2.13 | 0.11 | 0.08 | 0.7 | 0 | 0 | 1.61 | 0 | 0.78 | 0.02 |
| Tcf7 | NM_009331.3 | NM_009331.3:1810 | 5.25 | 3.36 | 4.22 | 5.64 | 6.08 | 5.91 | 4.26 | 2.91 | 1.61 | 5.95 | 1.24 | 4.53 |
| Tdo2 | NM_019911.2 | NM_019911.2:495 | 0.39 | 0.89 | 2.13 | 0.11 | 3.1 | 1.79 | 1.32 | 1.83 | 3.56 | 0 | 5.55 | 0.02 |
| Tek | NM_013690.2 | NM_013690.2:3580 | 0.39 | 0.08 | 2.13 | 0.11 | 0.08 | 0.7 | 0 | 0 | 1.61 | 0 | 0.78 | 0.02 |
| Tfe3 | NM_172472.3 | NM_172472.3:2715 | 9.05 | 9.51 | 8.75 | 8.99 | 9.37 | 8.34 | 8.3 | 8.62 | 9.33 | 9.36 | 8.81 | 8.96 |
| Tfeb | NM_001161723.1 | NM_001161723.1:534 | 0.39 | 0.08 | 2.13 | 0.11 | 0.08 | 0.7 | 0 | 0 | 1.61 | 3.01 | 0.78 | 4.82 |
| Tfrc | NM_011638.3 | NM_011638.3:1930 | 9.73 | 8.07 | 9.27 | 9.1 | 9.14 | 9.31 | 9.83 | 9.07 | 8.93 | 10.3 | 11 | 10.4 |
| Tgfb1 | NM_011577.1 | NM_011577.1:1470 | 9.5 | 9.84 | 8.48 | 8.66 | 9 | 9.07 | 9.07 | 10.12 | 9.39 | 8.76 | 8.07 | 8.6 |
| Tgfb2 | NM_009367.1 | NM_009367.1:1685 | 0.39 | 0.08 | 2.13 | 4.03 | 0.08 | 0.7 | 5.78 | 0 | 1.61 | 6.7 | 0.78 | 0.02 |
| Tgfb3 | NM_009368.2 | NM_009368.2:2410 | 8.59 | 9.09 | 8.02 | 7.79 | 8.84 | 8.76 | 8.72 | 8.59 | 8.77 | 8.47 | 7.64 | 7.67 |
| Tgfbr1 | NM_009370.2 | NM_009370.2:4425 | 10.59 | 10.8 | 10.93 | 9.21 | 11.19 | 10.63 | 10.06 | 10.73 | 11.2 | 9.7 | 9.98 | 10.96 |
| Tgfbr2 | NM_009371.2 | NM_009371.2:475 | 9.25 | 9.71 | 9.01 | 9.15 | 9.23 | 9.27 | 8.67 | 9.34 | 9.6 | 8.2 | 8.5 | 7.48 |
| Thbd | NM_009378.3 | NM_009378.3:2845 | 10.8 | 10.99 | 10.69 | 9.83 | 10.21 | 10.49 | 10.25 | 10.94 | 11.06 | 9.78 | 10.23 | 10.44 |
| Thbs1 | NM_011580.3 | NM_011580.3:702 | 11.87 | 12.38 | 12.04 | 9.31 | 9.52 | 10.34 | 10.86 | 11.86 | 12.52 | 10.49 | 11.65 | 11.93 |
| Thy1 | NM_009382.3 | NM_009382.3:425 | 0.39 | 0.08 | 2.13 | 0.11 | 0.08 | 0.7 | 0 | 0 | 1.61 | 0 | 0.78 | 0.02 |
| Ticam1 | NM_174989.4 | NM_174989.4:2159 | 0.39 | 0.08 | 2.13 | 0.11 | 0.08 | 0.7 | 2.19 | 6.06 | 1.61 | 3.52 | 0.78 | 0.02 |
| Ticam2 | NM_173394.2 | NM_173394.2:1243 | 8.62 | 6.66 | 8.92 | 8.14 | 7.76 | 7.39 | 6.24 | 7.02 | 8.16 | 8.24 | 6.08 | 4.53 |
| Tie1 | NM_011587.2 | NM_011587.2:2715 | 0.39 | 0.08 | 2.13 | 0.11 | 0.08 | 0.7 | 0 | 0 | 1.61 | 2.43 | 0.78 | 0.02 |
| Tigit | NM_001146325.1 | NM_001146325.1:730 | 1.39 | 6.86 | 2.13 | 0.28 | 0.08 | 0.7 | 4.09 | 4.59 | 1.61 | 1.43 | 1.24 | 0.02 |
| Timd4 | NM_178759.4 | NM_178759.4:236 | 10.01 | 8.64 | 9.73 | 9.61 | 9.65 | 10.09 | 8.67 | 8.25 | 8.46 | 8.21 | 8.8 | 5.22 |
| Tirap | NM_001177846.1 | NM_001177846.1:254 | 6.25 | 6.55 | 2.13 | 5.29 | 2.7 | 0.7 | 1.73 | 2.57 | 6.46 | 6.87 | 6.93 | 7.15 |
| Tlr1 | NM_030682.1 | NM_030682.1:805 | 9.88 | 10.12 | 9.74 | 10.35 | 9.56 | 9.87 | 9.67 | 9.66 | 10.26 | 8.95 | 7.35 | 9.38 |
| Tlr2 | NM_011905.2 | NM_011905.2:255 | 9.37 | 9.7 | 9.63 | 8.83 | 9.66 | 9.52 | 9.02 | 9.79 | 10.45 | 9.86 | 10.8 | 10.88 |
| Tlr3 | NM_126166.2 | NM_126166.2:1165 | 6.09 | 6.47 | 5.86 | 6.68 | 6.42 | 6.89 | 5.61 | 6.27 | 4.58 | 6.44 | 6.54 | 4.76 |
| Tlr4 | NM_021297.2 | NM_021297.2:2510 | 9.8 | 9.11 | 9.73 | 8.75 | 9.37 | 9 | 7.95 | 8.82 | 10 | 9.96 | 10.49 | 10.34 |
| Tlr5 | NM_016928.2 | NM_016928.2:560 | 0.39 | 0.08 | 2.13 | 0.11 | 2.91 | 0.7 | 5.69 | 0 | 1.61 | 7.98 | 0.78 | 0.92 |
| Tlr6 | NM_011604.3 | NM_011604.3:475 | 6.83 | 7.64 | 6.4 | 5.16 | 6.77 | 6.7 | 6.23 | 7.44 | 6.12 | 8.83 | 8 | 9.56 |
| Tlr7 | NM_133211.3 | NM_133211.3:3210 | 9.7 | 9.12 | 9.84 | 9.74 | 9.45 | 9.7 | 8.87 | 9.66 | 10.01 | 7.01 | 6.97 | 7.35 |
| Tlr8 | NM_133212.2 | NM_133212.2:110 | 10.21 | 10.65 | 9.67 | 10.04 | 9.68 | 10.04 | 9.49 | 10.18 | 9.73 | 8.85 | 9.07 | 9.73 |
| Tlr9 | NM_031178.2 | NM_031178.2:1801 | 7.65 | 6.72 | 7.43 | 6.38 | 6.08 | 4.88 | 4.59 | 6.51 | 6.46 | 1.43 | 4.52 | 1.97 |
| Tmed1 | NM_010744.3 | NM_010744.3:104 | 2.39 | 0.08 | 2.13 | 0.11 | 0.08 | 0.7 | 2.06 | 2.12 | 1.61 | 2.89 | 4.01 | 4.18 |
| Tmem173 | NM_028261.1 | NM_028261.1:1792 | 6.62 | 7.04 | 4.22 | 7.06 | 5.67 | 5.46 | 6.13 | 6.13 | 4.16 | 6.55 | 0.78 | 5.37 |
| Tnf | NM_013693.1 | NM_013693.1:1135 | 10.26 | 11.13 | 9.71 | 9.81 | 9.93 | 9.74 | 9.9 | 10.74 | 10.19 | 10.72 | 11.57 | 12.45 |
| Tnfaip3 | NM_009397.2 | NM_009397.2:232 | 10.52 | 11.04 | 10.42 | 9.99 | 10.18 | 10.5 | 9.4 | 10.64 | 11.03 | 11.74 | 12.37 | 13.47 |
| Tnfrsf10b | NM_020275.3 | NM_020275.3:1625 | 6.97 | 7.03 | 2.13 | 4.76 | 3.56 | 4.04 | 6.68 | 4.09 | 6.81 | 8 | 8.07 | 7.23 |
| Tnfrsf11a | NM_009399.3 | NM_009399.3:692 | 4.97 | 0.08 | 2.13 | 3.3 | 4.41 | 3.72 | 4.82 | 4.09 | 6.24 | 4.47 | 0.78 | 4.4 |
| Tnfrsf11b | NM_008764.3 | NM_008764.3:35 | 0.39 | 0.08 | 2.13 | 0.11 | 0.08 | 0.7 | 0 | 0 | 1.61 | 0 | 0.78 | 0.02 |
| Tnfrsf12a | NM_001161746.1 | NM_001161746.1:517 | 0.39 | 6.76 | 5.34 | 4.44 | 6.58 | 5.51 | 5.41 | 5 | 5.29 | 6.55 | 5.05 | 6.55 |
| Tnfrsf13b | NM_021349.1 | NM_021349.1:340 | 3.39 | 0.08 | 4.78 | 3.71 | 4.61 | 3.72 | 1.73 | 2.12 | 4.58 | 3.6 | 2.03 | 3.59 |
| Tnfrsf13c | NM_028075.2 | NM_028075.2:1170 | 6.41 | 5.85 | 6.79 | 7.1 | 6.95 | 6.46 | 0 | 0 | 1.61 | 3.24 | 0.78 | 3.7 |
| Tnfrsf14 | NM_178931.2 | NM_178931.2:625 | 6.37 | 8.34 | 7.01 | 6.22 | 7.38 | 5.41 | 6.15 | 5.48 | 7.22 | 6.13 | 6.43 | 7.86 |
| Tnfrsf17 | NM_011608.1 | NM_011608.1:140 | 0.39 | 0.08 | 2.13 | 0.11 | 0.08 | 0.7 | 0 | 0 | 1.61 | 0 | 0.78 | 0.02 |
| Tnfrsf18 | NM_009400.2 | NM_009400.2:840 | 8.37 | 7.05 | 6.07 | 7.73 | 6.54 | 7.72 | 6.27 | 0 | 6.18 | 7.01 | 8.37 | 8.78 |
| Tnfrsf1a | NM_011609.2 | NM_011609.2:615 | 8.16 | 6.58 | 6.79 | 6.57 | 7.75 | 6.97 | 5.9 | 6.55 | 7.25 | 8.45 | 8.08 | 9.51 |
| Tnfrsf1b | NM_011610.3 | NM_011610.3:3270 | 14.01 | 14.11 | 13.92 | 13.73 | 13.57 | 13.74 | 12.98 | 13.81 | 14.33 | 14.58 | 14.92 | 15.87 |

| | | | | | | | | | | | | | | |
|---|---|---|---|---|---|---|---|---|---|---|---|---|---|---|
| Tnfrsf4 | NM_011659.2 | NM_011659.2:320 | 0.39 | 2.33 | 4.78 | 5.76 | 6.67 | 3.31 | 0 | 4.54 | 7.06 | 6.32 | 0.78 | 2.8 |
| Tnfrsf8 | NM_009401.2 | NM_009401.2:1275 | 0.39 | 3.36 | 2.13 | 3.59 | 2.44 | 0.87 | 0 | 2.75 | 4.16 | 0 | 0.78 | 0.02 |
| Tnfrsf9 | NM_001077508.1 | NM_001077508.1:1590 | 0.39 | 0.08 | 2.13 | 0.11 | 0.08 | 0.7 | 0 | 0 | 1.61 | 0 | 0.78 | 0.02 |
| Tnfsf10 | NM_009425.2 | NM_009425.2:2055 | 2.97 | 4.06 | 4.78 | 5.79 | 5.79 | 5.87 | 5.14 | 2.12 | 5.59 | 5.68 | 0.78 | 4.18 |
| Tnfsf11 | NM_011613.3 | NM_011613.3:615 | 0.39 | 0.08 | 2.13 | 0.11 | 0.08 | 0.7 | 0 | 0 | 1.61 | 5.77 | 0.78 | 0.02 |
| Tnfsf12 | NM_011614.3 | NM_011614.3:1215 | 8.98 | 10.44 | 8.21 | 7.53 | 7.64 | 8.43 | 7.91 | 7.1 | 8.59 | 6.48 | 4.15 | 6.52 |
| Tnfsf13 | NM_023517.2 | NM_023517.2:1443 | 7.48 | 7.1 | 6.73 | 8.04 | 7.4 | 7.68 | 6.9 | 8.41 | 6.64 | 6.03 | 8.06 | 6.81 |
| Tnfsf13b | NM_033622.1 | NM_033622.1:225 | 0.39 | 0.08 | 2.13 | 0.11 | 3.1 | 0.7 | 0 | 2.75 | 1.61 | 0 | 0.78 | 5.63 |
| Tnfsf14 | NM_019418.2 | NM_019418.2:1060 | 5.71 | 5.36 | 6.32 | 1.2 | 4.48 | 3.31 | 0 | 4.68 | 2.51 | 8.34 | 9.78 | 10.21 |
| Tnfsf15 | NM_177371.3 | NM_177371.3:4695 | 3.2 | 0.89 | 5.97 | 2.94 | 3.56 | 4.41 | 0 | 3.31 | 4.76 | 0 | 0.78 | 0.02 |
| Tnfsf18 | NM_183391.3 | NM_183391.3:1445 | 0.39 | 0.08 | 2.13 | 5.58 | 1.17 | 0.7 | 0 | 0 | 1.61 | 3.34 | 0.78 | 7.06 |
| Tnfsf4 | NM_009452.2 | NM_009452.2:346 | 2.97 | 5.85 | 2.46 | 5.25 | 4.18 | 4.04 | 0 | 0 | 1.61 | 0 | 0.78 | 4.87 |
| Tnfsf8 | NM_009403.2 | NM_009403.2:125 | 0.39 | 0.08 | 2.13 | 0.11 | 0.08 | 0.7 | 0 | 0 | 2.51 | 1.43 | 0.78 | 0.02 |
| Tollip | NM_023764.3 | NM_023764.3:260 | 6.78 | 6.87 | 7.57 | 5.44 | 5.17 | 5.83 | 6.65 | 8.8 | 7.56 | 7.45 | 1.24 | 7.56 |
| Tpsab1 | NM_031187.4 | NM_031187.4:663 | 4.71 | 3.96 | 5.34 | 4.7 | 3.42 | 3.89 | 2.82 | 3.79 | 4.16 | 3.24 | 0.78 | 0.02 |
| Traf2 | NM_009422.2 | NM_009422.2:1334 | 4.56 | 3.03 | 5.86 | 1.2 | 1.17 | 0.7 | 4.84 | 2.12 | 3.13 | 0 | 0.78 | 5.53 |
| Traf3 | NM_001048206.1 | NM_001048206.1:6385 | 7.38 | 7.59 | 7.35 | 8.21 | 8.12 | 6.63 | 6.65 | 8.3 | 8.16 | 7.55 | 6.67 | 8.73 |
| Traf6 | NM_009424.2 | NM_009424.2:980 | 7.34 | 7.53 | 7.01 | 7.58 | 7.44 | 5.75 | 5.41 | 6.01 | 6.12 | 5.24 | 6.72 | 6.78 |
| Trem1 | NM_021406.5 | NM_021406.5:1704 | 9.23 | 9.88 | 7.97 | 6.76 | 7.98 | 8.6 | 6.48 | 7.86 | 7.98 | 10.79 | 11.36 | 12.82 |
| Trem2 | NM_031254.2 | NM_031254.2:646 | 8.92 | 9.13 | 8.51 | 8.38 | 9.3 | 8.54 | 8.34 | 8.96 | 9.07 | 6.57 | 4.72 | 7.04 |
| Trp53 | NM_011640.1 | NM_011640.1:1835 | 9.38 | 9.75 | 8.91 | 8.93 | 9.13 | 8.69 | 9.06 | 9.56 | 8.79 | 9.2 | 9.65 | 8.88 |
| Twist1 | NM_011658.2 | NM_011658.2:994 | 0.39 | 3.86 | 2.13 | 0.11 | 0.08 | 0.7 | 0 | 0 | 1.61 | 1.75 | 0.78 | 0.02 |
| Txk | NM_001122754.1 | NM_001122754.1:835 | 0.39 | 0.08 | 2.13 | 3.83 | 0.08 | 0.7 | 2.19 | 2.12 | 1.61 | 2.01 | 0.78 | 0.02 |
| Txnip | NM_023719.1 | NM_023719.1:2340 | 11.02 | 11.01 | 10.5 | 10.74 | 10.51 | 10.63 | 10.99 | 10.89 | 10.52 | 11.77 | 11.83 | 12.7 |
| Tyk2 | NM_001205312.1 | NM_001205312.1:1532 | 5.25 | 5.32 | 2.13 | 0.11 | 0.08 | 0.7 | 0 | 5.88 | 7.27 | 4.56 | 0.78 | 0.02 |
| Ubc | NM_019639.4 | NM_019639.4:2193 | 13.47 | 13.86 | 13.1 | 13.08 | 12.72 | 13.15 | 12.83 | 13.36 | 13.17 | 14.01 | 13.84 | 14.64 |
| Ulbp1 | NM_029975.2 | NM_029975.2:595 | 0.39 | 0.08 | 2.13 | 0.11 | 0.08 | 0.7 | 2.19 | 4.85 | 3.56 | 1.43 | 0.78 | 0.02 |
| Usp18 | NM_011909.2 | NM_011909.2:1190 | 6.85 | 7.02 | 7.57 | 6.49 | 6.88 | 6.77 | 7.05 | 6.85 | 8.06 | 6.4 | 5.5 | 5.53 |
| Usp9y | NM_148943.2 | NM_148943.2:480 | 0.39 | 0.08 | 2.13 | 0.11 | 0.08 | 0.7 | 0 | 2.57 | 1.61 | 0 | 0.78 | 0.02 |
| Vcam1 | NM_011693.2 | NM_011693.2:1440 | 8.83 | 9.09 | 8.05 | 8.36 | 9.66 | 8.65 | 6.6 | 8.45 | 7.56 | 7.09 | 5.96 | 8.11 |
| Vegfa | NM_001025250.3 | NM_001025250.3:3015 | 15.01 | 15.15 | 14.8 | 14.65 | 14.15 | 14.76 | 14.08 | 14.73 | 14.89 | 14.83 | 15.25 | 15.74 |
| Vegfc | NM_009506.2 | NM_009506.2:1320 | 0.39 | 0.08 | 2.13 | 0.11 | 0.08 | 0.7 | 0 | 0 | 1.61 | 0 | 0.78 | 0.02 |
| Vhl | NM_009507.3 | NM_009507.3:820 | 3.71 | 2.6 | 4.78 | 3.71 | 2.13 | 2.34 | 0 | 5.2 | 3.89 | 0 | 0.78 | 3.7 |
| Vim | NM_011701.4 | NM_011701.4:34 | 11.2 | 12.02 | 11.23 | 9.74 | 11.43 | 11.43 | 10.55 | 11.73 | 11.8 | 11.18 | 10.54 | 10.87 |
| Vwf | NM_011708.3 | NM_011708.3:6354 | 1.97 | 5.39 | 5.18 | 4.37 | 2.44 | 3.89 | 4.64 | 1.47 | 4.16 | 6.8 | 5.96 | 6.71 |
| Xaf1 | NM_001037713.3 | NM_001037713.3:100 | 8.42 | 9.29 | 8.44 | 8.11 | 7.77 | 7.94 | 7.97 | 8.13 | 8.77 | 6.8 | 8.27 | 7.5 |
| Xbp1 | NM_013842.2 | NM_013842.2:825 | 9.05 | 9.02 | 8.8 | 8.64 | 8.92 | 9.02 | 8.66 | 8.58 | 9.55 | 9.55 | 10.08 | 9.51 |
| Xcl1 | NM_008510.1 | NM_008510.1:103 | 0.39 | 0.08 | 4.78 | 3.59 | 2.13 | 0.7 | 0 | 0 | 6.12 | 3.75 | 0.78 | 0.02 |
| Xcr1 | NM_011798.4 | NM_011798.4:350 | 2.71 | 3.03 | 2.13 | 5.92 | 6.04 | 5.56 | 0 | 0 | 1.61 | 0 | 0.78 | 0.02 |
| Ythdf2 | NM_145393.4 | NM_145393.4:750 | 8.55 | 8.76 | 8.53 | 8.62 | 8.08 | 8.03 | 8.38 | 9.43 | 8.59 | 7.07 | 9.58 | 9.09 |
| Yy1 | NM_009537.3 | NM_009537.3:760 | 11.41 | 11.23 | 10.97 | 11.26 | 10.16 | 10.71 | 10.44 | 11.22 | 11.39 | 10.88 | 11.11 | 11.38 |
| Zap70 | NM_009539.2 | NM_009539.2:1030 | 0.39 | 0.08 | 2.13 | 2.16 | 0.08 | 0.7 | 0 | 0 | 1.61 | 0 | 0.78 | 0.02 |
| Zbp1 | NM_021394.2 | NM_021394.2:473 | 10.14 | 9.92 | 10.52 | 10.47 | 9.33 | 10 | 8.9 | 9.89 | 10.44 | 7.39 | 7.8 | 8.57 |
| Zfp13 | NM_011747.2 | NM_011747.2:419 | 0.39 | 1.54 | 2.13 | 0.11 | 0.08 | 0.7 | 0 | 0.98 | 1.61 | 3.24 | 0.78 | 0.02 |
| Abcf1 | NM_013854.1 | NM_013854.1:875 | 6.64 | 7.25 | 7.1 | 6.81 | 7.11 | 6.66 | 7.68 | 6.15 | 6.99 | 7.63 | 0.78 | 7.96 |
| Alas1 | NM_020559.2 | NM_020559.2:1034 | 7.28 | 7.72 | 3.84 | 5.02 | 6.02 | 6.06 | 6.62 | 4.97 | 6.06 | 8.53 | 7.44 | 10.73 |
| Edc3 | NM_153799.3 | NM_153799.3:2570 | 3.85 | 0.08 | 2.13 | 3.3 | 1.73 | 0.87 | 2.98 | 4.5 | 1.61 | 2.43 | 0.78 | 0.92 |
| Eef1g | NM_026007.4 | NM_026007.4:338 | 8.59 | 8.13 | 7.6 | 6.26 | 7.21 | 7.73 | 8.68 | 7.15 | 8.53 | 8.64 | 7.78 | 7.51 |
| Eif2b4 | NM_001127355.1 | NM_001127355.1:1562 | 6.64 | 6.65 | 2.46 | 6.26 | 6.64 | 5.51 | 6.25 | 6.16 | 6.89 | 7.4 | 7.01 | 6.64 |
| G6pdx | NM_008062.2 | NM_008062.2:2030 | 11.09 | 11.13 | 11.36 | 11.47 | 11.41 | 11.42 | 11.26 | 11.14 | 11.95 | 11.38 | 11.82 | 11.3 |
| Gusb | NM_010368.1 | NM_010368.1:1735 | 7.81 | 8.98 | 7.6 | 9.12 | 8.53 | 8.43 | 8.51 | 8.71 | 8.19 | 7.84 | 8.72 | 9.07 |
| Hdac3 | NM_010411.2 | NM_010411.2:144 | 5.03 | 5.93 | 5.18 | 3.45 | 3.27 | 5.87 | 5.79 | 4.64 | 5.18 | 4.43 | 2.53 | 8.28 |
| Hprt | NM_013556.2 | NM_013556.2:428 | 11.66 | 11.44 | 11.22 | 11.94 | 11.36 | 11.42 | 11.36 | 11.45 | 11.39 | 11.47 | 11.48 | 11.56 |
| Nubp1 | NM_011955.2 | NM_011955.2:284 | 1.97 | 5.39 | 2.13 | 0.11 | 0.08 | 0.7 | 2.98 | 4.93 | 1.61 | 2.75 | 0.78 | 0.02 |
| Oaz1 | NM_008753.4 | NM_008753.4:809 | 13.69 | 13.72 | 13.5 | 13 | 13.34 | 13.29 | 13.29 | 13.53 | 13.93 | 13.53 | 13.7 | 14.11 |
| Polr1b | NM_009086.2 | NM_009086.2:2795 | 3.85 | 0.08 | 2.13 | 0.11 | 0.08 | 0.7 | 0 | 0 | 1.61 | 4.07 | 0.78 | 0.02 |
| Polr2a | NM_009089.2 | NM_009089.2:2220 | 7.12 | 8.33 | 6.07 | 6.36 | 6.49 | 5.91 | 6.91 | 8.01 | 7.81 | 7.88 | 8.54 | 9.22 |
| Ppia | NM_008907.1 | NM_008907.1:390 | 11.62 | 11.71 | 11.55 | 12.04 | 11.03 | 11.21 | 11.44 | 11.4 | 11.77 | 12.19 | 11.61 | 12.37 |
| Rpl19 | NM_009078.2 | NM_009078.2:92 | 9.84 | 10.08 | 10.52 | 9.42 | 9.73 | 9.56 | 9.47 | 9.01 | 10.29 | 9.99 | 9.33 | 9.8 |
| Sap130 | NM_172965.2 | NM_172965.2:2182 | 7.72 | 5.43 | 3.3 | 4.7 | 6.68 | 7.26 | 8.29 | 8.34 | 8.44 | 7.95 | 7.99 | 7.97 |
| Sdha | NM_023281.1 | NM_023281.1:250 | 1.97 | 4.97 | 2.13 | 1.76 | 4.18 | 4.8 | 4.49 | 5.6 | 4.91 | 4.89 | 5.55 | 2.57 |
| Sf3a3 | NM_029157.3 | NM_029157.3:40 | 7.31 | 6.3 | 6.85 | 7 | 7.32 | 7.36 | 7.51 | 7.12 | 5.4 | 6.97 | 6.25 | 5.26 |
| Tbp | NM_013684.3 | NM_013684.3:70 | 8.84 | 8.63 | 9.59 | 8.1 | 9.16 | 9.1 | 9.12 | 9.63 | 9.08 | 8.97 | 9.45 | 8.69 |
| Tubb5 | NM_011655.4 | NM_011655.4:2260 | 12.57 | 12.43 | 12.24 | 12.34 | 12.55 | 12.64 | 12.47 | 12.43 | 12.43 | 12.09 | 12.06 | 12.27 |
